# A long-acting prolactin to combat lactation insufficiency

**DOI:** 10.1101/2023.12.15.571886

**Authors:** Kasia Kready, Kailyn Doiron, Katherine Redfield Chan, Jeffrey Way, Quincey Justman, Camille E. Powe, Pamela Silver

## Abstract

Human infants are born to breastfeed. While 50% of lactating persons struggle to make enough milk, there are no governmentally-approved drugs to enhance lactation^1^. Here, we engineer a variant of the naturally-occurring driver of lactation, the hormone Prolactin, to increase its serum half-life and produce a viable drug candidate. Our engineered variant, Prolactin-eXtra Long-acting (Prolactin-XL), is comprised of endogenously active human prolactin fused to an engineered human IgG Fc domain designed to overcome the unique drug development challenges specific to the lactating person-infant dyad. Our Prolactin-XL has a serum half-life of 70.9h in mice, 2,625-fold longer than endogenously active prolactin alone (70.9h v. 0.027h). We demonstrate that Prolactin-XL increases milk production and restores growth of pups fed by dams with pharmacologically-ablated lactation. We show that Prolactin-XL-enhanced lactation is accompanied by reversible, lactocyte-driven changes in mammary gland morphology. This work establishes long-acting prolactins as a potentially powerful pharmacologic means to combat insufficient lactation.

## Introduction

Breastfeeding is the biological norm and standard of care for every day infant nutrition worldwide^2^. There is universal consensus amongst scientists, clinicians, major health organizations, national governments, and the infant formula industry that breastfeeding is best for most infants^1–13^. Breastfeeding is also crucial for protecting vulnerable infants when there is no electricity, no safe water, disrupted supply chains, or infant formula shortages, contaminations, or recalls^9,14–18^. However, 50% of all lactating persons struggle to make enough milk^1^.

The naturally-occurring hormone, Prolactin, is a strong driver of lactation in all mammals^19^. Human prolactin (hPRL) is a small 23 kDa protein that binds to prolactin receptor (PRLR) dimers expressed in lactocytes of the mammary gland to induce milk production^19^. During lactation, infant suckling induces secretion of prolactin from the pituitary to the blood stream, whereby it increases milk production in the mammary gland and creates a positive feedback loop. However, prolactin is short-lived, with a serum half-life of ∼40 minutes, and within several hours, prolactin is cleared from the body by the kidneys^20^.

Clinical evidence shows that twice-daily subcutaneous injections of recombinant human prolactin increases milk production in parents with insufficient milk supply^21^. Although native prolactin’s short-lived half-life makes it an unfavorable drug candidate, recent clinical successes demonstrate that a therapeutic protein’s serum half-life can be increased by fusing it to the fragment crystallizable (Fc) region of the constant domain of human immunoglobulins (IgG) ^22–26^. The resulting fusion integrates the multiple functions of the Fc domain and the pharmacological properties of the active drug protein.

Here, we engineer a long-acting, human Fc-prolactin fusion called Prolactin-XL. Prolactin-XL has a serum half-life of 70.9h in mice, 2,625-fold longer than endogenously active prolactin alone (70.9h v. 0.027h). We demonstrate that Prolactin-XL increases milk production and restores growth of pups fed by dams with pharmacologically-ablated lactation. We show that Prolactin-XL-enhanced lactation is accompanied by reversible, lactocyte-driven changes in mammary gland morphology. This work establishes engineered, long-acting prolactins as a powerful means to combat insufficient lactation and study enhancing milk supply across all mammals.

## Results

### Design and identification of a long-acting human Fc-prolactin fusion variant, Prolactin-XL

To engineer a safe and effective long-acting Fc-prolactin fusion variant, our design challenge was five-fold: (1) fuse prolactin to an Fc domain without disrupting PRLR signaling, (2) decrease Fc domain’s binding to off-target Fc receptors, (3) improve the serum half-life vs. prolactin, (4) minimize uptake by the breastfeeding infant by decreasing oral bioavailability in the infant gut, and (5) increase milk production without producing pathological effects on breast tissue. To meet this five-fold challenge, we used existing structural information from solved structures of hPRL, hPRLR, and homologous hormones and receptors in highly similar complexes (PDB 1f6f^27^, 3d48^28^, 1rw5^29^) to rationally engineer and score 29 hPRL-Fc fusion variants (Supplemental Tables S1-3) as follows:

#### Maintenance of hPRLR signaling

The 29 engineered variants vary in domain topology, linker regions, and are composed of homo-and heterodimers to maximize the likelihood of productive signaling through the hPRLR in the mammary gland^30,31,32^ (Fig. 1B-II). In addition, because glycosylation sterically occludes hPRL from binding to hPRLR^33^, all engineered variants have the de-glycosylated hPRL mutant N59D. Engineered variants were scored for their ability signal via the hPRLR or mouse PRLR (mPRLR) by measuring *in vitro* proliferation of murine Ba/F3 cells stably expressing hPRLR or mPRLR, respectively (Fig. 2C, Extended Data Fig. 2-5).

**Figure 1:**
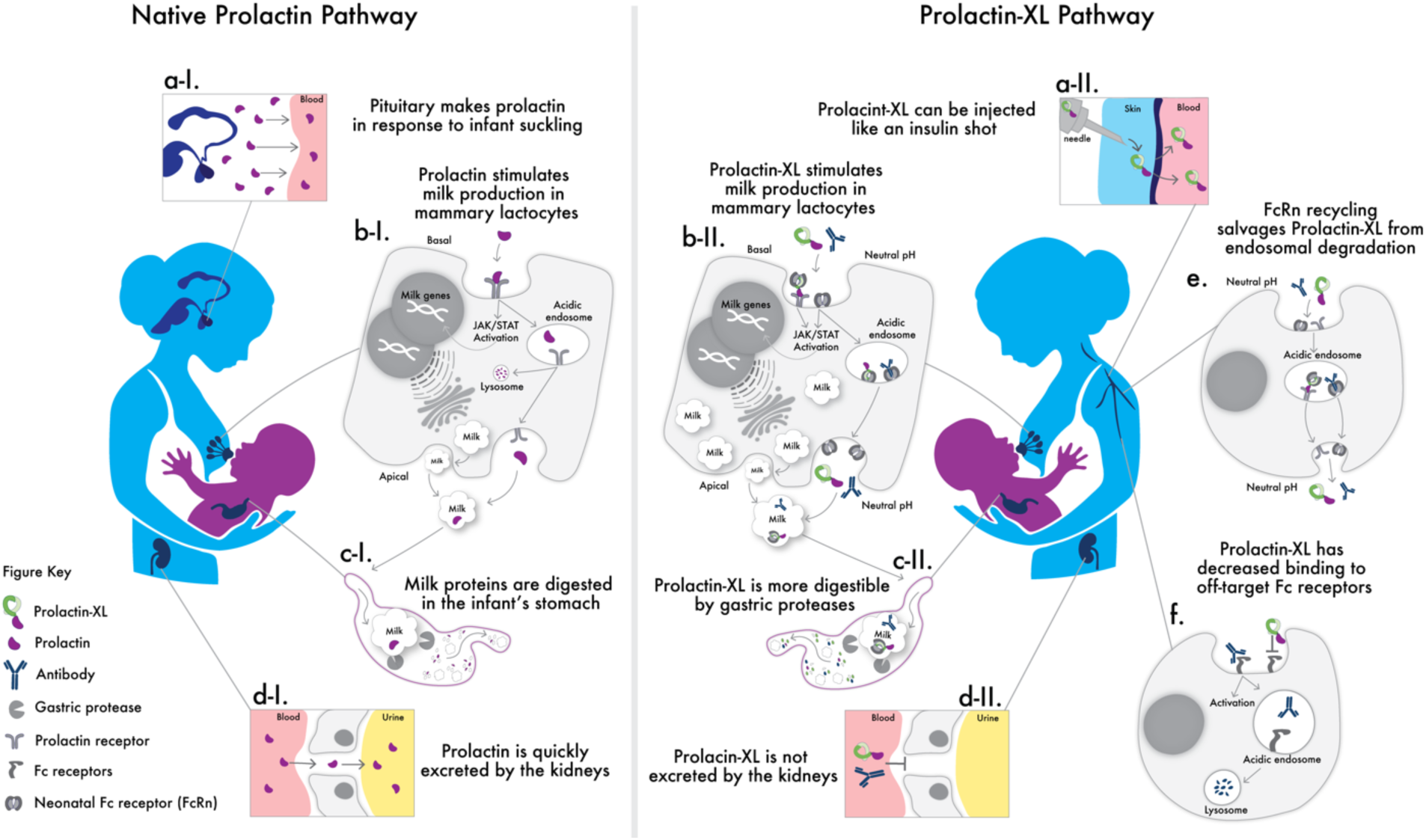
Prolactin-XL is designed to overcome the unique drug development challenges specific to the lactating person-infant dyad. Our top performing long-acting prolactin, called Prolactin-XL, is designed with tailored pharmacological properties to create a biologic that shares the milk-stimulating effects of endogenous prolactin while addressing several disadvantages that prevent recombinant prolactin from being a viable treatment. **a-i** endogenous prolactin which is secreted by the pituitary in response to infant suckling. **a-ii** Prolactin-XL is a long-acting injectable consisting of Fc, a subdomain of an IgG1 antibody, fused to the endogenously active, deglycosylated prolactin mutant N59D. Prolactin-XL is compatible with self-administered subcutaneous injections, like insulin shots. **b-i** Prolactin stimulates mammary lactocytes to produce milk. **b-ii** Likewise, Prolactin-XL maintains prolactin-receptor-signaling-induced milk production in mammary lactocytes. **c-i** Milk proteins, including endogenous Prolactin, are digested by gastric proteases in the infant’s stomach. **c-ii** Prolactin-XL is engineered to be more digestible by gastric proteases to decrease the oral bioavailability to the infant. **d-i** Prolactin is a small 23kDa protein that is quickly excreted by the kidneys. **d-ii** However, Prolactin-XL is a large 75kDa protein that is not excreted by the kidneys. **e** pH-dependent binding to the neonatal Fc receptor (FcRn), which salvages Prolactin-XL from endosomal degradation. **f** Prolactin-XL has been mutated to prevent off-target binding to Fc receptors on immune cells.

**Figure 2:**
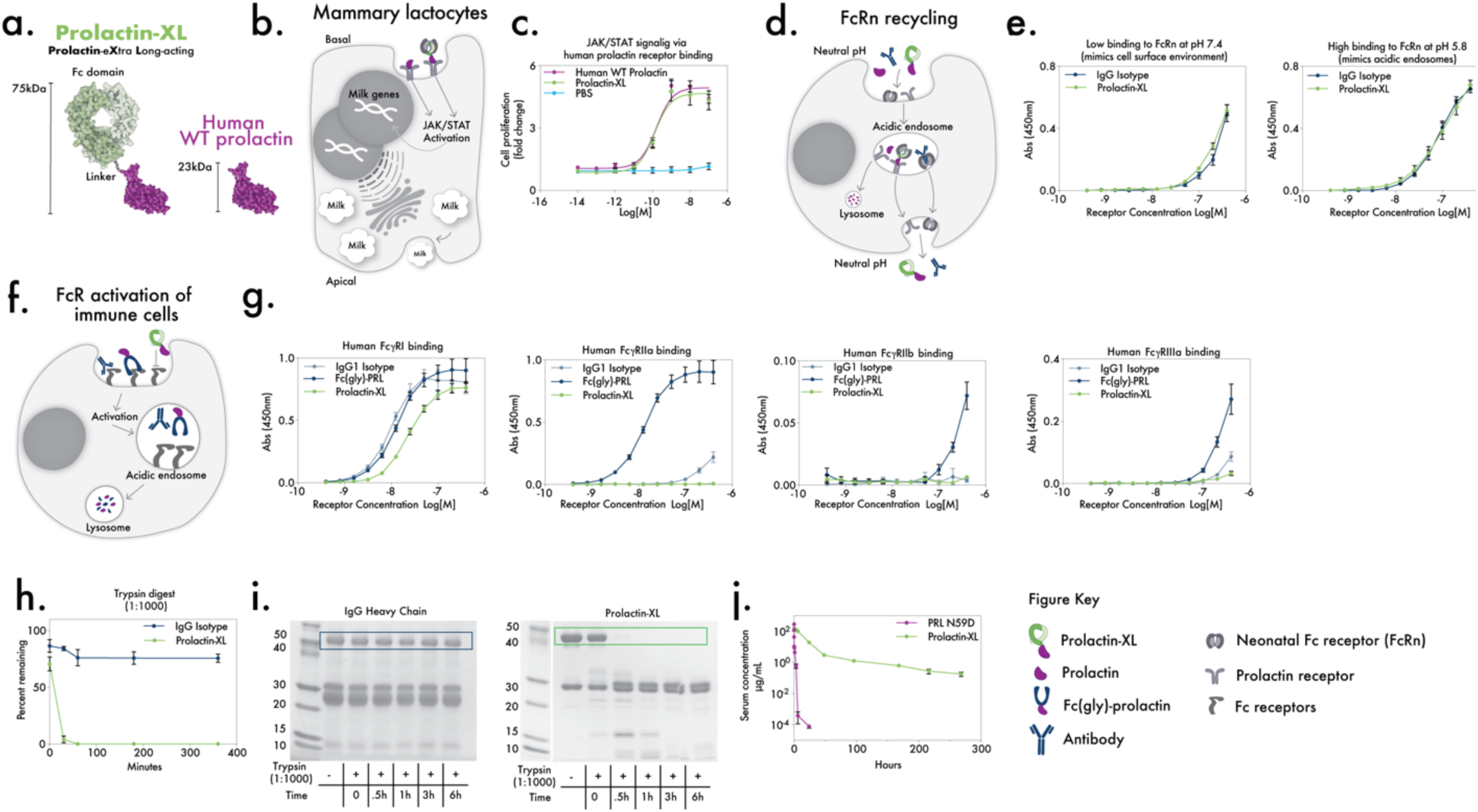
Prolactin-XL is the top performing variant in *in vitro* screens and has longest *in vivo* serum half-life. **a** A cartoon of Prolactin-XL compared to human WT prolactin. **b** A cartoon of a Fc-prolactin fusions binding to prolactin receptor (PRLR) to activate the JAK/Stat proliferation on mammary lactocytes. **c** An cell-based assay measuring Fc-prolactin fusions induction of *in vitro* signaling via human PRLR stably expressed in Ba/F3 cells. Data are represented as mean ± SEM of triplicates, and PRISM was used to fit non-linear curves. **d** A cartoon of FcRn-mediated recycling of Fc-prolactin fusions. **e** Binding of Fc-prolactin fusions to human FcRn:β2m at pH 7.4 or pH 5.8 measured by ELISA. Data are represented as mean ± SEM of triplicates. **f** A cartoon of Fc-prolactin fusions interacting with Fc receptors to activate immune effector cells. **g** Binding of Fc-prolactin fusions to human FcɣRI, human FcɣRIIa, human FcɣRIIb, or human FcɣRIIIa measured by ELISA. Data are represented as mean ± SEM of triplicates. IgG isotype control or Prolactin-XL were digested by Trypsin at a ratio 1:1000 (enzyme:protein). The percent of the remaining fusion were analyzed by SDS-PAGE (**i**) and measured by densitometry (**h**). Data are represented as mean ± SEM of triplicates. **j** Nulliparous Tg276 mice were injected with 5mg/kg I.V. of Prolactin-XL (n=4) and PRL N59D (n=5). Blood was collected by tail nick post injection, and the concentration of the fusions in serum was measured by ELISA. The data are depicted as mean ± SEM. PRISM was used to fit either a one-phase decay (PRL N59D) or a two-phase decay (Prolactin-XL).

#### Maintenance of FcRn recycling

Fc domains can increase serum half-life via at least two mechanisms: cellularly, by promoting endosomal recycling over lysosomal degradation (Fig. 1E), and at the organ level, by raising the construct’s molecular weight over 70kDa to prevent renal clearance^22^(Fig. 1D-II). Our engineered variants have molecular weights of ∼75kDa and contain Fc domain mutations previously shown to enhance FcRn-mediated endosomal recycling^34^. Variants were scored for their ability to bind to hFcRn *in vitro* at both neutral pH, to mimic cell surface conditions, and acidic pH, to mimic conditions in endosomes (Fig. 2E and Extended Data Fig. 6A-B).

#### Decreased off-target binding

There are 6 Fc receptors (FcRs) variably expressible on immune cells that bind Fc domains, then stimulate downstream signaling and receptor-mediated endosomal degradation^35^ (Fig 1F). The Fc domains of our engineered variants were modified to contain previously identified mutations shown to reduce off-target FcR binding^34,36,37^. Variants were scored for reduced off-target binding to FcRs *in vitro* (Fig. 2G and Extended Data Fig.1C-F).

#### Reduced transfer from mother to infant via reduced oral bioavailability

The Fc used in the fusions is a subdomain of IgG1; IgG1s have a natively low oral bioavailability (measured to be <25% in infants)^38,39^. Our variants were further engineered to be more digestible in the stomach by removing the F(ab) domain and de-glycosylating the Fc domain to enhance digestion by gastric proteases (Fig. 1C-II)^38^. Engineered variants were scored for their ability to be digested *in vitro* by the canonical gastric proteases, trypsin, chymotrypsin, and pepsin (Fig. 2H-I and Extended Data Fig. 7)^38^. These results were confirmed in *in vivo* pharmacokinetic studies in pups fed by dams dosed with an engineered variant (Extended Data Fig. 8D).

#### Improved serum half-life

In aggregate, the top-performing variants were Fc-PRL-3, Fc-PRL-7, Fc-PRL-13, and Fc-PRL-17. To confirm long-lasting bioavailability *in vivo*, nulliparous Tg276 mice which have mouse FcRn knocked-out and are hemizygotic for transgenic for hFcRn, were administered a single dose (intravenous 5mg/kg) of the Fc-prolactin fusions or the endogenously active, deglycosylated prolactin mutant (PRL-N59D). Then serum half-life (β-phase T_1/2_) was measured by ELISA (Extended Data Fig. 8A and E). Compared to PRL-N59D, Fc-PRL-13 was the best performer and had a 2,625-fold longer half-life (70.9h vs. 0.027h), 96-fold higher total drug exposure time (AUC_inf_), and 105-fold lower clearance rate (Fig. 2J and Table 1). Fc-PRL-13 was carried forward for *in vivo* studies and is hereafter called Prolactin-XL, short for Prolactin-eXtra Long-acting. Tg276 mice are difficult to breed making lactating studies logistically challenging. Therefore, C57bl/6j (B6) mice were used for further *in vivo* studies. The pharmacokinetic results for Prolactin-XL were confirmed in nulliparous and lactating C57bl/6j (B6) mice (Extended Data 8B).

**Table 1:**
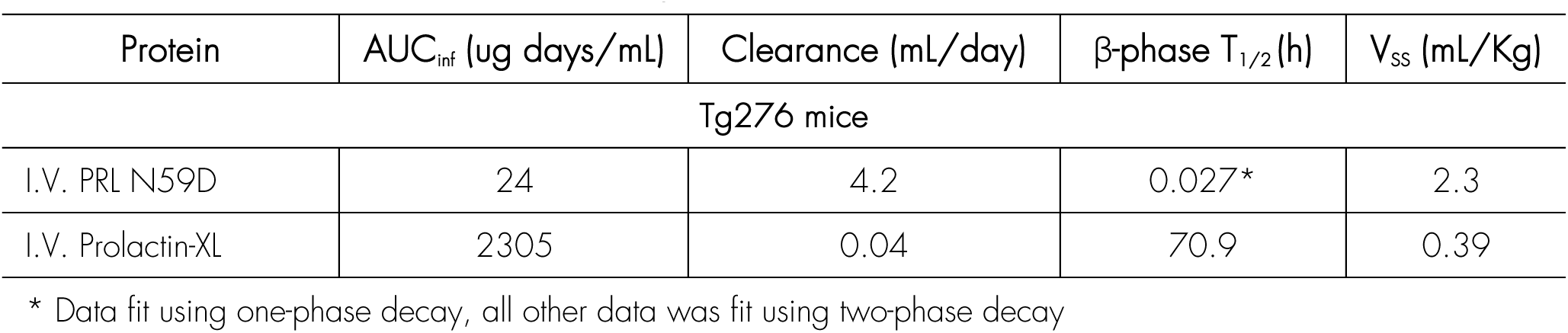
Pharmacokinetic parameters of Prolactin-XL and Prolactin N59D.

In subsequent studies to characterize *in vivo* efficacy, lactating C57bl/6j (B6) dams were administered subcutaneous doses of Prolactin-XL or vehicle (PBS) on day 7 post-partum. Litter numbers were normalized (n=5) at day 7, pups were weighed daily, and pup weight was used as a surrogate measure of milk production. Pathologically underweight pups were sacrificed when they weighed 20% less than positive controls; surviving pups were weaned at 21d.

### Prolactin-XL restores pharmacologically-ablated lactation

First, we assessed Prolactin-XL’s ability to restore lactation ablated by Bromocriptine (BR), a dopamine agonist prescribed off-label to inhibit lactation by disrupting native prolactin secretion^40^. Dams were injected twice-daily with BR (0.2mg) or vehicle (0.2mg tartaric acid dissolved in PBS with 20% ethanol) starting on day 7 post-partum. All pups fed by vehicle control-treated dams survive to 21d and are weaned (Fig. 3B-D and Table S4). By contrast, pups fed by dams receiving twice-daily BR treatment were pathologically underweight, and all pups were sacrificed by day 17. However, a single dose of Prolactin-XL on day 7 postpartum increased milk production for 2 days in BR-treated dams and 21% of pups survived until weaning (Fig. 3B-D and Table S4). Pups fed by +Prolactin-XL/+BR dams achieved maximum weight gain on day 9 post-partum and were 0.83g heavier (z-score of 0.93± 0.18 SEM) than pups fed by vehicle control-treated dams (Fig. 3B-D and Table S4). Repeat dosing of Prolactin-XL in BR-treated dams increased pup weight and survival in a dose-dependent manner (timing of Prolactin-XL treatment: every 2 days starting on day 7 postpartum; high dose: 5mg/kg, medium dose: 0.5 mg/kg, and low dose: 0.05mg/kg, all subcutaneous). 53% of pups survive until weaning when their BR-treated dams receive repeated high doses of Prolactin-XL, compared to 13% of pups for equivalent dosing of PRL-N59D (Fig. 3F-L and Table S6). At weaning, the surviving pups fed by BR-treated dams in the high dose +Prolactin-XL group achieved equivalent weight gain as pups fed by vehicle control-treated dams.

**Figure 3:**
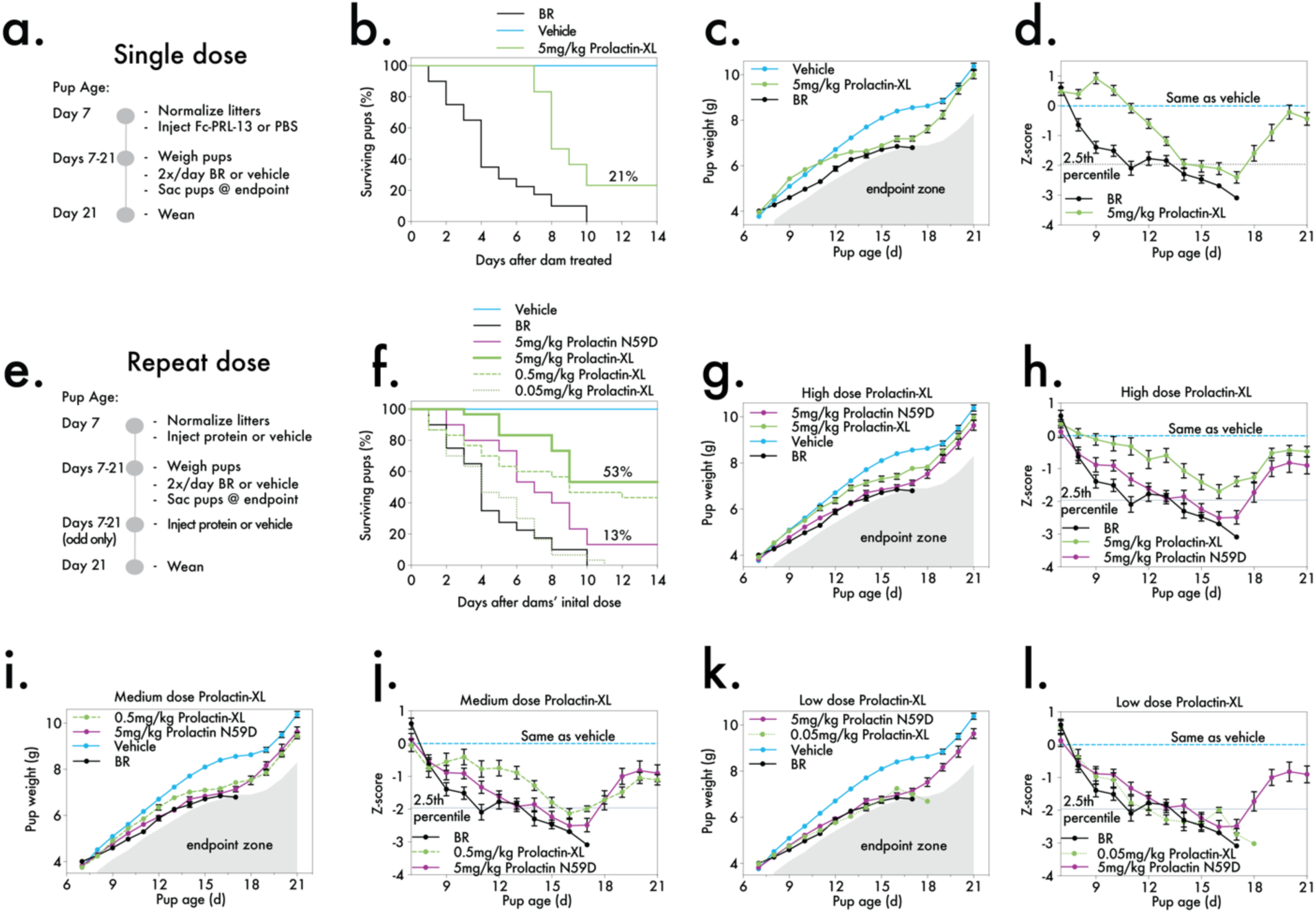
Prolactin-XL restores pharmacologically ablated lactation. **a** A timeline of a repeat dose experiment where the B6 dams are dosed with BR or vehicle (0.2mg tartaric acid dissolved in PBS with 20% ethanol) twice-daily (S.C.) and administered S.C. Prolactin-XL or vehicle (PBS) every other day. The dose groups are Vehicle (n=8), BR + 5mg/kg Prolactin-XL (n=6), or BR (n=8). The pups reach endpoint when they weigh 20% less than the pups in the vehicle control group. **b** The percentage of pups remaining in the study each day was recorded. The data are depicted as mean ± SE. **c** The weight of the pups was measured daily starting on day 7 post-partum (n=40 for Vehicle, n=30 for BR + 5mg/kg of Prolactin-XL, and n=40 for BR) and **d** the z-score for each pup was calculated. For **c** and **d**, the data are depicted as mean ± SEM. A Two-way ANOVA with Bonferroni correction for multiple comparison was used to calculate the statistical significance of pup weight gain, and the p-values are listed in Table S4. **e** A timeline of a repeat dose experiment where the B6 dams are dosed with BR or vehicle (0.2mg tartaric acid dissolved in PBS with 20% ethanol) S.C. twice-daily and S.C. administered Prolactin-XL, Prolactin N59D, or vehicle (PBS) every other day. The dose groups are Vehicle (n=8), Br + 5mg/kg Prolactin N59D (n=6), BR + 0.5mg/kg Prolactin-XL (n=6), BR + 0.05mg/kg Prolactin-XL (n=6), BR + 5mg/kg Prolactin-XL (n=6), or BR (n=8). The pups reach endpoint when they weigh 20% less than the pups in the vehicle control group. **f** The percentage of pups remaining in the study was recorded each day. The data are depicted as mean ± SE. **g**, **i** and **k** The weight of the pups was measured daily starting on day 7 post-partum (n=40 for Vehicle, n=30 for BR + 5mg/kg of Prolactin N59D, n=30 for BR + 0.05mg/kg of Prolactin-XL, n=30 for BR + 0.5mg/kg of Prolactin-XL, n=30 for BR + 5mg/kg of Prolactin-XL, and n=40 for BR) and the z-score for each pup was calculated (**h**, **j**, and **l)**. For **g**-**l**, the data are depicted as mean ± SEM. A Two-way ANOVA with Bonferroni correction for multiple comparison was used to calculate the statistical significance of pup weight gain, and the p-values are listed in Table S6.

### Prolactin-XL increases milk supply in mice with uncomplicated lactation accompanied by transient changes in mammary tissue

Next, we assessed Prolactin-XL’s ability to enhance milk supply in mice with uncomplicated lactation. Otherwise untreated dams were administered a single high dose (5mg/kg, subcutaneous or intravenous) of Prolactin-XL or vehicle (PBS) on day 7 post-partum. Under these conditions, all pups survive to weaning-age (Fig. 4B-C and Table S5), and on day 2 post-injection (day 9 post-partum), the pups fed by intravenously Prolactin-XL-treated dams were statistically heavier by 0.4g (z-score of 0.7± 0.6 SEM) than pups fed by vehicle control-treated dams (Fig. 4B-C, and Table S5). At weaning, the weight of the pups fed by the Prolactin-XL-treated dams was comparable to pups fed by the vehicle control dams (Fig. 4B-C and Table S5). Repeated high dosing of Prolactin-XL (every 2 days, 5mg/kg, subcutaneous) increases pup weight, although not statistically significantly. Pups weigh 0.6g (z-score of 0.4 ± 0.1 SEM) more than pups fed by vehicle control-treated dams at weaning (Fig. 4E-F and Table S7). These findings suggest that Prolactin-XL enhances lactation but does not cause pathological oversupply of milk or pathological weight gain of pups.

**Figure 4:**
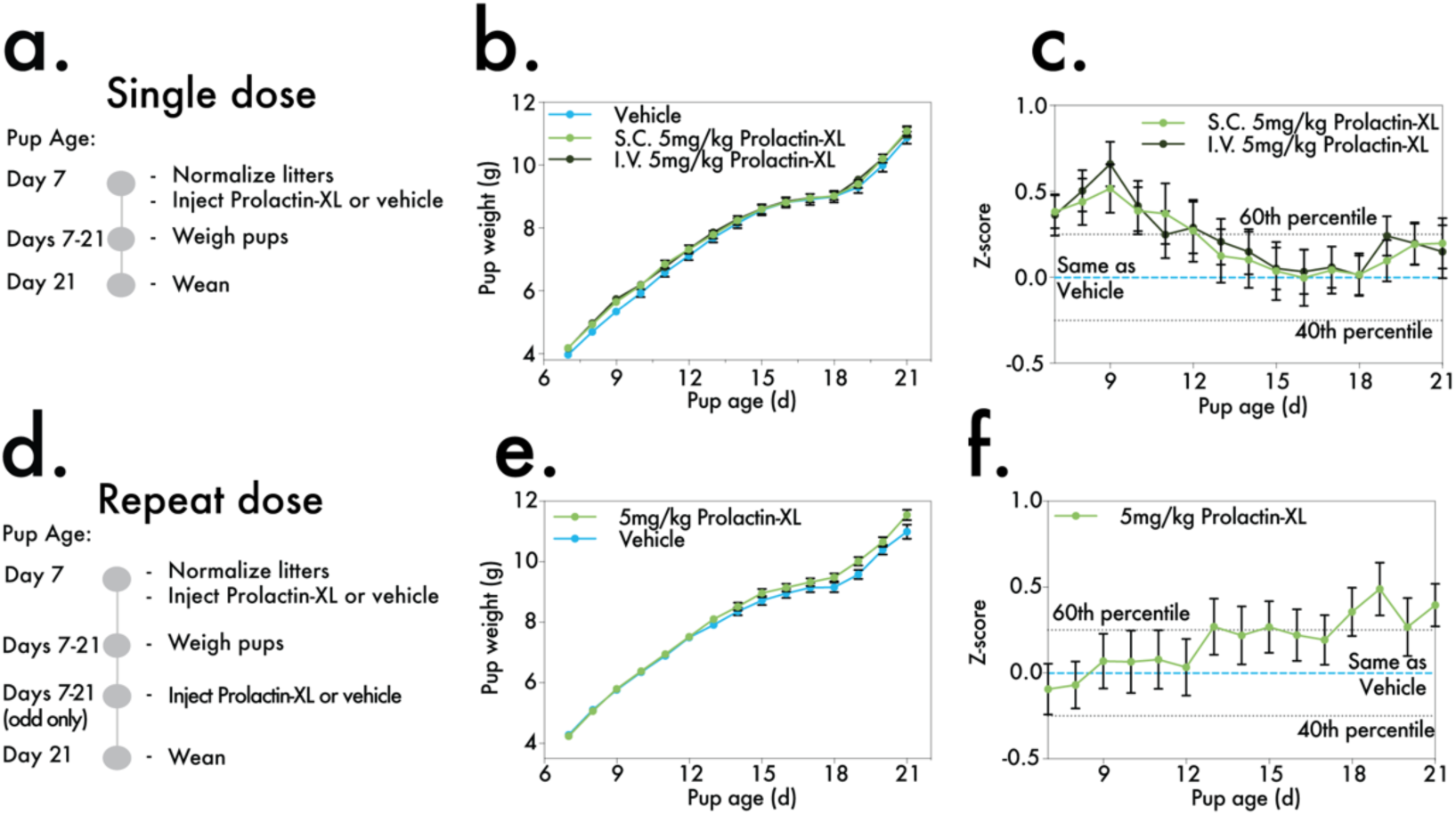
Prolactin-XL increases milk supply in mice with uncomplicated lactation. **a** A timeline of single dose experiment where B6 dams are given a single dose of subcutaneous or intravenous 5mg/kg Prolactin-XL or vehicle (PBS) on day 7 post-partum. The dose groups are vehicle (n=6), 5mg/kg I.V. Prolactin-XL (n=5), or 5mg/kg S.C. Prolactin-XL (n=7). **b** The weight of the pups was measured daily starting on day 7 post-partum (n=30 for vehicle, n=25 for I.V. 5mg/kg Prolactin-XL, and n=35 for S.C. Prolactin-XL) and **c** the z-score for each pup was calculated. For **b** and **c**, the data are depicted as mean ± SEM. A Two-way ANOVA with Bonferroni correction for multiple comparison was used to calculate the statistical significance of pup weight gain, and the p-values are listed in Table S5. On day 9 post-partum, the weight of pups fed by I.V. Prolactin-XL-dosed dams are significantly higher than the weight of pups fed by vehicle control dams. **d** A timeline of a repeat dose experiment where B6 dams are dosed with subcutaneously administered Prolactin-XL or vehicle every other day (n=6 and n=7, respectively). **e** the weight of the pups was measured daily starting on day 7 postpartum (n=30 for Prolactin-XL and n=35 for vehicle), and **f** the z-score for each pup was calculated. For **e** and **f**, the data are depicted as mean ± SEM. Multiple unpaired t-tests with Bonferroni correction for multiple comparison were used to calculate the statistical significance of pup weight gain, and the p-values are listed in Table S7.

Increased pup weight corresponds to transient, Prolactin-XL-dependent changes in dams’ mammary tissue. Repeatedly-dosed Prolactin-XL is measurable in serum up to 40µg/mL and penetrates mammary tissue as detected by ELISA (Extended Data Fig. 8B and Fig. 9A). However, JAK2/STAT5 activation and levels of β-casein, a principal milk protein, in the mammary tissue of Prolactin-XL-treated dams is indistinguishable from vehicle controls, suggesting that Prolactin-XL-induced signaling downstream of PRLR is not pathological (Extended Data Fig. 9B-F, J-M).

A board-certified rodent histopathologist found no pathologies in H&E-stained mammary tissues of Prolactin-XL-dosed dams. Marked changes in mammary tissue, consistent with a hypersecretory-lactation phenotype, are observed at day 10 post-partum between Prolactin-XL-and vehicle control-treated dams (Fig. 5D). These hypersecretory features resolve by weaning at day 21 coincident with involution of the mammary gland (Fig. 5D and Extended Fig. 9G-I, N-O and 10Q-X). The milk-producing alveoli within the mammary gland are composed of two primary cell types: milk-producing lactocytes, which are luminal epithelial cells marked by CK18, and myoepithelial cells, marked by CK14 and smooth muscle actin (SMA). CK18^+^ lactocytes are the primary drivers of the Prolactin-XL-induced organ-level morphology changes during lactation that peak at day 10 post-partum (Fig. 5D-M and Extended Data Fig.10A-I, EE-FF). We found no significant differences in CK14^+^/SMA^+^ myoepithelial cells, CD45^+^ lymphocytes, or other proliferating (Ki67^+^) cells (Extended Data Fig. 10J-P, Y-DD) across time, within treatments over time, or between treatments (i.e., Prolactin-XL vs. vehicle controls). These observations suggest that Prolactin-XL acts specifically to increase milk production without producing off-target pathology in the mammary gland.

**Figure 5:**
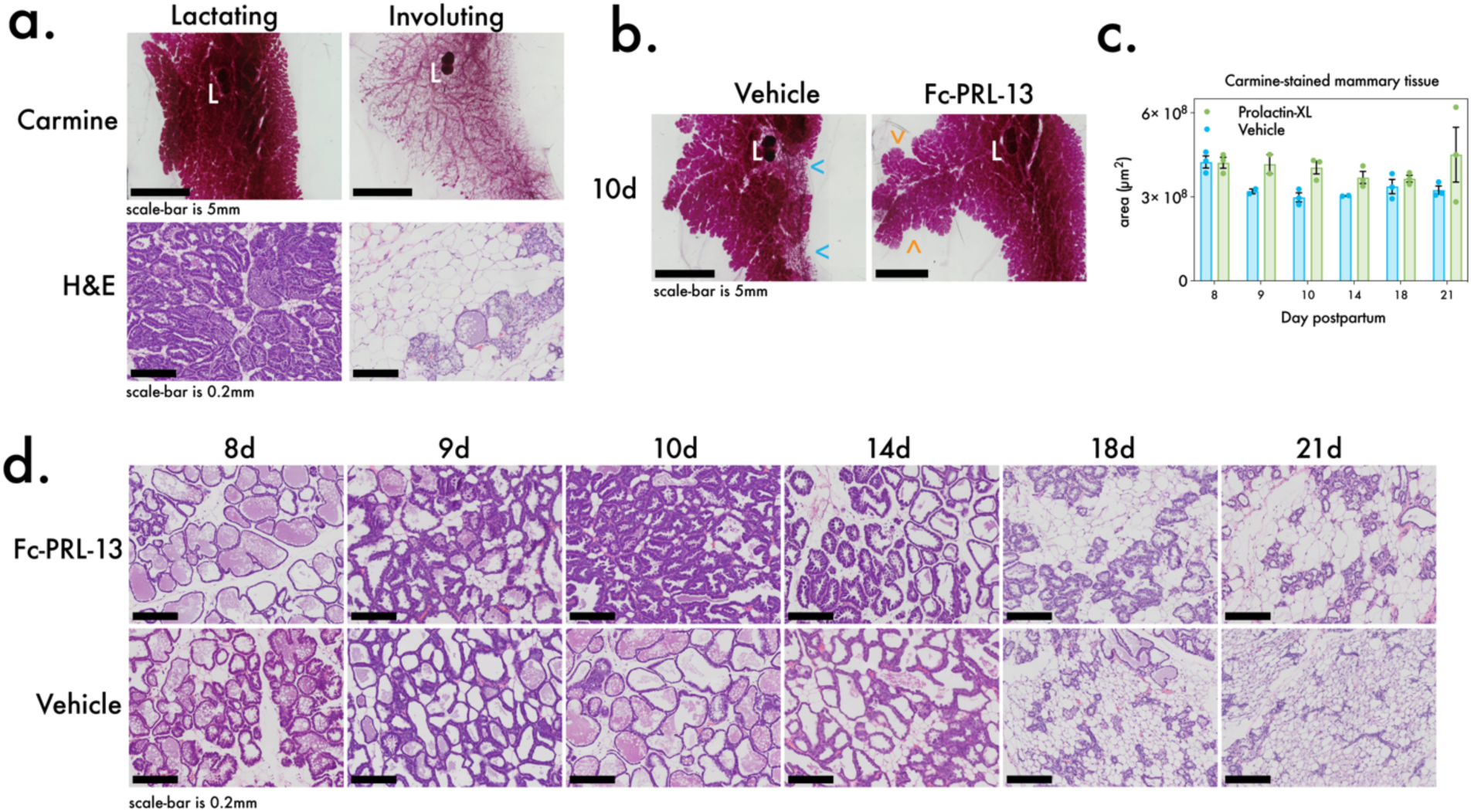
Prolactin-XL induces transient changes in mammary tissue in mice with uncomplicated lactation. **a** Examples of lactating (left) and involuting (right) mammary glands stained with carmine (top; scale bar=5mm) or H&E (bottom; scale bar=0.2mm). The lymph node is labeled (white L). For **b**-**m**, lactating mice (n=3) were repeatedly dosed with subcutaneously administered vehicle (PBS) or Prolactin-XL (5mg/kg) every other day beginning on the 7^th^ day postpartum. Litters were normalized (n=5) on the 7^th^ day postpartum. Mammary glands from the lactating mice were collected on 8d, 9d, 10d, 14d, 18d, 21d post-partum for subsequent analysis. Pups are weaned on the 21^st^ day postpartum. **b** Whole mount mammary glands were carmine-stained from (n=3 except for 9d Prolactin-XL and 14d vehicle where n=2). Representative images are depicted. In this example on the 10^th^ day post-partum, the vehcile-dosed mouse has a duct with low density alveoli branching out from it (blue <; scale bar =5mm). Whereas the Prolactin-XL dosed mouse has densely packed alveoli and an additional branch of alveoli highlighted (orange <; scale bar = 5mm). **c** The area of the same whole mount, carmine-stained mammary glands as in **b** was measured using Qupath. The data are depicted as mean ± SEM. **d** Whole mammary glands formalin-fixed, paraffin-embedded, and stained with H&E (n=3 per time point). Representative images are depicted (scale bar = 0.2mm).

## Discussion

Human infants are born to breastfeed. Despite 50% of lactating persons struggling to make milk, there are no approved drugs to enhance lactation.^1^ Breastfeeding is the biological norm, the everyday stand of care, and a key element of emergency preparedness^2,9,14^. Previous clinical studies identified recombinant prolactin, the natural hormonal driver of lactation, as a drug candidate to enhance lactation^21^. However, its short serum half-life requires twice-daily injections and limits is real-world viability as a drug candidate^19–21^.

Here, we engineer a long-acting human IgG1 Fc domain-human prolactin fusion, Prolactin-XL. Our pioneering studies in mice show Prolactin-XL has a long-lasting serum half-life and enhances lactation. Prolactin-XL has a 2,625-fold longer half-life than Prolactin (70.9h vs. 0.027h). We demonstrate its efficacy in stimulating lactation in mice and restoring growth of pups fed by dams with pharmacologically-ablated lactation. We propose Prolactin-XL as a potential first-in-class therapeutic that should be further developed to combat lactation insufficiency.

We hope Prolactin-XL can help lactating persons enhance their milk supply in a world inadvertently designed to disrupt human milk production. The key biology underlying lactation is a positive feedback loop between infant suckling mechanically stimulating pituitary prolactin production and milk production^41,42^. It requires constant stimulation on the timescale of several hours because prolactin is quickly cleared by the kidneys^20^. This key biology is easily disrupted because our societies are not built around stimulating this time-sensitive positive feedback loop. For example, it is still common medical practice to separate lactating persons from their newborns immediately following delivery; most maternity wards are physically separated from the nurseries, NICU’s, and pediatric departments^43^. From the onset of lactation, the positive feedback loop is already being disrupted. Lactating persons are shunned from breastfeeding in public spaces. Without adequate parental leave, most lactating persons are forced to return to the workforce where they are separated from their infants for the entire workday^44–48^. Sustained separation of lactating persons from their infants is the engrained norm. Lactating persons need more lactation support, novel technologies, and societal shifts to help them reach their breastfeeding goals.

Lactation has largely been studied in a limited number of biological contexts – mostly in postpartum female mammals. This is the easiest and most obvious first step because postpartum female mammals have a higher prevalence of lactation. But to exhaustively understand mammalian mammary gland biology across the full spectrum of biological variation, we need to recontextualize lactation. Thinking of lactating as only a ‘female postpartum phenomenon’ hinders us from building a deeper understating of biology. For example, prolactin drives milk production in males of some mammalian species including bats, flying foxes, and goats^49–52^. In humans, males, intersex individuals, adoptive parents, trans people, post-menopausal people, and newborns of all sexes can make milk^53–58^. Prolactin even drives milk production in some non-mammals such as male emperor penguins and in pigeons and flamingos of both sexes^59–61^. This underscores the diverse biological contexts where mammary glands or other nutritive-secreting organs are stimulated by prolactin to make milk without being coupled to pregnancy and female sexes.

Until now, there has been no reliable tool to study enhancing lactation across these diverse biological contexts. Because both Prolactin and Fc are highly conserved across all mammals, and with few exceptions, mammary glands are found in almost all mammalian sexes^19^, Prolactin-XL and homologous fusions are also a powerful set of tools to study the basic biology of scaling-up lactation. Exogenous prolactin alone does not increase milk production in rodents, ruminants and swine, primarily because its cleared too quickly^62–64^. In humans, exogenous prolactin only increases milk because of frequent daily injections^21^. Prolactin-XL and homologous fusions can easily be probed or imaged, like research antibodies, and they are compatible with most state-of-the-art research techniques. We look forward to seeing our technology applied to future lactation studies that encompass the broader contexts and full spectrum of biological variation and possibilities.

Long-acting prolactins are not only important for understanding mammary gland biology but have broad implications for conservation of endangered mammals. There is no artificial milk substitute for most mammals besides humans. Many zoos and rehabilitation centers that care for young born to mothers with insufficient milk supply primarily rely on dairy. Dairy cannot meet the nutritional requirements of all species of mammalian young. For example, cow’s milk is about 3% fat whereas orca’s milk is about 30% fat^65^. Not only has breastmilk been evolutionarily tailored to meet the nutritional needs of the young, but it provides personalized immunological support and specialized fuel for the development of gastrointestinal microbiota via antibodies and milk oligosaccharides^66^. We propose that homologous fusions to Prolactin-XL should be developed to enhance lactation in not just endangered mammals, but all mammals.

Additionally, enhancing milk supply in livestock has broad implication for sustainable agriculture. Dairy is an emission-heavy product to begin with, and infant formulas have added emissions due to the intense manufacturing process, worldwide transportation, packaging, and additional water use^13,14,48,67–71^. Homologous fusions to Prolactin-XL could sustainably enhance milk production per animal to decrease the overall number of animals required to produce the same amount of milk. Additionally, Prolactin-XL could reduce human reliance on infant formula which would reduce our infant formula-associated emissions. Prolactin-XL and homologous fusions have the potential to greatly reduce our agriculture-based impacts on the planet that are accelerating climate change.

### Lactation has deep evolutionary roots but is also uniquely and beautifully human

Breastfeeding touches the lives of everyone on the planet is some way. Breastfeeding care-work builds people, families, friendships, and communities. There is universal consensus amongst scientists, clinicians, major health organizations, national governments, and the infant formula industry to protect and promote everyday breastfeeding^1–13^. As climate disasters are intensifying in scale, frequency and intensity, breastfeeding becomes even more life-and-death for vulnerable infants in emergencies^9,14^. Breastfeeding is the safest option when there is no electricity, no safe water, disrupted supply chains, or infant formula shortages, contaminations and recalls^9,14–16,18^. These emergencies disproportionally impact marginalized communities globally. They have borne most of the intersecting burdens of climate change, systemic and structural racism, racial healthcare disparities, and inadequate lactation support^1,17,48,72–76^. We hope our Prolactin-XL can be an everyday bolster for breastfeeding and stockpiled for emergency situations, like vaccines and essential medicines. Doing so would make our fragile infant feeding systems more resilient which is imperative for advancing health equity worldwide.

## Authorship contributions

KK, KC, and JW conceived of the long-acting, Fc-prolactin fusion. KK and JW designed initial constructs, and KK and KD designed experiments. KD, KC, JW, CP, and QJ gave technical support and conceptual advice. KK performed all experiments, analyzed data, and wrote the main paper with QJ. KK wrote the Supplementary Information and Extended Data. All authors discussed the results and implications, and commented on the manuscript. CP and PS supervised the project.

## Acknowledgements

This research was sponsored by the Blavatnik Biomedical Accelerator at Harvard Medical School and the Wyss Institute for Biologically Inspired Engineering. Kasia Kready is supported by a Herchel Smith Graduate Fellowship, a Pharmacological Sciences Training Grant (5T32GM132089-03), and a Fuji Fellowship. Kaillyn Doiron is also supported by a Fuji Fellowship. We thank Dana-Farber/Harvard Cancer Center in Boston, MA, for the use of the Rodent Histopathology Core, which provided preparation of histology slides, slide interpretation and histopathological interpretation services. Dana-Farber/Harvard Cancer Center is supported in part by a NCI Cancer Center Support Grant #NIH 5 P30 CA06516. The authors would also like to thank the Neurobiology Imaging Facility at Harvard Medical School, Boston, MA for imaging histology sections. We sincerely thank Amanda Graveline, Andyna Vernet, Sarai Bardales, and Melinda Sanchez from the Wyss Institute at Harvard for their technical support with the animal experiments.

## Data Availability

All sequences used in this work are listed in the supplement. All the other data that support the findings of this study are available from the corresponding author upon request.

## Code Availability

No original code was generated for this work. Scripts used to generate figures will be made available to reviewers upon requestion.

## Methods

### DNA constructs

The DNA sequence for human PRL wild-type was derived by reverse translating and codon optimizing the protein sequence from UniProt (P01236). The PRL deglycosylated mutant N59D was constructed by introducing a codon change in the wild-type sequence. The DNA sequence for mouse PRL wild-type was derived by reverse translating and codon optimizing the protein sequence from UniProt (P06879). The DNA sequence for human and mouse PRLR wild-type was derived by reverse translating and codon optimizing the protein sequences from UniProt (P16471 and Q08501, respectively). The DNA sequence for human IgG1-Fc wild-type was derived by reverse translating and codon optimizing the protein sequence from UniProt (P0DOX5). It was modified (C220S) to remove the unpaired cysteine in the hinge region and a protease cleavage site at the c-terminus (K447A) by making a codon change in the wild-type sequence. This modified IgG1-Fc sequence was used as the base for the Fc-PRL fusions. Fc-PRL fusions were generated by fusing the IgG1-Fc sequence to PRL N59D. The following features of the Fc-PRL fusions were varied and introduced into the Fc-PRL fusions by codon changes:

(1) The order of PRL fused to Fc (i.e., N-PRL-Fc-C or N-Fc-PRL-C)
(2) Use of linker between prolactin and Fc (GGsGG)
(3) Use of Fc heterodimer (Fc Knob (T266W) fused to Fc Hole (T366S, L368A, Y407V)^30,31^ or Fc Zw1 A (T350V, L351Y, F405A, Y407V) fused to Fc Zw1 B (T350V, T366L, K293L, T394W)^32^
(4) Fc-hole without Protein A binding to get rid of unwanted homodimers (‘RF’ mutations: H435R, Y435F)^77^
(5) Use of mutations to Fc to enhance half-life (i.e., “EDHS” mutations: V264E, L309D, Q311H, N434S or “YTE” mutations: M252Y, S254T, T256E)^34^
(6) Use of mutations to Fc to decrease FcR binding (i.e., “LALA” mutations: L234A, L235A or “LALAPG” mutations: L234A, L235A, P329G)^78^

The DNA sequence for human FcRg1, FcgRIIa, FcgRIIb, and FcgRIIIa were derived by reverse translating and codon optimizing the protein sequences from UniProt (P12314, P12318, P31994, and P08637 respectively). The DNA sequence for mouse FcgRI, FcgRIIb, FcgRIII, and FcgRIV were derived by reverse translating and codon optimizing the protein sequence from UniProt (P26151, P08101, P8508, and A0A0B4J1G0, respectively). See Supplementary Information for individual sequences.

### Mammalian cell culture

FreeStyle 293-F and Ba/F3 cell lines were obtained from Dr. Jungmin Lee (Harvard Medical School), and Hek293-T cell lines were obtained from Rui Truong (Harvard Medical School).

FreeStyle 293-F cell lines were cultured in FreeStyle 293 Expression Medium (Thermo Fisher Scientific; 12338026). Hek293-T cell lines were cultured in DMEM (ATCC; 30-2002) supplemented with 10% FBS (Thermo Fisher Scientific; 10082147), 1% Penicillin-Streptomycin (Thermo Fisher Scientific; 15140122), 1x Sodium Pyruvate (Thermo Fisher Scientific; 11360070), 1x MEM Non-essential Amino Acids (Thermo Fisher Scientific; 11140050), and 1x HEPES (Thermo Fisher Scientific; 15630080). Ba/F3 cell lines were cultured in RPMI 1640 (ATCC; 30-2001) supplemented with 10% FBS (Thermo Fisher Scientific; 10082147), 1% Pen-Strep (Thermo Fisher Scientific; 15140122), and 10ng/mL mouse IL-3 (PeproTech; 213-13), unless specified otherwise. FreeStyle 293-F cells were cultured at 37C in 8% CO2 with shaking at 125 RPM (volumes 30-90mLs) and 90 RPM (volumes120-1000mL). Hek 293T cells and Ba/f3 cells were cultured at 37C in 5% CO2.

### Yeast strains and cultivations

NRRL Y-11430 was obtained from (ATCC; 76273). Strains were generated to co-express the heterodimeric Fc-PRL-13 fusion using pPICZ A obtained from Amazir Bredl (Harvard Medical School). Competent cells were prepared and transformed as described elsewhere^79^. Strains were first generated to express the Fc-only monomer under the control of the *DAS2* promoter and with Kanamycin resistance. Cells were allowed to recover overnight at room temperature without shaking and then plated on YPD agar plates supplemented with 200ug/mL G418 (Invivogen; ant-gn). Plates were incubated at 30C until colonies formed. All of the colonies were then scraped off the plate, made competent, and transformed to express the Fc-PRL monomer under the control of the *AOX1* promoter and Zeocin resistance. Cells were allowed to recover overnight at room temperature without shaking and then plated on YPD agar plates supplemented with 200ug/mL G418 and 200ug/mL Zeocin (Invivogen; ant-zn). Plates were incubated at 30C until colonies formed.

For protein production from the strains, BMGY and BMMY media was prepared. 1L of BMY media was prepared with 900mL of miliQ water, 100mL of 10x Potassium Phosphate Buffer (22.99g dipotassium phosphate (Sigma; 60353) and 118.14 g monopotassium phosphate (Sigma; P5655) in 1L of miliQ water, pH 6.5), 10g yeast extract (Gibco; 212750), 20g peptone (Sigma; P5905), 13.4g yeast nitrogenous base without amino acids (Sigma; Y0626), and 3.07g of L-glutathione (Simga; G4251). BMY media was then sterilized in the autoclave, and after cooling it was filtered using a 0.2uM filter. To prepare 1L of BMGY media, 40mL of 100% glycerol (Sigma; G5516) was added to BMY media. To prepare 1L of BMMY media, 15mL of methanol (Fisher Scientific; AC325740025) was added to BMY media. To screen for protein expression, colonies co-expressing both plasmids and the WT strain were grown in 24-well deep well plates (25C, 300RPM) with 3mL of BMGY media for 24hrs of biomass accumulation. Cells were then pelleted and resuspended in 3mL of BMMY media and allowed to grown for an additional 24 hrs. Cells were pelleted and samples of the supernatant were collected to analyze expression of Fc-PRL-13 via ELISA.

To quantify the amount of Fc-PRL-13 expressed by the strains, 50ul of 0.1ug/mL anti-human PRL antibodies (Abcam; ab244037) were diluted in PBS and used to coat MaxiSorp 96-well ELISA plates (Sigma; M9410) for 1hr at room temperature. Plates were washed 3 times in PBST (PBS with 0.05% Tween 20 (Sigma; P2287), and then the plates were blocked with 200uL of blocking buffer (PBS with 3% BSA) overnight at 4C. Plates were again washed 3 times in PBST. 50uL of supernatant diluted in PBS (1/100, 1/1000, 1/10,000) was added to the plates and incubated at room temperature for 1hr. Plates were then washed 5 times in PBST. 50uL of HRP-conjugated anti-human IgG Fc (Invivogen; 31413) diluted 1:5000 in blocking buffer was added to the plates and incubated at room temperature for 1hr. Plates were then washed 5 times with PBST. 100uL of TMB Substrate Solution (Thermo Fisher Scientific; 34028) equilibrated to room temperature was added to the plates and incubated for 5.5 minutes or until desired color develops. Reactions were stopped by adding 100uL of 2M Sulfuric Acid to each well. The absorbance was measured at 450nm. A standard curve was generated by serially diluting purified Fc-PRL-13 into PBS. Prism (GraphPad Prism) was used to fit a sigmoidal curve to the data and used to quantify Fc-PRL-13 in the supernatant samples. The clone with the highest expression of Fc-PRL-13 was chosen for large-scale protein production.

### Recombinant protein expression and purification from mammalian cells

For small batches (<10ugs), proteins were transiently expressed in FreeStyle 293-F cells using pSecTag2A plasmids and 293Fectin transfection reagent according to the suppliers’s protocol (Thermo Fisher Scientific; 12347019). 4-6 days after transfection, protein levels in the supernatant were assayed by SimplyBlue (Thermo Fisher Scientific; LC6065) stained SDS-PAGE and western blots using either anti-6xHis-HRP antibodies (Abcam; ab1187) or HRP-conjugated anti-human IgG Fc (Invivogen; 31413). Proteins from transient transfections were either purified via His-tag or Protein A purification. His-tagged proteins were purified with His60 Ni Superflow resin (TaKaRa; 635677) according to the manufacturer’s instructions, and Fc fusions were purified with Protein A agarose resin (Thermo Fisher Scientific; 20334) according to the manufacturer’s instructions. Cell supernatant and each purification fraction were run on Tris-glycine gels, 4-20% (ThermoFisher; XP04205BOX) under reducing and denaturing conditions, and then stained with SimplyBlue (Thermo Fisher Scientific; LC6065).

Fractions containing eluted proteins were combined, concentrated, and de-salted into endotoxin-free PBS (Teknova; P0300) using Macrosep Advanced centrifugal devices (VWR; 891310980). The purity of the top 4 Fc-PRL fusions (Fc-PRL 3,7, 13, and 17) were confirmed by size exclusion chromatography and was over 95%. Protein concentration was measured by BCA assay (Thermo Fisher Scientific; 23227) according to the manufacturer’s instructors.

Proteins were stored at 4C throughout the described process, ultimately stored as single-use aliquots at −80C, and thawed once before use. Only endotoxin-free reagents were used. Size exclusion chromatography (SEC) was performed using a Superdex 200 Increase 10/300GL column (Cytiva; 28990944) on an Äkta Pure 25 HPLC system and analyzed using Unicorn v.6.3 software.

### Recombinant protein expression and purification from yeast

For large-scale batches, Fc-PRL-13 was produced from strain Fc-PRL-13-c6. Fc-PRL-13-c6 was grown in 200mL of BMGY media supplemented 1tab/50mL of cOmplete Protease Inhibitor Cocktail (Sigma; 11836145001) and 10uM Pepstatin A (Sigma; P5318) in baffled 2L flasks. Cells were incubated for 24hrs of biomass accumulation at 25C and 300RPM. Cells were then pelleted and resuspended with 200mL of BMMY media supplemented with 1tab/50mL of O Complete Protease Inhibitors and 10uM Pepstatin A. Cells were again incubated for 24hrs of biomass accumulation at 25C and 300RPM. Fc-PRL-13 levels in the supernatant were assayed by Coomassie Blue stained SDS-PAGE, Western blots using HRP-conjugated anti-human IgG Fc (Invivogen; 31413) diluted 1:10000 in blocking buffer, or ELISA using anti-PRL capture antibodies and anti-IgG Fc detection antibodies as previously described. Fc-PRL-13 was purified with Protein A agarose resin (Thermo Fisher Scientific; 20334) according to the manufacturer’s instructions with the exception that all buffers were supplemented with 1tab/50mL of cOmplete Protease Inhibitor Cocktail and 10uM Pepstatin A. Cell supernatant and each purification fraction were run on Tris-glycine gels, 4-20% (ThermoFisher; XP04205BOX) under reducing and denaturing conditions, and then stained with SimplyBlue (Thermo Fisher Scientific; LC6065). Fractions containing eluted proteins were combined, concentrated, and de-salted into endotoxin-free PBS (Teknova; P0300) using Macrosep Advanced centrifugal devices (VWR; 891310980). The purity of the top Fc-PRL-13 was confirmed by anion exchange and was over 95%. Protein concentration was measured by nanodrop (Emolar = 96260 M-1cm-1 and Mw = 75.46 kDa). Proteins were stored at 4C throughout the described process, ultimately stored as single-use aliquots at −80C, and thawed once before use. Only endotoxin-free reagents were used.

### ELISA assay of FcRn binding

50uL of Fc-PRL fusions or IgG1 isotype control were diluted in PBS to 2ug/mL were used to coat MaxiSorp 96-well ELISA plates (Sigma; M9410) and incubated overnight at 4c. Plates were then washed 3 times with PBST (PBS with 0.05% Tween2 (Sigma; P2287)), and then the plates were blocked with 200uL of blocking buffer (PBS with 3% BSA) for 1hr at room temperature. Plates were again washed 3 times with PBST. Plates were then incubated 1hr at room temperature with 50uL serially diluted His-tagged human and mouse FcRs starting at 400nM and diluted 1:2. Plates were then washed 3 times with PBST (pH 7.4 or 5.8). 50uL of anti-6xHis-HRP antibody (Abcam; ab1187) diluted 1:10000 in PBS (pH 7.4 or pH 5.8) was added to the plates and incubated 1hr at room temperature. Plates were then washed 3 times with PBST (pH 7.4 or 5.8). 50uL of TMB Substrate Solution (Thermo Fisher Scientific; 34028) equilibrated to room temperature was added to the plates and incubated for 1min or until desired color develops. Reactions were stopped by adding 50uL of 2M Sulfuric Acid to each well. The absorbance was measured at 450nm. Prism (GraphPad Prism) was used to fit a sigmoidal curve to the data and used to determine the Log(EC_50_) of the pH 5.8 curves only. Reported data represent mean +/- SEM of three replicates.

### ELISA assay of FcR binding

50uL of Fc-PRL fusions or IgG1 isotype were control diluted in PBS to 2ug/mL were used to coat MaxiSorp 96-well ELISA plates (Sigma; M9410) and incubated overnight at 4c. Plates were then washed 3 times with PBST (PBS with 0.05% Tween2 (Sigma; P2287)), and then the plates were blocked with 200uL of blocking buffer (PBS with 3% BSA) for 1hr at room temperature. Plates were again washed 3 times with PBST. Purified His-tagged human and mouse FcRs were prepared as previously described. Plates were then incubated 1hr at room temperature with 50uL serially diluted FcRs starting at 400nM and diluted 1:2. Plates were then washed 3 times with PBST. 50uL of anti-6xHis-HRP antibody (Abcam; ab1187) diluted 1:10000 in PBS was added to the plates and incubated 1hr at room temperature. Plates were then washed 3 times with PBST. 50uL of TMB Substrate Solution (Thermo Fisher Scientific; 34028) equilibrated to room temperature was added to the plates and incubated for 1min or until desired color develops. Reactions were stopped by adding 50uL of 2M Sulfuric Acid to each well. The absorbance was measured at 450nm. Reported data represent mean +/- SEM of three replicates.

### Protease degradation study

Trypsin (Promega; V5111), chymotrypsin (Promega; V1061), and pepsin (Promega; V1959) were resuspended according to the manufacturer’s instructions. Trypsin, chymotrypsin, or pepsin were added on ice to Fc-PRL fusions or IgG1 isotype control at 1:1000, 1:100, and 1:5000, respectively. Samples were vortexed for 10s, and then a 3-5uL aliquot was taken for timepoint 0. Reactions were then incubated on a thermocycler at 25C (for chymotrypsin) or 37C (for trypsin and pepsin). 3-5uL aliquots were taken at 25mins, 50mins, 4hrs, 6hrs, and 24hrs, and stored at −20C. 2x Laemmli (Bio Rad; 1610737) with 5% BME was added to all aliquots at 1:1 ration and boiled at 100C for 10mins. The aliquots were run on Tris-glycine gels, 4-20% (ThermoFisher; XP04205BOX) under reducing and denaturing conditions, and then stained with SimplyBlue (Thermo Fisher Scientific; LC6065). The percent of remaining protein was analyzed by densitometry on Image Lab software. Reported data represent mean +/- SEM of three replicates.

### PRLR signaling cell-based assay

Stable cell lines expressing human or mouse PRLR were generated by lentiviral transduction followed by 0.7 ug/mL puromycin selection (Sigma; P9620). 12ug of each pTwist Lenti Puro SFFV WPRE lentiviral construct encoding either human or mouse PRLR were co-transfected with 9ug of psPAX2 (Addgene; 12260) and 3ug of pMD2.G (Addgene; 12259) 2^nd^ generation lentiviral packaging plasmids into HEK 293T cells at 60-80% confluency with 60uL Lipofectamine3000 (Thermo Fisher Scientific; L3000008) and 50uL P3000 reagent (Thermo Fisher Scientific; L3000008) diluted into 2.5mL of Opti-Mem (Thermo Fisher Scientific; 31985070). After 4hrs, the media was aspirated and replaced with 15mL of culture media (DMEM, 10% FBS, 1% PS, 1x sodum pyruvate, 1x MEM NEA, and 1x HEPES). Supernatant containing lentivirus was collected after a subsequent 24hrs and 48hrs and combined.

Lentivirus-containing supernatant was centrifuged at 400xg for 5 mins to pellet cell debris and then filtered with a 0.45um Durapore PVDF membrane steriflip (Coleparmer; EW-29969-26). 4x Lenti-X concentrator reagent (TaKaRa; 631231) was added to the lentivirus-containing supernatant at a final concentration of 1x and incubated at 4C for 1hr. After incubation, the samples were centrifuged at 1500xg for 45mins. Then the supernatant was removed and the white virus pellet was resuspended in 1mL of culture media (DMEM, 10% FBS, 1% PS, 1x sodum pyruvate, 1x MEM NEA, and 1x HEPES). Single use aliquots were stored at −80C until use. Two days before transduction, 250,000 Ba/F3 cells were plated in 6 well-plates. Cells were transduced by replacing media with 20-250uL of concentrated virus, 0.4mL 10x polybrene (Sigma; TR-1003-G), and complete media without antibiotics (RPMI 1640, 10% FBS and 10ng/mL mouse IL3) to a final volume of 3.6mL. After 48hrs incubation with the virus, cells were expanded into 2mL of complete media supplemented with 0.7ug/mL puromycin and 10ng/mL purified human prolactin or mouse prolactin. After selection recovery, stable pools of cells were analyzed by flow cytometry using APC anti-Flag-tag antibody (Biolegend; 637307). Stable pools went through another round of selection in complete media supplemented with with 0.7ug/mL puromycin and 1ng/mL human prolactin or purified mouse prolactin and were analyzed by flow cytometry. Highest expressing stable pools were used for the cell-based PRLR proliferation assay.

The cell-based PRLR proliferation assay was modified from elsewhere^80^. 100uL of Ba/F3 cells stably expressing human or mouse PRLR were seeded in 96-well plates at 50,000 cells/well in RPMI media supplemented with 10% FBS, 1% PS, and 0.7ug/mL puromycin. The cells were incubated for 6hrs at 37C, 5% CO2. 100uL of 2x Fc-PRL fusions, human prolactin wild-type, human prolactin N59D mutant, or mouse prolactin wild-type serially diluted in PBS was added to the plate and incubated for 48hrs at 37C, 5% CO2. 20uL of WST-1 (Thomas Scientific; C755B06) or MTS (Abcam; ab197010) was added to each well and incubated at 37C, 5% CO2 for 1-4hrs until the desired color developed. The absorbance was measured at 450nm and 650nm for WST-1 and 490 for MTS. Prism (GraphPad Prism) was used to fit logistic curves to the data and generate Log(EC_50_) and Emax values. Reported data represent mean +/- SEM of three replicates.

### Pharmacokinetic studies in non-lactating mice

All animal work was approved by the Harvard Medical School IACUC under protocol IS00003310. The animal experiment was modified from elsewhere^34^. 2019. On day 0, Tg276 mice (Jackson; 004919) or C57bl/6j mice (Jackson; 000664) were administered a 5mg/kg dose of Fc-PRL fusions or PRL N59D either by intravenous injection into the tail vein or by subcutaneous injection. Blood samples were collected from the tail vein at various time points.

The serum concentrations of the Fc-PRL fusions were determined using a quantitative ELISA as previously described. Anti-human PRL antibodies were coated onto MaxiSorp 96-well ELISA plates to capture Fc-PRL fusions from 50uL of serially diluted serum. HRP-conjugated anti-human IgG1 Fc antibodies were used for detection. A standard curve was generated by serially diluting purified Fc-PRL-13 into PBS. Prism (GraphPad Prism) was used to fit a sigmoidal curve to the data and used to quantify Fc-PRL-13 in the serum samples. Reported Fc-PRL-13 concentrations is represented as mean +/- SEM.

The serum concentration of His-tagged PRL N59D was determined using a quantitative ELISA as follows: 50ul of 0.1ug/mL anti-human PRL antibodies (Abcam; ab244037) were diluted in PBS and used to coat MaxiSorp 96-well ELISA plates for 1hr at room temperature. Plates were washed 3 times in PBST (PBS with 0.05% Tween 20, and then the plates were blocked with 200uL of blocking buffer (PBS with 3% BSA) overnight at 4C. Plates were again washed 3 times in PBST. 50uL of serum serially diluted in PBS was added to the plates and incubated at room temperature for 1hr. Plates were then washed 5 times in PBST. 50uL of HRP-conjugated anti-6xHis (Abcam; 1187) diluted 1:5000 in blocking buffer was added to the plates and incubated at room temperature for 1hr. Plates were then washed 5 times with PBST. 50uL of TMB Substrate Solution (Thermo Fisher Scientific; 34028) equilibrated to room temperature was added to the plates and incubated for 7 minutes or until desired color develops. Reactions were stopped by adding 50uL of 2M Sulfuric Acid to each well. The absorbance was measured at 450nm. A standard curve was generated by serially diluting purified PRL N59D into PBS. Prism (GraphPad Prism) was used to fit a sigmoidal curve to the data and used to quantify PRL N59D in the serum samples. Reported PRL N59D concentrations is represented as mean +/- SEM.

For Fc-PRL fusions, the half-life of the distribution and elimination phases were calculated by fitting two-phase decay curves to the data in PRISM (GraphPad Prism). A one-phase decay was fit to the PRL N59D data in PRISM and used to calculate the elimination half-life. AUC_inf_ and AUMC_inf_ was also calculated using PRISM. Serum clearance was calculated as:

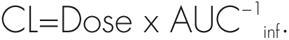

Volumes of distribution at steady stat were estimated as:

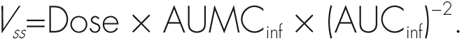

### Pharmacokinetic studies in lactating mice

All animal work was approved by the Harvard Medical School IACUC under protocol IS00003310. C57bl/6j mice (Jackson; 000664) were mated at 8 weeks of age. On day 7 post-partum, litters were normalized to 5 pups per dam. Pups were weighed daily until they were weaned at day 21 post-partum or they reached endpoint (20% weight loss compared to positive controls). The weight of the pups is reported as mean +/- sem. PRISM (GraphPad Prism) was used to perform multiple unpaired t-tests or two-way ANOVA to compare pup weights.

Additionally, on day 7 post-partum dams were administered either a 5mg/kg dose of Fc-PRL-13 or a PBS control either by intravenous injection into the tail vein or by subcutaneous injection. Blood samples were collected from the tail vein at various time points. Serum concentrations of Fc-PRL-13 and pharmacokinetic parameters were determined as previously described.

### Bromocriptine-induced lactation insufficiency studies in mice

All animal work was approved by the Harvard Medical School IACUC under protocol IS00003310. The experiment was modified from elsewhere^81^. C57bl/6j mice (Jackson; 000664) were mated at 8 weeks of age. On day 7 post-partum, litters were normalized to 5 pups per dam. Pups were weighed daily until they were weaned at day 21 post-partum or they reached endpoint (20% weight loss compared to positive controls). The weight of the pups is reported as mean +/- sem. PRISM (GraphPad Prism) was used to perform two-way ANOVA with Bonferroni correction to compare the weight of the pups and to perform Kaplan-Meier analysis of the pup’s survival. The survival proportions are represented as a percentage +/- SE.

On day 7 post-partum dams were administered either a 0.05mg/kg, 0.5mg/kg, or 5mg/kg dose of Fc-PRL-13, 5mg/kg dose of PRL N59D, or a PBS control by subcutaneous injection once on day 7 or every other dat. Additionally, beginning on day 7 post-partum dams were administered twice-daily a 200ug dose of Bromocriptine (2mg/mL bromocriptine mesylate (Sigma; 1076501) and 2mg/mL tartaric acid (Sigma; PHR1472) dissolved in PBS and 20% ethanol) or100uL of vehicle (2mg/mL tartaric acid (Sigma; PHR1472) dissolved in PBS with 20% ethanol). Dams were dosed until all their pups reached endpoint or were weaned at day 21 post-partum.

### Increasing lactation in mice

All animal work was approved by the Harvard Medical School IACUC under protocol IS00003310. C57bl/6j mice (Jackson; 000664) were mated at 8 weeks of age. On day 7 post-partum, litters were normalized to 5 pups per dam. Pups were weighed daily until they were weaned at day 21 post-partum or they reached endpoint (20% weight loss compared to positive controls). Beginning on day 7 post-partum, and every other day thereafter, dams were administered 100ug dose of Fc-PRL-13 or a PBS control by subcutaneous. Dams were dosed until their pups were weaned at day 21 post-partum. The weight of the pups is reported as mean +/- sem. PRISM (GraphPad Prism) was used to perform multiple unpaired t-tests with Bonferroni correction to compare the weight of the pups.

### Mammary gland harvesting in lactating mice

All animal work was approved by the Harvard Medical School IACUC under protocol IS00003310. C57bl/6j mice (Jackson; 000664) were mated at 8 weeks of age. On day 7 post-partum, litters were normalized to 5 pups per dam. Pups were weighed daily until their dams were sacrificed for mammary gland harvesting, at which point the pups were also sacrificed. Beginning on day 7 post-partum, and every other day thereafter, dams were administered a5mg/kg dose of Fc-PRL-13 or a PBS control by subcutaneous injection. Dams were dosed until they were sacrificed for mammary gland harvesting at various time points between day 7 post-partum and day 21 post-partum (n=3 per time point). Blood was collected from the dams via cardiac puncture, and then cardiac perfusion was performed.

After cardiac perfusion, the mammary glands were harvested. The inguinal mammary glands from one flank were harvested for whole carmine staining and the inguinal mammary glands from the other side were used to generate FFPE sections. The abdominal mammary gland from one side was harvested, stored in RNALater (Thermo Fisher, AM702) for 24h at 4C, and stored at −80C until further study. The proteins from the other abdominal mammary gland from was extracted as follows: The abdominal mammary glands were weighed and homogenized in RIPA Lysis buffer supplemented with cOmplete protease inhibitor cocktail (Sigma; 11836145001) and Phosphatase inhibitor cocktail (Thermo Fisher; A32957) according to the manufacturer’s instructions. The amount of protein in the supernatant was quantified via BCA (Thermo Fisher; 23227) according to the manufacturer’s instructions. Aliquots were stored at −20C until further use.

### H&E staining

Inguinal mammary glands were fixed in 10% neutral-buffered formalin for 24h. Hematoxylin and eosin (H&E) slides were prepared by the Rodent Histopathology core at Harvard Medical School. Dr. Roderick Bronson, a rodent histopathology from the Rodent Histopathology core at Harvard Medical School, reviewed all H&E slides. Imaging of the slides was provided by the Neurobiology Imaging Facility (NIF) at Harvard Medical School, Boston, MA.

### Carmine staining

Whole mount carmine-stained mammary glands were prepared as described elsewhere^82^. Imaging of the slides was provided by the Neurobiology Imaging Facility (NIF) at Harvard Medical School, Boston, MA. Qupath was used to measure the area of the mammary glands.

### Immunohistochemistry

Immunohistochemistry was performed by HistoWiz Inc. (histowiz.com) using a Standard Operating Procedure and fully automated workflow. Samples were processed, embedded in paraffin, and sectioned at 4µm. Immunohistochemistry was performed on a Bond Rx autostainer (Leica Biosystems) with enzyme treatment (1:1000) using standard protocols.

Antibodies used were rat monoclonal F4/80 primary antibody (eBioscience, 14-4801, 1:200) and rabbit anti-rat secondary (Vector, 1:100). Bond Polymer Refine Detection (Leica Biosystems) was used according to the manufacturer’s protocol. After staining, sections were dehydrated and film coverslipped using a TissueTek-Prisma and Coverslipper (Sakura). Whole slide scanning (40x) was performed on an Aperio AT2 (Leica Biosystems). Qupath was used to measure overall DAB staining, annotate the area of the mammary gland, and to perform cell segmentation. For each cell, Qupath measures a variety of morphology parameters including nuclear area, perimeter, and max calipers. Downstream analysis of the segmented cells was done in Python.

### ELISA assay of Fc-PRL in mouse mammary glands

The concentrations of Fc-PRL-13 in the abdominal mammary glands were determined using a quantitative ELISA as previously described. Anti-human PRL antibodies (Abcam; ab244037) were coated onto MaxiSorp 96-well ELISA plates to capture Fc-PRL-13 from serially diluted mammary gland protein extracts. HRP-conjugated anti-human IgG Fc antibodies (Invivogen; 31413) were used for detection. A standard curve was generated by serially diluting purified Fc-PRL-13 into PBS. Prism (GraphPad) was used to fit a sigmoidal curve to the data and used to quantify Fc-PRL-13 in the protein extract samples. Reported Fc-PRL-13 concentrations is represented as mean +/- SEM of biological triplicates.

### Western blot analysis of lactation biomarkers

20ug of abdominal mammary gland protein extracts were combined with equal parts (v:v) of 2x laemmli BME buffer (Bio Rad 1610737), boiled for 10mins at 100C, and separated on 4-20% Tris-glycine gels (Thermo Fisher; XP04205BOX) in Tris-glycine SDS running buffer (Thermo Fisher; LC2675). The protein was then transferred onto nitrocellulose membrane (Thermo Fisher; IB23002) and blocker for 1h in TBST with 1% BSA. The primary antibodies were diluted in blocking buffer at 1:1000 anti-prolactin receptor (Abcam; ab214303), 1:1000 anti-β-casein (Thermo Fisher, MA1-46056), 1:1000 anti-STAT5 (Cell Signaling Technologies; 94205), 1:1000 anti-pSTAT5 (Cell Signaling Technologies; 9351S), 1:1000 anti-STAT3 (Cell Signaling Technologies; 12640), 1:1000 anti-pSTAT3 (Cell Signaling Technologies; 9145), and 1:10,000 anti-β-actin (Cell Signaling Technologies; 3700S).

Secondary antibodies were diluted in blocking buffer at 1:10,000 anti-mouse-HRP (Thermo Fisher; 31340) and 1:10,000 anti-rabbit-HRP (Thermo Fisher; 31460). Chemiluminescence was detected using SuperSignal West Dura Extended Duration Substrate (Thermo Fisher; 34075). The densitometry of the western blots was analyzed by the Image Lab software. The data is reported as mean +/- SEM of biological triplicates.

### Pharmacokinetics studies in pups

All animal work was approved by the Harvard Medical School IACUC under protocol IS00003310. C57bl/6j mice (Jackson; 000664) were mated at 8 weeks of age. On day 7 post-partum, litters were normalized to 5 pups per dam. Beginning on day 7 post-partum, and every other day thereafter, dams were administered a 5mg/kg dose of Fc-PRL-13 or a PBS control by subcutaneous injection (n=3 per time point). Dams were dosed until their pups were sacrificed. Pups were weighed daily until they were sacrificed by CO2 asphyxiation and decapitation at various timepoints for blood collection (n=15 per time point). Serum concentrations of Fc-PRL-13 was determined by a quantitative ELISA as previously described. Reported Fc-PRL-13 concentrations is represented as mean +/- SEM.

