## Supplemental Information for "A long-acting prolactin to combat lactation insufficiency"

Human Prolactin WT (P01236 29-227)
LPICPGGAARCQVTLRDLFDRAVVLSHYIHNLSSEMFSEFDKRYTHGRGFITKAINSCHTSSLATPEDKEQAQQMNQKDFL
SLIVSILRSWNEPLYHLVTEVRGMQEAPEAILSKAVEIEEQTKRLLEGMELIVSQVHPETKENEIYPVWSGLPSLQMADEESRLS
AYYNLLHCLRRDSHKIDNYLKLLKCRIIHNNNCEQKLISEEDLNSAVDHHHHHH

C-myc tag + 6xHis tag: EQKLISEEDLNSAVDHHHHHH

Human Prolactin N59D
LPICPGGAARCQVTLRDLFDRAVVLSHYIHDLSSEMFSEFDKRYTHGRGFITKAINSCHTSSLATPEDKEQAQQMNQKDFLS
LIVSILRSWNEPLYHLVTEVRGMQEAPEAILSKAVEIEEQTKRLLEGMELIVSQVHPETKENEIYPVWSGLPSLQMADEESRLSA
YYNLLHCLRRDSHKIDNYLKLLKCRIIHNNNCEQKLISEEDLNSAVDHHHHHH

C-myc tag + 6xHis tag: EQKLISEEDLNSAVDHHHHHH

Human Prolactin Receptor (P16471)
DYKDDDDKQLPPGKPEIFKCRSPNKETFTCWWRPGTDGGLPTNYSLTYHREGETLMHECPDYITGGPNSCHFGKQYTS
MWRTYIMMVNATNQMGSSFSDELYVDVTYIVQPDPPLELAVEVKQPEDRKPYLWIKWSPPTLIDLKTGWFTLLYEIRLKPEKA
AEWEIHFAGQQTEFKILSLHPGQKYLVQVRCKPDHGYWSAWSPATFIQIPSDFTMNDTTVWISVAVLSAVICLIIVWAVA
LKGYSMVTCIFPPVPGPKIKGFDAHLLEKGKSEELLSALGCQDFPPTSDYEDLLVEYLEVDDSEDQHLMSVHSKEHPSQGMK
PTYLDPDTDSGRGSCDSPSLLSEKCEEPQANPSTFYDPEVIEKPENPETTHTWDPQCISMEGKIPYFHAGGSKCSTWPLPQ
PSQHNPRSSYHNITDVCELAVGPAGAPATLLNEAGKDALKSSQTIKSREEGKATQQREVESFHSETDQDTPWLLPQEKTPF
GSAKPLDYVEIHKVNKDGALSLLPKQRENSGKPKKPGTPENNKEYAKVSGVMDNNILVLVPDPHAKNVACFEESAKEAPP
SLEQNQAEKALANFTATSSKCRLQLGGLDYLDPACFTHSFH

Flag Tag: DYKDDDDK

Mouse Prolactin (P06879)
LPICSAGDCQTSLRELFDRVVILSHYIHTLYTDMFIEFDKQYVQDREFMVKVINDCPTSSLATPEDKEQALKVPPEVLLNLILSLVQ
SSSDPLFQLITGVGGIQEAPEYILSRAKEIEEQNKQLLEGVEKIISQAYPEAKGNGIYFVWSQLPSLQGVDEESKILSLRNTIRC
LRRDSHKVDNFLKVLRCQIAHQNNCEQKLISEEDLNSAVDHHHHHH

C-myc tag + 6xHis tag: EQKLISEEDLNSAVDHHHHHH

Mouse Prolactin Receptor (Q08501)
DYKDDDDKQSPPGKPEIHKCRSPDKETFTCWWNPGSDGGLPTNYSLTYSKEGEKNTYECPDYKTSGPNSCFFSKQYTS
IWKIYIITVNATNEMGSSTSDPLYVDVTYIVEPEPPRNLTLEVKQLKDKKTYLWVKWLPPTITDVKTGWFTMEYEIRLKSEEADE
WEIHFTGHQTQFKVFDLYPGQKYLVQTRCKPDHGYWSRWGQEKSIEIPNDFTLKDTTVWIIVAVLSAVICLIMVWAVALK
GYSMMTCIFPPVPGPKIKGFDTHLLEKGKSEELLSALGCQDFPPTSDCEDLLVEFLEVDDNEDERLMPSHSKEYPGQGVKPT
HLDPDSDSGHGSYDSHSLLSEKCEEPQAYPPAFHIPEITEKPENPEANIPPTPNPQNNTPNCHTDTSKSTTWPLPPGQHTR
RSPYHSIADVCKLAGSPGDTLDSFLDKAEENVLKLSEDAGEEEVAVQEGAKSFPSDKQNTSWPPLQEKGPIVYAKPPDYVEI
HKVNKDGVLSLLPKQRENHQTENPGVPETSKEYAKVSGVTDNNILVLVPDSRAQNTALLEESAKKVPPSLEQNQSEKDLAS
FTATSSNCRLQLGRLDYLDPTCFMHSFH

Flag Tag: DYKDDDDK

Human IgG1 Fc WT
EPKSCDKTHTCPPCPAPELLGGPSVFLFPPKPKDTLMISRTPEVTCVVVDVSHEDPEVKFNWYVDGVEVHNAKTKPREEQYN
STYRVVSVLTVLHQDWLNGKEYKCKVSNKALPAPIEKTISKAKGQPREPQVYTLPPSRDELTKNQVSLTCLVKGFYPSDIAVE
WESNGQPENNYKTTPPVLDSDGSFFLYSKLTVDKSRWQQGNVFSCSVMHEALHNHYTQKSLSLSPGK

Human FcγRI (P12314 16-292)
QVDTTKAVITLQPPWVSVFQEETVTLHCEVLHLPGSSSTQWFLNGTATQTSTPSYRITSASVNDSGEYRCQRGLSGRSDPI
QLEIHRGWLLLQVSSRVFTEGEPLALRCHAWKDKLVYNVLYYRNGKAFKFFHWNSNLTILKTNISHNGTYHCSGMGKHR
YTSAGISVTVKELFPAPVLNASVTSPLLEGNLVTLSCETKLLLQRPGLQLYFSFYMGSKTLRGRNTSSEYQILTARREDSGLYWCE
AATEDGNVLKRSPELELQVLGLQLPTPVWFH EQKLISEEDLNSAVDHHHHHH

C-myc tag + 6xHis tag: EQKLISEEDLNSAVDHHHHHH

Human FcγRIIa (P12318 34-217)
QAAAPPKAVLKLEPPWINVLQEDSVTLTCQGARSPESDSIQWFHNGNLIPTHTQPSYRFKANNNDSGEYTCQTGQTSL
SDPVHLTVLSEWLVLQTPHLEFQEGETIMLRCHSWKDKPLVKVTFFQNGKSQKFSHLDPTFSIPQANHSHSGDYHCTGNI
GYTLFSSKPVTITVQVPSMGSSSPMG EQKLISEEDLNSAVDHHHHHH

C-myc tag + 6xHis tag: EQKLISEEDLNSAVDHHHHHH

Human FcγRIIb (P31994 43-217)
TPAAPPKAVLKLEPQWINVLQEDSVTLTCRGTHSPESDSIQWFHNGNLIPTHTQPSYRFKANNNDSGEYTCQTGQTSLS
DPVHLTVLSEWLVLQTPHLEFQEGETIVLRCHSWKDKPLVKVTFFQNGKSKKFSRSDPNFSIPQANHSHSGDYHCTGNIG
YTLYSSKPVTITVQAP EQKLISEEDLNSAVDHHHHHH

C-myc tag + 6xHis tag: EQKLISEEDLNSAVDHHHHHH

Human FcγRIIIa (P08637 17-208)
GMRTEDLPKAVVFLEPQWYRVLEKDSVTLKCQGAYSPEDNSTQWFHNESLISSQASSYFIDAATVDDSGEYRCQTNLSTL
SDPVQLEVHIGWLLLQAPRWVFKEEDPIHLRCHSWKNTALHKVTYLQNGKGRKYFHHNSDFYIPKATLKDSGSYFCRGLF
GSKNVSSETVNITITQGLAVSTISSFFPPGYQEQKLISEEDLNSAVDHHHHHH

C-myc tag + 6xHis tag: EQKLISEEDLNSAVDHHHHHH

Mouse FcγRI (P26151 25-297)
EVVNATKAVITLQPPWVSIFQKENVTLWCEGPHLPGDSSTQWFINGTAVQISTPSYSIPEASFQDSGEYRCQIGSSMPS
DPVQLQIHNDWLLLQASRRVLTEGEPLALRCHGWKNKLVYNVVFYRNGKSFQFSSDSEVAILKTNLSHSGIYHCSGTGR
HRYTSAGVSITVKELFTTPVLRASVSSPFPEGSLVTLNCETNLLLQRPGLQLHFSFYVGSKILEYRNTSSEYHIARAEREDAGFYW
CEVATEDSSVLKRSPELELQVLGPQSSAPEQKLISEEDLNSAVDHHHHHH

C-myc tag + 6xHis tag: EQKLISEEDLNSAVDHHHHHH

Mouse FcγRIIb (P08101 30-210)
THDLPKAVVKLEPPWIQVLKEDTVTLTCEGTHNPGNSSTQWFHNGRSIRSQVQASYTFKATVNDSGEYRCQMEQTRLS
DPVDLGVISDWLLLQTPQLVFLEGETITLRCHSWRNKLLNRISFFHNEKSVRYHHYSSNFSIPKANHSHSGDYYCKGSLGRT
LHQSKPVTITVQGPKSSRSLPEQKLISEEDLNSAVDHHHHHH

C-myc tag + 6xHis tag: EQKLISEEDLNSAVDHHHHHH

Mouse FcγRIII (P08508 31-215)
ALPKAVVKLDPPWIQVLKEDMVTLMCEGTHNPGNSSTQWFHNGRSIRSQVQASYTFKATVNDSGEYRCQMEQTRLS
DPVDLGVISDWLLLQTPQRVFLEGETITLRCHSWRNKLLNRISFFHNEKSVRYHHYKSNFSIPKANHSHSGDYYCKGSLGST
QHQSKPVTITVQDPATTSSISLVWYHTEQKLISEEDLNSAVDHHHHHH

C-myc tag + 6xHis tag: EQKLISEEDLNSAVDHHHHHH

Mouse FcγRIV (A0A0B4J1G0 21-203)
GLQKAVVNLDPKWVRVLEEDSVTLRCQGTFSPEDNSIKWFHNESLIPHQDANYVIQSARVKDSGMYRCQTALSTISDPV
QLEVHMGWLLLQTTKWLFQEGDPIHLRCHSWQNRPVRKVTYLQNGKGKKYFHENSELLIPKATHNDSGSYFCRGLIGH
NNKSSASFRISLGDPGSPSMFPPWH EQKLISEEDLNSAVDHHHHHH

C-myc tag + 6xHis tag: EQKLISEEDLNSAVDHHHHHH

Table S1: Fc-prolactin fusion variants

| Fc-PRL- # | Variants |
|---|---|
| 1 | **Fc** - **PRL** (N59D) |
| 2. | **Fc** (C220S, N297D, K427A) - **PRL** WT |
| 3. | **Fc** (C220S, N297D, K427A) - **PRL** (N59D) |
| 4. | **Fc** (C220S, N297D, K427A) - **GGsGG** - **PRL** (N59D) |
| 5. | **Fc** (C220S, K447A, N297D, L234A, L235A, V264E, L309D, Q311H, N434S) - **PRL** (N59D) |
| 6. | **Fc** (C220S, K447A, N297D, L234A, L235A, P329G, V264E, L309D, Q311H, N434S) - **PRL** (N59D) |
| 7. | **Fc** (C220S, K447A, N297D, L234A, L235A, M252Y, S254T, T256E) - **PRL** (N59D) |
| 8. | **Fc Knob** (C220S, N297D, K427A, T366W) - **PRL** (N59D) / **Fc Hole** (C220S, N297D, K427A, T366S, L368A, Y407V) |
| 9. | **Fc Knob** (C220S, N297D, K427A, T366W) - **GGsGG** - **PRL** (N59D) / **Fc Hole** (C220S, N297D, K427A, T366S, L368A, Y407V) |
| 10. | **Fc Knob** (C220S, K447A, N297D, T366W, L234A, L235A, V264E, L309D, Q311H, N434S) - **GGsGG** - **PRL** (N59D) / **Fc Hole** (C220S, N297D, K447A, T366S, L368A, Y407V, H435R, Y436F, L234A, L235A, V264E, L309D, Q311H, N434S) |
| 11. | **Fc Knob** (C220S, K447A, N297D, T366W, L234A, L235A, P329G, V264E, L309D, Q311H, N434S) - **GGsGG** - **PRL** (N59D) / **Fc Hole** (C220S, N297D, K447A, T366S, L368A, Y407V, H435R, Y436F, L234A, L235A, P329G, V264E, L309D, Q311H, N434S) |
| 12. | **Fc Knob** (C220S, K447A, N297D, L234A, L235A, M252Y, S254T, T256E) - **GGsGG** - **PRL** (N59D) / **Fc Hole** (C220S, N297D, K447A, T366S, L368A, Y407V, H435R, Y436F, L234A, L235A, M252Y, S254T, T256E) |
| 13. | **Fc A** (C220S, K447A, N297D, T350V, L351Y, F405A, Y407V) - **GGsGG** - **PRL** (N59D) / **Fc B** (C220S, N297D, K427A, T350V, T366L, K293L, T394W, H435R, Y436F) |
| 14. | **Fc A** (C220S, K447A, N297D, T350V, L351Y, F405A, Y407V, L234A, L235A, V264E, L309D, Q311H, N434S) - **GGsGG** - **PRL** (N59D) / **Fc B** (N297D, C220S, K447A, L234A, L235A, V264E, L309D, Q311H, N434S) |
| 15. | **Fc A** (C220S, K447A, N297D, T350V, L351Y, F405A, Y407V, L234A, L235A, V264E, L309D, Q311H, N434S) - **GGsGG** - **PRL** (N59D) / **Fc B** (N297D, C220S, K447A, L234A, L235A, V264E, L309D, Q311H, N434S, H435R, Y436F) |
| 16. | **Fc A** (C220S, K447A, N297D, T350V, L351Y, F405A, Y407V, L234A, L235A, P329G, V264E, L309D, Q311H, N434S) - **GGsGG** - **PRL** (N59D) / **Fc B** (N297D, C220S, K447A, L234A, L235A, P329G, V264E, L309D, Q311H, N434S, H435R, Y436F) |
| 17. | **Fc A** (C220S, K447A, N297D, T350V, L351Y, F405A, Y407V, L234A, L235A, M252Y, S254T, T256E) - **GGsGG** - **PRL** (N59D) / **Fc B** (N297D, K447A, H435R, Y436F, L234A, L235A, M252Y, S254T, T256E) |
| 18. | **PRL** (N59D) - **Fc Knob** (C220S, N297D, K427A, T366W) / **Fc Hole** (C220S, N297D, K427A, T366S, L368A, Y407V) |
| 19. | **PRL** (N59D) - **GGsGG** - **Fc Knob** (C220S, N297D, K427A, T366W) / **Fc Hole** (C220S, N297D, K427A, T366S, L368A, Y407V) |
| 20. | **PRL** (N59D, C191S, C199S) - **Fc Knob** (C220S, N297D, K427A, T366W) / **Fc Hole** (C220S, N297D, K427A, T366S, L368A, Y407V) |

| | |
|---|---|
| 21. | **PRL** (N59D, C191S, C199S) - **Fc Knob** (C220S, C226S, C229S, N297D, K427A, T366W) / **Fc Hole** (C220S, N297D, K427A, T366S, L368A, Y407V) |
| 22. | **PRL** (N59D) - **GGsGG**- **Fc Knob** (C220S, K447A, N297D, T366W, L234A, L235A, V264E, L309D, Q311H, N434S) / **Fc Hole** (C220S, N297D, K447A, T366S, L368A, Y407V, H435R, Y436F, L234A, L235A, V264E, L309D, Q311H, N434S) |
| 23. | **PRL** (N59D) - **GGsGG**- **Fc Knob** (C220S, K447A, N297D, T366W, L234A, L235A, P329G, V264E, L309D, Q311H, N434S) / **Fc Hole** (C220S, N297D, K447A, T366S, L368A, Y407V, H435R, Y436F, L234A, L235A, P329G, V264E, L309D, Q311H, N434S) |
| 24. | **PRL** (N59D) - **GGsGG** - **Fc Knob** (C220S, K447A, N297D, L234A, L235A, M252Y, S254T, T256E) / **Fc Hole** (C220S, N297D, K447A, T366S, L368A, Y407V, H435R, Y436F, L234A, L235A, M252Y, S254T, T256E) |
| 25. | **PRL** (N59D) - **GGsGG** - **Fc A** (C220S, K447A, N297D, T350V, L351Y, F405A, Y407V) / **Fc B** (C220S, N297D, K427A, T350V, T366L, K293L, T394W) |
| 26. | **PRL** (N59D) - **GGsGG**- **Fc A** (C220S, K447A, N297D, T350V, L351Y, F405A, Y407V, L234A, L235A, V264E, L309D, Q311H, N434S) / **Fc A** (N297D, C220S, K447A, L234A, L235A, V264E, L309D, Q311H, N434S) |
| 27. | **PRL** (N59D) - **GGsGG**- **Fc A** (C220S, K447A, N297D, T350V, L351Y, F405A, Y407V, L234A, L235A, V264E, L309D, Q311H, N434S) / **Fc A** (N297D, C220S, K447A, L234A, L235A, V264E, L309D, Q311H, N434S, H435R, Y436F) |
| 28. | **PRL** (N59D) - **GGsGG**- **Fc A** (C220S, K447A, N297D, T350V, L351Y, F405A, Y407V, L234A, L235A, P329G, V264E, L309D, Q311H, N434S) / **Fc B** (N297D, C220S, K447A, L234A, L235A, P329G, V264E, L309D, Q311H, N434S, H435R, Y436F) |
| 29. | **PRL** (N59D) - **GGsGG** - **Fc A** (C220S, K447A, N297D, T350V, L351Y, F405A, Y407V, L234A, L235A, M252Y, S254T, T256E) / **Fc B** (N297D, K447A, H435R, Y436F, L234A, L235A, M252Y, S254T, T256E) |

Table S2: Amino acid sequences of Fc-prolactin fusions

| Fc-PRL-# | Sequence |
|---|---|
| 1 | Fc - PRL (N59D):<br>EPKSCDKTHTCPPCPAPELLGGPSVFLFPPKPKDTLMISRTPEVTCVVVDVSHEDPEVKFNWYVDGVEVHNA<br>KTKPREEQYNSTYRVVSVLTVLHQDWLNGKEYKCKVSNKALPAPIEKTISKAKGQPREPQVYTLPPSRDELTK<br>NQVSLTCLVKGFYPSDIAVEWESNGQPENNYKTTPPVLDSDGSFFLYSKLTVDKSRWQQGNVFSCSV<br>MHEALHNHYTQKSLSLSPGKLPICPGGAARCQVTLRDLFDRAVVLSHYIHDLSSEMFSEFDKRYTHGRGFIT<br>KAINSCHTSSLATPEDKEQAQQMNQKDFLSLIVSILRSWNEPLYHLVTEVRGMQEAPEAILSKAVEIEEQTK<br>RLLEGMELIVSQVHPETKENEIYPVWSGLPSLQMADEESRLSAYYNLLHCLRRDSHKIDNYLKLLKCRIIHNN<br>NC |
| 2. | Fc (C220S, N297D, K427A) - PRL WT:<br>EPKSsDKTHTCPPCPAPELLGGPSVFLFPPKPKDTLMISRTPEVTCVVVDVSHEDPEVKFNWYVDGVEVHNA<br>KTKPREEQYdSTYRVVSVLTVLHQDWLNGKEYKCKVSNKALPAPIEKTISKAKGQPREPQVYTLPPSRDELTK<br>NQVSLTCLVKGFYPSDIAVEWESNGQPENNYKTTPPVLDSDGSFFLYSKLTVDKSRWQQGNVFSCSV<br>MHEALHNHYTQKSLSLSPGaLPICPGGAARCQVTLRDLFDRAVVLSHYIHNLSSEMFSEFDKRYTHGRGFIT<br>KAINSCHTSSLATPEDKEQAQQMNQKDFLSLIVSILRSWNEPLYHLVTEVRGMQEAPEAILSKAVEIEEQTK<br>RLLEGMELIVSQVHPETKENEIYPVWSGLPSLQMADEESRLSAYYNLLHCLRRDSHKIDNYLKLLKCRIIHNN<br>NCEQKLISEEDLNSAVDHHHHHH |
| 3. | Fc (C220S, N297D, K427A) - PRL (N59D):<br>EPKSsDKTHTCPPCPAPELLGGPSVFLFPPKPKDTLMISRTPEVTCVVVDVSHEDPEVKFNWYVDGVEVHNA<br>KTKPREEQYdSTYRVVSVLTVLHQDWLNGKEYKCKVSNKALPAPIEKTISKAKGQPREPQVYTLPPSRDELTK<br>NQVSLTCLVKGFYPSDIAVEWESNGQPENNYKTTPPVLDSDGSFFLYSKLTVDKSRWQQGNVFSCSV<br>MHEALHNHYTQKSLSLSPGaLPICPGGAARCQVTLRDLFDRAVVLSHYIHDLSSEMFSEFDKRYTHGRGFIT<br>KAINSCHTSSLATPEDKEQAQQMNQKDFLSLIVSILRSWNEPLYHLVTEVRGMQEAPEAILSKAVEIEEQTK<br>RLLEGMELIVSQVHPETKENEIYPVWSGLPSLQMADEESRLSAYYNLLHCLRRDSHKIDNYLKLLKCRIIHNN<br>NC |
| 4. | Fc (C220S, N297D, K427A) - GGsGG - PRL (N59D):<br>EPKSsDKTHTCPPCPAPELLGGPSVFLFPPKPKDTLMISRTPEVTCVVVDVSHEDPEVKFNWYVDGVEVHNA<br>KTKPREEQYdSTYRVVSVLTVLHQDWLNGKEYKCKVSNKALPAPIEKTISKAKGQPREPQVYTLPPSRDELTK<br>NQVSLTCLVKGFYPSDIAVEWESNGQPENNYKTTPPVLDSDGSFFLYSKLTVDKSRWQQGNVFSCSV<br>MHEALHNHYTQKSLSLSPGaGGsGGLPICPGGAARCQVTLRDLFDRAVVLSHYIHDLSSEMFSEFDKRYT<br>HGRGFITKAINSCHTSSLATPEDKEQAQQMNQKDFLSLIVSILRSWNEPLYHLVTEVRGMQEAPEAILSKA<br>VEIEEQTKRLLEGMELIVSQVHPETKENEIYPVWSGLPSLQMADEESRLSAYYNLLHCLRRDSHKIDNYLKLLK<br>CRIIHNNNCEQKLISEEDLNSAVDHHHHHH |
| 5. | Fc (C220S, K447A, N297D, L234A, L235A, V264E, L309D, Q311H, N434S) - PRL<br>(N59D):<br>EPKSsDKTHTCPPCPAPEAAGGPSVFLFPPKPKDTLMISRTPEVTCVVEDVSHEDPEVKFNWYVDGVEVHN<br>AKTKPREEQYdSTYRVVSVLTVDHHDWLNGKEYKCKVSNKALPAPIEKTISKAKGQPREPQVYTLPPSRDELT<br>KNQVSLTCLVKGFYPSDIAVEWESNGQPENNYKTTPPVLDSDGSFFLYSKLTVDKSRWQQGNVFSCSV<br>MHEALHSHYTQKSLSLSPGaLPICPGGAARCQVTLRDLFDRAVVLSHYIHDLSSEMFSEFDKRYTHGRGFITK<br>AINSCHTSSLATPEDKEQAQQMNQKDFLSLIVSILRSWNEPLYHLVTEVRGMQEAPEAILSKAVEIEEQTKR<br>LLEGMELIVSQVHPETKENEIYPVWSGLPSLQMADEESRLSAYYNLLHCLRRDSHKIDNYLKLLKCRIIHNNN<br>C |
| 6. | Fc (C220S, K447A, N297D, L234A, L235A, P329G, V264E, L309D, Q311H, N434S) -<br>PRL N59D:<br>EPKSsDKTHTCPPCPAPEAAGGPSVFLFPPKPKDTLMISRTPEVTCVVEDVSHEDPEVKFNWYVDGVEVHN<br>AKTKPREEQYdSTYRVVSVLTVDHHDWLNGKEYKCKVSNKALGAPIEKTISKAKGQPREPQVYTLPPSRDEL<br>TKNQVSLTCLVKGFYPSDIAVEWESNGQPENNYKTTPPVLDSDGSFFLYSKLTVDKSRWQQGNVFSCS<br>VMHEALHSHYTQKSLSLSPGaLPICPGGAARCQVTLRDLFDRAVVLSHYIHDLSSEMFSEFDKRYTHGRGFIT<br>KAINSCHTSSLATPEDKEQAQQMNQKDFLSLIVSILRSWNEPLYHLVTEVRGMQEAPEAILSKAVEIEEQTK |

| | |
|---|---|
| | RLLEGMELIVSQVHPETKENEIYPVWSGLPSLQMADEESRLSAYYNLLHCLRRDSHKIDNYLKLLKCRIIHNN<br>NC |
| 7. | Fc (C220S, K447A, N297D, L234A, L235A, M252Y, S254T, T256E) - PRL N59D:<br>EPKSsDKTHTCPPCPAPEAAGGPSVFLFPPKPKDTLYITREPEVTCVVVDVSHEDPEVKFNWYVDGVEVHNA<br>KTKPREEQYdSTYRVVSVLTVLHQDWLNGKEYKCKVSNKALPAPIEKTISKAKGQPREPQVYTLPPSRDELTK<br>NQVSLTCLVKGFYPSDIAVEWESNGQPENNYKTTPPVLDSDGSFFLYSKLTVDKSRWQQGNVFSCSV<br>MHEALHNHYTQKSLSLSPGaLPICPGGAARCQVTLRDLFDRAVVLSHYIHDLSSEMFSEFDKRYTHGRGFIT<br>KAINSCHTSSLATPEDKEQAQQMNQKDFLSLIVSILRSWNEPLYHLVTEVRGMQEAPEAILSKAVEIEEQTK<br>RLLEGMELIVSQVHPETKENEIYPVWSGLPSLQMADEESRLSAYYNLLHCLRRDSHKIDNYLKLLKCRIIHNN<br>NC |
| 8. | Fc Knob (C220S, N297D, K427A, T366W) - PRL (N59D):<br>EPKSsDKTHTCPPCPAPELLGGPSVFLFPPKPKDTLMISRTPEVTCVVVDVSHEDPEVKFNWYVDGVEVHNA<br>KTKPREEQYdSTYRVVSVLTVLHQDWLNGKEYKCKVSNKALPAPIEKTISKAKGQPREPQVYTLPPSRDELTK<br>NQVSLWCLVKGFYPSDIAVEWESNGQPENNYKTTPPVLDSDGSFFLYSKLTVDKSRWQQGNVFSCSV<br>MHEALHNHYTQKSLSLSPGaLPICPGGAARCQVTLRDLFDRAVVLSHYIHDLSSEMFSEFDKRYTHGRGFIT<br>KAINSCHTSSLATPEDKEQAQQMNQKDFLSLIVSILRSWNEPLYHLVTEVRGMQEAPEAILSKAVEIEEQTK<br>RLLEGMELIVSQVHPETKENEIYPVWSGLPSLQMADEESRLSAYYNLLHCLRRDSHKIDNYLKLLKCRIIHNN<br>NCEQKLISEEDLNSAVDHHHHHH<br><br>Fc Hole (C220S, N297D, K427A, T366S, L368A, Y407V):<br>EPKSSDKTHTCPPCPAPELLGGPSVFLFPPKPKDTLMISRTPEVTCVVVDVSHEDPEVKFNWYVDGVEVHNA<br>KTKPREEQYDSTYRVVSVLTVLHQDWLNGKEYKCKVSNKALPAPIEKTISKAKGQPREPQVYTLPPSRDELTK<br>NQVSLSCAVKGFYPSDIAVEWESNGQPENNYKTTPPVLDSDGSFFLVSKLTVDKSRWQQGNVFSCSV<br>MHEALHNHYTQKSLSLSPGA |
| 9. | Fc Knob (C220S, N297D, K427A, T366W) - GGsGG - PRL (N59D):<br>EPKSsDKTHTCPPCPAPELLGGPSVFLFPPKPKDTLMISRTPEVTCVVVDVSHEDPEVKFNWYVDGVEVHNA<br>KTKPREEQYdSTYRVVSVLTVLHQDWLNGKEYKCKVSNKALPAPIEKTISKAKGQPREPQVYTLPPSRDELTK<br>NQVSLWCLVKGFYPSDIAVEWESNGQPENNYKTTPPVLDSDGSFFLYSKLTVDKSRWQQGNVFSCSV<br>MHEALHNHYTQKSLSLSPGaGGsGGLPICPGGAARCQVTLRDLFDRAVVLSHYIHDLSSEMFSEFDKRYT<br>HGRGFITKAINSCHTSSLATPEDKEQAQQMNQKDFLSLIVSILRSWNEPLYHLVTEVRGMQEAPEAILSKA<br>VEIEEQTKRLLEGMELIVSQVHPETKENEIYPVWSGLPSLQMADEESRLSAYYNLLHCLRRDSHKIDNYLKLLK<br>CRIIHNNNC<br><br>Fc Hole (C220S, N297D, K427A, T366S, L368A, Y407V):<br>EPKSSDKTHTCPPCPAPELLGGPSVFLFPPKPKDTLMISRTPEVTCVVVDVSHEDPEVKFNWYVDGVEVHNA<br>KTKPREEQYDSTYRVVSVLTVLHQDWLNGKEYKCKVSNKALPAPIEKTISKAKGQPREPQVYTLPPSRDELTK<br>NQVSLSCAVKGFYPSDIAVEWESNGQPENNYKTTPPVLDSDGSFFLVSKLTVDKSRWQQGNVFSCSV<br>MHEALHNRFTQKSLSLSPGA |
| 10. | Fc Knob (C220S, K447A, N297D, T366W, L234A, L235A, V264E, L309D, Q311H,<br>N434S) - GGsGG - PRL (N59D):<br>EPKSsDKTHTCPPCPAPEAAGGPSVFLFPPKPKDTLMISRTPEVTCVVEDVSHEDPEVKFNWYVDGVEVHN<br>AKTKPREEQYdSTYRVVSVLTVDHHDWLNGKEYKCKVSNKALPAPIEKTISKAKGQPREPQVYTLPPSRDELT<br>KNQVSLWCLVKGFYPSDIAVEWESNGQPENNYKTTPPVLDSDGSFFLYSKLTVDKSRWQQGNVFSCS<br>VMHEALHSHYTQKSLSLSPGaGGsGGLPICPGGAARCQVTLRDLFDRAVVLSHYIHDLSSEMFSEFDKRYT<br>HGRGFITKAINSCHTSSLATPEDKEQAQQMNQKDFLSLIVSILRSWNEPLYHLVTEVRGMQEAPEAILSKA<br>VEIEEQTKRLLEGMELIVSQVHPETKENEIYPVWSGLPSLQMADEESRLSAYYNLLHCLRRDSHKIDNYLKLLK<br>CRIIHNNNC<br><br>Fc Hole (C220S, N297D, K447A, T366S, L368A, Y407V, H435R, Y436F, L234A,<br>L235A, V264E, L309D, Q311H, N434S): |

| | |
|---|---|
| | EPKSSDKTHTCPPCPAPEAAGGPSVFLFPPKPKDTLMISRTPEVTCVVEDVSHEDPEVKFNWYVDGVEVHN<br>AKTKPREEQYDSTYRVVSVLTVDHHDWLNGKEYKCKVSNKALPAPIEKTISKAKGQPREPQVYTLPPSRDEL<br>TKNQVSLSCAVKGFYPSDIAVEWESNGQPENNYKTTPPVLDSDGSFFLVSKLTVDKSRWQQGNVFSC<br>SVMHEALHSRFTQKSLSLSPGA |
| 11. | Fc Knob (C220S, K447A, N297D, T366W, L234A, L235A, P329G, V264E, L309D, Q311H, N434S) - GGsGG - PRL N59D:<br>EPKSsDKTHTCPPCPAPEAAGGPSVFLFPPKPKDTLMISRTPEVTCVVEDVSHEDPEVKFNWYVDGVEVHN<br>AKTKPREEQYdSTYRVVSVLTVDHHDWLNGKEYKCKVSNKALGAPIEKTISKAKGQPREPQVYTLPPSRDEL<br>TKNQVSLWCLVKGFYPSDIAVEWESNGQPENNYKTTPPVLDSDGSFFLYSKLTVDKSRWQQGNVFSC<br>SVMHEALHSHYTQKSLSLSPGaGGsGGLPICPGGAARCQVTLRDLFDRAVVLSHYIHDLSSEMFSEFDKRY<br>THGRGFITKAINSCHTSSLATPEDKEQAQQMNQKDFLSLIVSILRSWNEPLYHLVTEVRGMQEAPEAILSK<br>AVEIEEQTKRLLEGMELIVSQVHPETKENEIYPVWSGLPSLQMADEESRLSAYYNLLHCLRRDSHKIDNYLKLL<br>KCRIIHNNNC<br><br>Fc Hole (C220S, N297D, K447A, T366S, L368A, Y407V, H435R, Y436F, L234A, L235A, P329G, V264E, L309D, Q311H, N434S):<br>EPKSSDKTHTCPPCPAPEAAGGPSVFLFPPKPKDTLMISRTPEVTCVVEDVSHEDPEVKFNWYVDGVEVHN<br>AKTKPREEQYDSTYRVVSVLTVDHHDWLNGKEYKCKVSNKALGAPIEKTISKAKGQPREPQVYTLPPSRDEL<br>TKNQVSLSCAVKGFYPSDIAVEWESNGQPENNYKTTPPVLDSDGSFFLVSKLTVDKSRWQQGNVFSC<br>SVMHEALHSRFTQKSLSLSPGA |
| 12. | Fc Knob (C220S, K447A, N297D, L234A, L235A, M252Y, S254T, T256E) - GGsGG - PRL N59D:<br>EPKSsDKTHTCPPCPAPEAAGGPSVFLFPPKPKDTLYITREPEVTCVVVDVSHEDPEVKFNWYVDGVEVHNA<br>KTKPREEQYdSTYRVVSVLTVLHQDWLNGKEYKCKVSNKALPAPIEKTISKAKGQPREPQVYTLPPSRDELTK<br>NQVSLWCLVKGFYPSDIAVEWESNGQPENNYKTTPPVLDSDGSFFLYSKLTVDKSRWQQGNVFSCSV<br>MHEALHNHYTQKSLSLSPGaGGsGGLPICPGGAARCQVTLRDLFDRAVVLSHYIHDLSSEMFSEFDKRYT<br>HGRGFITKAINSCHTSSLATPEDKEQAQQMNQKDFLSLIVSILRSWNEPLYHLVTEVRGMQEAPEAILSKA<br>VEIEEQTKRLLEGMELIVSQVHPETKENEIYPVWSGLPSLQMADEESRLSAYYNLLHCLRRDSHKIDNYLKLLK<br>CRIIHNNNC<br><br>Fc Hole (C220S, N297D, K447A, T366S, L368A, Y407V, H435R, Y436F, L234A, L235A, M252Y, S254T, T256E):<br>EPKSSDKTHTCPPCPAPEAAGGPSVFLFPPKPKDTLYITREPEVTCVVVDVSHEDPEVKFNWYVDGVEVHN<br>AKTKPREEQYDSTYRVVSVLTVLHQDWLNGKEYKCKVSNKALPAPIEKTISKAKGQPREPQVYTLPPSRDELT<br>KNQVSLSCAVKGFYPSDIAVEWESNGQPENNYKTTPPVLDSDGSFFLVSKLTVDKSRWQQGNVFSCS<br>VMHEALHNRFTQKSLSLSPGA |
| 13. | Fc A (C220S, K447A, N297D, T350V, L351Y, F405A, Y407V) - GGsGG - PRL (N59D):<br>EPKSsDKTHTCPPCPAPELLGGPSVFLFPPKPKDTLMISRTPEVTCVVVDVSHEDPEVKFNWYVDGVEVHNA<br>KTKPREEQYdSTYRVVSVLTVLHQDWLNGKEYKCKVSNKALPAPIEKTISKAKGQPREPQVYVYPPSRDELT<br>KNQVSLTCLVKGFYPSDIAVEWESNGQPENNYKTTPPVLDSDGSFALVSKLTVDKSRWQQGNVFSCSV<br>MHEALHNHYTQKSLSLSPGaGGsGGLPICPGGAARCQVTLRDLFDRAVVLSHYIHDLSSEMFSEFDKRYT<br>HGRGFITKAINSCHTSSLATPEDKEQAQQMNQKDFLSLIVSILRSWNEPLYHLVTEVRGMQEAPEAILSKA<br>VEIEEQTKRLLEGMELIVSQVHPETKENEIYPVWSGLPSLQMADEESRLSAYYNLLHCLRRDSHKIDNYLKLLK<br>CRIIHNNNC<br><br>Fc B (C220S, N297D, K427A, T350V, T366L, K293L, T394W, H435R, Y436F):<br>EPKSsDKTHTCPPCPAPELLGGPSVFLFPPKPKDTLMISRTPEVTCVVVDVSHEDPEVKFNWYVDGVEVHNA<br>KTKPREEQYdSTYRVVSVLTVLHQDWLNGKEYKCKVSNKALPAPIEKTISKAKGQPREPQVYVLPPSRDELTK<br>NQVSLLCLVKGFYPSDIAVEWESNGQPENNYLTWPPVLDSDGSFFLYSKLTVDKSRWQQGNVFSCSV<br>MHEALHNRFTQKSLSLSPGa |

| | |
|---|---|
| 14. | Fc A (C220S, K447A, N297D, T350V, L351Y, F405A, Y407V, L234A, L235A, V264E, L309D, Q311H, N434S) - GGsGG - PRL (N59D):<br>EPKSsDKTHTCPPCPAPEAAGGPSVFLFPPKPKDTLMISRTPEVTCVVEDVSHEDPEVKFNWYVDGVEVHN<br>AKTKPREEQYdSTYRVVSVLTVDHHDWLNGKEYKCKVSNKALPAPIEKTISKAKGQPREPQVYVYPPSRDEL<br>TKNQVSLTCLVKGFYPSDIAVEWESNGQPENNYKTTPPVLDSDGSFALVSKLTVDKSRWQQGNVFSCS<br>VMHEALHSHYTQKSLSLSPGaGGsGGLPICPGGAARCQVTLRDLFDRAVVLSHYIHDLSSEMFSEFDKRYT<br>HGRGFITKAINSCHTSSLATPEDKEQAQQMNQKDFLSLIVSILRSWNEPLYHLVTEVRGMQEAPEAILSKA<br>VEIEEQTKRLLEGMELIVSQVHPETKENEIYPVWSGLPSLQMADEESRLSAYYNLLHCLRRDSHKIDNYLKLLK<br>CRIIHNNNC<br><br>Fc A (N297D, C220S, K447A, L234A, L235A, V264E, L309D, Q311H, N434S):<br>EPKSsDKTHTCPPCPAPEAAGGPSVFLFPPKPKDTLMISRTPEVTCVVVDVSHEDPEVKFNWYVDGVEVHN<br>AKTKPREEQYdSTYRVVSVLTVDHHDWLNGKEYKCKVSNKALPAPIEKTISKAKGQPREPQVYVLPPSRDEL<br>TKNQVSLLCLVKGFYPSDIAVEWESNGQPENNYLTWPPVLDSDGSFFLYSKLTVDKSRWQQGNVFSCS<br>VMHEALHSHYTQKSLSLSPGa |
| 15. | Fc A (C220S, K447A, N297D, T350V, L351Y, F405A, Y407V, L234A, L235A, V264E, L309D, Q311H, N434S) - GGsGG - PRL (N59D):<br>EPKSsDKTHTCPPCPAPEAAGGPSVFLFPPKPKDTLMISRTPEVTCVVEDVSHEDPEVKFNWYVDGVEVHN<br>AKTKPREEQYdSTYRVVSVLTVDHHDWLNGKEYKCKVSNKALPAPIEKTISKAKGQPREPQVYVYPPSRDEL<br>TKNQVSLTCLVKGFYPSDIAVEWESNGQPENNYKTTPPVLDSDGSFALVSKLTVDKSRWQQGNVFSCS<br>VMHEALHSHYTQKSLSLSPGaGGsGGLPICPGGAARCQVTLRDLFDRAVVLSHYIHDLSSEMFSEFDKRYT<br>HGRGFITKAINSCHTSSLATPEDKEQAQQMNQKDFLSLIVSILRSWNEPLYHLVTEVRGMQEAPEAILSKA<br>VEIEEQTKRLLEGMELIVSQVHPETKENEIYPVWSGLPSLQMADEESRLSAYYNLLHCLRRDSHKIDNYLKLLK<br>CRIIHNNNC<br><br>Fc A (N297D, C220S, K447A, L234A, L235A, V264E, L309D, Q311H, N434S, H435R, Y436F):<br>EPKSsDKTHTCPPCPAPEAAGGPSVFLFPPKPKDTLMISRTPEVTCVVVDVSHEDPEVKFNWYVDGVEVHN<br>AKTKPREEQYdSTYRVVSVLTVDHHDWLNGKEYKCKVSNKALPAPIEKTISKAKGQPREPQVYVLPPSRDEL<br>TKNQVSLLCLVKGFYPSDIAVEWESNGQPENNYLTWPPVLDSDGSFFLYSKLTVDKSRWQQGNVFSCS<br>VMHEALHSRFTQKSLSLSPGa |
| 16. | Fc A (C220S, K447A, N297D, T350V, L351Y, F405A, Y407V, L234A, L235A, P329G, V264E, L309D, Q311H, N434S) - GGsGG - PRL N59D:<br>EPKSsDKTHTCPPCPAPEAAGGPSVFLFPPKPKDTLMISRTPEVTCVVEDVSHEDPEVKFNWYVDGVEVHN<br>AKTKPREEQYdSTYRVVSVLTVDHHDWLNGKEYKCKVSNKALGAPIEKTISKAKGQPREPQVYVYPPSRDE<br>LTKNQVSLTCLVKGFYPSDIAVEWESNGQPENNYKTTPPVLDSDGSFALVSKLTVDKSRWQQGNVFSC<br>SVMHEALHSHYTQKSLSLSPGaGGsGGLPICPGGAARCQVTLRDLFDRAVVLSHYIHDLSSEMFSEFDKRY<br>THGRGFITKAINSCHTSSLATPEDKEQAQQMNQKDFLSLIVSILRSWNEPLYHLVTEVRGMQEAPEAILSK<br>AVEIEEQTKRLLEGMELIVSQVHPETKENEIYPVWSGLPSLQMADEESRLSAYYNLLHCLRRDSHKIDNYLKLL<br>KCRIIHNNNC<br><br>Fc B (N297D, C220S, K447A, L234A, L235A, P329G, V264E, L309D, Q311H, N434S, H435R, Y436F):<br>EPKSsDKTHTCPPCPAPEAAGGPSVFLFPPKPKDTLMISRTPEVTCVVVDVSHEDPEVKFNWYVDGVEVHN<br>AKTKPREEQYdSTYRVVSVLTVDHHDWLNGKEYKCKVSNKALGAPIEKTISKAKGQPREPQVYVLPPSRDEL<br>TKNQVSLLCLVKGFYPSDIAVEWESNGQPENNYLTWPPVLDSDGSFFLYSKLTVDKSRWQQGNVFSCS<br>VMHEALHSRFTQKSLSLSPGa |
| 17. | Fc A (C220S, K447A, N297D, T350V, L351Y, F405A, Y407V, L234A, L235A, M252Y, S254T, T256E) - GGsGG - PRL N59D: |

| | |
|---|---|
| | EPKSsDKTHTCPPCPAPEAAGGPSVFLFPPKPKDTLYITREPEVTCVVVDVSHEDPEVKFNWYVDGVEVHNA<br>KTKPREEQYdSTYRVVSVLTVLHQDWLNGKEYKCKVSNKALPAPIEKTISKAKGQPREPQVYVYPPSRDELT<br>KNQVSLTCLVKGFYPSDIAVEWESNGQPENNYKTTPPVLDSDGSFALVSKLTVDKSRWQQGNVFSCSV<br>MHEALHNHYTQKSLSLSPGaGGsGGLPICPGGAARCQVTLRDLFDRAVVLSHYIHDLSSEMFSEFDKRYT<br>HGRGFITKAINSCHTSSLATPEDKEQAQQMNQKDFLSLIVSILRSWNEPLYHLVTEVRGMQEAPEAILSKA<br>VEIEEQTKRLLEGMELIVSQVHPETKENEIYPVWSGLPSLQMADEESRLSAYYNLLHCLRRDSHKIDNYLKLLK<br>CRIIHNNNC<br><br>Fc B (N297D, K447A, H435R, Y436F, L234A, L235A, M252Y, S254T, T256E):<br>EPKSsDKTHTCPPCPAPEAAGGPSVFLFPPKPKDTLYITREPEVTCVVVDVSHEDPEVKFNWYVDGVEVHNA<br>KTKPREEQYdSTYRVVSVLTVLHQDWLNGKEYKCKVSNKALPAPIEKTISKAKGQPREPQVYVLPPSRDELTK<br>NQVSLLCLVKGFYPSDIAVEWESNGQPENNYLTWPPVLDSDGSFFLYSKLTVDKSRWQQGNVFSCSV<br>MHEALHNRFTQKSLSLSPGa |
| 18. | PRL (N59D) - Fc Knob (C220S, N297D, K427A, T366W):<br>LPICPGGAARCQVTLRDLFDRAVVLSHYIHDLSSEMFSEFDKRYTHGRGFITKAINSCHTSSLATPEDKEQA<br>QQMNQKDFLSLIVSILRSWNEPLYHLVTEVRGMQEAPEAILSKAVEIEEQTKRLLEGMELIVSQVHPETKENE<br>IYPVWSGLPSLQMADEESRLSAYYNLLHCLRRDSHKIDNYLKLLKCRIIHNNNCEPKSsDKTHTCPPCPAPEL<br>LGGPSVFLFPPKPKDTLMISRTPEVTCVVVDVSHEDPEVKFNWYVDGVEVHNAKTKPREEQYdSTYRVVSVL<br>TVLHQDWLNGKEYKCKVSNKALPAPIEKTISKAKGQPREPQVYTLPPSRDELTKNQVSLWCLVKGFYPSDI<br>AVEWESNGQPENNYKTTPPVLDSDGSFFLYSKLTVDKSRWQQGNVFSCSVMHEALHNHYTQKSLSLS<br>PGaEQKLISEEDLNSAVDHHHHHH<br><br>Fc Hole (C220S, N297D, K427A, T366S, L368A, Y407V):<br>EPKSSDKTHTCPPCPAPELLGGPSVFLFPPKPKDTLMISRTPEVTCVVVDVSHEDPEVKFNWYVDGVEVHNA<br>KTKPREEQYDSTYRVVSVLTVLHQDWLNGKEYKCKVSNKALPAPIEKTISKAKGQPREPQVYTLPPSRDELTK<br>NQVSLSCAVKGFYPSDIAVEWESNGQPENNYKTTPPVLDSDGSFFLVSKLTVDKSRWQQGNVFSCSV<br>MHEALHNRFTQKSLSLSPGA |
| 19. | PRL (N59D) - GGsGG - Fc Knob (C220S, N297D, K427A, T366W):<br>LPICPGGAARCQVTLRDLFDRAVVLSHYIHDLSSEMFSEFDKRYTHGRGFITKAINSCHTSSLATPEDKEQA<br>QQMNQKDFLSLIVSILRSWNEPLYHLVTEVRGMQEAPEAILSKAVEIEEQTKRLLEGMELIVSQVHPETKENE<br>IYPVWSGLPSLQMADEESRLSAYYNLLHCLRRDSHKIDNYLKLLKCRIIHNNNCGGsGGEPKSsDKTHTCP<br>PCPAPELLGGPSVFLFPPKPKDTLMISRTPEVTCVVVDVSHEDPEVKFNWYVDGVEVHNAKTKPREEQYdST<br>YRVVSVLTVLHQDWLNGKEYKCKVSNKALPAPIEKTISKAKGQPREPQVYTLPPSRDELTKNQVSLWCLVK<br>GFYPSDIAVEWESNGQPENNYKTTPPVLDSDGSFFLYSKLTVDKSRWQQGNVFSCSVMHEALHNHYT<br>QKSLSLSPGa<br><br>Fc Hole (C220S, N297D, K427A, T366S, L368A, Y407V):<br>EPKSSDKTHTCPPCPAPELLGGPSVFLFPPKPKDTLMISRTPEVTCVVVDVSHEDPEVKFNWYVDGVEVHNA<br>KTKPREEQYDSTYRVVSVLTVLHQDWLNGKEYKCKVSNKALPAPIEKTISKAKGQPREPQVYTLPPSRDELTK<br>NQVSLSCAVKGFYPSDIAVEWESNGQPENNYKTTPPVLDSDGSFFLVSKLTVDKSRWQQGNVFSCSV<br>MHEALHNRFTQKSLSLSPGA |
| 20. | PRL (N59D, C191S, C199S) - Fc Knob (C220S, N297D, K427A, T366W):<br>LPICPGGAARCQVTLRDLFDRAVVLSHYIHDLSSEMFSEFDKRYTHGRGFITKAINSCHTSSLATPEDKEQA<br>QQMNQKDFLSLIVSILRSWNEPLYHLVTEVRGMQEAPEAILSKAVEIEEQTKRLLEGMELIVSQVHPETKENE<br>IYPVWSGLPSLQMADEESRLSAYYNLLHCLRRDSHKIDNYLKLLKCRIIHNNNSEPKSsDKTHTCPPCPAPELL<br>GGPSVFLFPPKPKDTLMISRTPEVTCVVVDVSHEDPEVKFNWYVDGVEVHNAKTKPREEQYdSTYRVVSVLT<br>VLHQDWLNGKEYKCKVSNKALPAPIEKTISKAKGQPREPQVYTLPPSRDELTKNQVSLWCLVKGFYPSDIA<br>VEWESNGQPENNYKTTPPVLDSDGSFFLYSKLTVDKSRWQQGNVFSCSVMHEALHNHYTQKSLSLSP<br>GaEQKLISEEDLNSAVDHHHHHH |

| | |
|---|---|
| | Fc Hole (C220S, N297D, K427A, T366S, L368A, Y407V):<br>EPKSSDKTHTCPPCPAPELLGGPSVFLFPPKPKDTLMISRTPEVTCVVVDVSHEDPEVKFNWYVDGVEVHNA<br>KTKPREEQYDSTYRVVSVLTVLHQDWLNGKEYKCKVSNKALPAPIEKTISKAKGQPREPQVYTLPPSRDELTK<br>NQVSLSCAVKGFYPSDIAVEWESNGQPENNYKTTPPVLDSDGSFFLVSKLTVDKSRWQQGNVFSCSV<br>MHEALHNRFTQKSLSLSPGA |
| 21. | PRL (N59D, C191S, C199S) - Fc Knob (C220S, C226S, C229S, N297D, K427A, T366W):<br>LPICPGGAARCQVTLRDLFDRAVVLSHYIHDLSSEMFSEFDKRYTHGRGFITKAINSCHTSSLATPEDKEQA<br>QQMNQKDFLSLIVSILRSWNEPLYHLVTEVRGMQEAPEAILSKAVEIEEQTKRLLEGMELIVSQVHPETKENE<br>IYPVWSGLPSLQMADEESRLSAYYNLLHCLRRDSHKIDNYLKLLKCRIIHNNNSEPKSsDKTHTSPPSPAPELL<br>GGPSVFLFPPKPKDTLMISRTPEVTCVVVDVSHEDPEVKFNWYVDGVEVHNAKTKPREEQYdSTYRVVSVLT<br>VLHQDWLNGKEYKCKVSNKALPAPIEKTISKAKGQPREPQVYTLPPSRDELTKNQVSLWCLVKGFYPSDIA<br>VEWESNGQPENNYKTTPPVLDSDGSFFLYSKLTVDKSRWQQGNVFSCSVMHEALHNHYTQKSLSLSP<br>GaEQKLISEEDLNSAVDHHHHHH<br><br>Fc Hole (C220S, N297D, K427A, T366S, L368A, Y407V):<br>EPKSSDKTHTCPPCPAPELLGGPSVFLFPPKPKDTLMISRTPEVTCVVVDVSHEDPEVKFNWYVDGVEVHNA<br>KTKPREEQYDSTYRVVSVLTVLHQDWLNGKEYKCKVSNKALPAPIEKTISKAKGQPREPQVYTLPPSRDELTK<br>NQVSLSCAVKGFYPSDIAVEWESNGQPENNYKTTPPVLDSDGSFFLVSKLTVDKSRWQQGNVFSCSV<br>MHEALHNRFTQKSLSLSPGA |
| 22. | PRL (N59D) - GGsGG- Fc Knob (C220S, K447A, N297D, T366W, L234A, L235A, V264E, L309D, Q311H, N434S):<br>LPICPGGAARCQVTLRDLFDRAVVLSHYIHDLSSEMFSEFDKRYTHGRGFITKAINSCHTSSLATPEDKEQA<br>QQMNQKDFLSLIVSILRSWNEPLYHLVTEVRGMQEAPEAILSKAVEIEEQTKRLLEGMELIVSQVHPETKENE<br>IYPVWSGLPSLQMADEESRLSAYYNLLHCLRRDSHKIDNYLKLLKCRIIHNNNCGGsGGEPKSsDKTHTCP<br>PCPAPEAAGGPSVFLFPPKPKDTLMISRTPEVTCVVEDVSHEDPEVKFNWYVDGVEVHNAKTKPREEQYdS<br>TYRVVSVLTVDHHDWLNGKEYKCKVSNKALPAPIEKTISKAKGQPREPQVYTLPPSRDELTKNQVSLWCLV<br>KGFYPSDIAVEWESNGQPENNYKTTPPVLDSDGSFFLYSKLTVDKSRWQQGNVFSCSVMHEALHSHY<br>TQKSLSLSPGa<br><br>Fc Hole (C220S, N297D, K447A, T366S, L368A, Y407V, H435R, Y436F, L234A, L235A, V264E, L309D, Q311H, N434S):<br>EPKSSDKTHTCPPCPAPEAAGGPSVFLFPPKPKDTLMISRTPEVTCVVEDVSHEDPEVKFNWYVDGVEVHN<br>AKTKPREEQYDSTYRVVSVLTVDHHDWLNGKEYKCKVSNKALPAPIEKTISKAKGQPREPQVYTLPPSRDEL<br>TKNQVSLSCAVKGFYPSDIAVEWESNGQPENNYKTTPPVLDSDGSFFLVSKLTVDKSRWQQGNVFSC<br>SVMHEALHSRFTQKSLSLSPGA |
| 23. | PRL N59D - GGsGG- Fc Knob (C220S, K447A, N297D, T366W, L234A, L235A, P329G, V264E, L309D, Q311H, N434S):<br>LPICPGGAARCQVTLRDLFDRAVVLSHYIHDLSSEMFSEFDKRYTHGRGFITKAINSCHTSSLATPEDKEQA<br>QQMNQKDFLSLIVSILRSWNEPLYHLVTEVRGMQEAPEAILSKAVEIEEQTKRLLEGMELIVSQVHPETKENE<br>IYPVWSGLPSLQMADEESRLSAYYNLLHCLRRDSHKIDNYLKLLKCRIIHNNNCGGsGGEPKSsDKTHTCP<br>PCPAPEAAGGPSVFLFPPKPKDTLMISRTPEVTCVVEDVSHEDPEVKFNWYVDGVEVHNAKTKPREEQYdS<br>TYRVVSVLTVDHHDWLNGKEYKCKVSNKALGAPIEKTISKAKGQPREPQVYTLPPSRDELTKNQVSLWCLV<br>KGFYPSDIAVEWESNGQPENNYKTTPPVLDSDGSFFLYSKLTVDKSRWQQGNVFSCSVMHEALHSHY<br>TQKSLSLSPGa<br><br>Fc Hole (C220S, N297D, K447A, T366S, L368A, Y407V, H435R, Y436F, L234A, L235A, P329G, V264E, L309D, Q311H, N434S):<br>EPKSSDKTHTCPPCPAPEAAGGPSVFLFPPKPKDTLMISRTPEVTCVVEDVSHEDPEVKFNWYVDGVEVHN<br>AKTKPREEQYDSTYRVVSVLTVDHHDWLNGKEYKCKVSNKALGAPIEKTISKAKGQPREPQVYTLPPSRDEL |

| | |
|---|---|
| | TKNQVSLSCAVKGFYPSDIAVEWESNGQPENNYKTTPPVLDSDGSFFLVSKLTVDKSRWQQGNVFSC SVMHEALHSRFTQKSLSLSPGA |
| 24. | PRL N59D - GGsGG - Fc Knob (C220S, K447A, N297D, L234A, L235A, M252Y, S254T, T256E):<br>LPICPGGAARCQVTLRDLFDRAVVLSHYIHDLSSEMFSEFDKRYTHGRGFITKAINSCHTSSLATPEDKEQA QQMNQKDFLSLIVSILRSWNEPLYHLVTEVRGMQEAPEAILSKAVEIEEQTKRLLEGMELIVSQVHPETKENE IYPVWSGLPSLQMADEESRLSAYYNLLHCLRRDSHKIDNYLKLLKCRIIHNNNCGGsGGEPKSsDKTHTCP PCPAPEAAGGPSVFLFPPKPKDTLYISREPEVTCVVVDVSHEDPEVKFNWYVDGVEVHNAKTKPREEQYdST YRVVSVLTVLHQDWLNGKEYKCKVSNKALPAPIEKTISKAKGQPREPQVYTLPPSRDELTKNQVSLWCLVK GFYPSDIAVEWESNGQPENNYKTTPPVLDSDGSFFLYSKLTVDKSRWQQGNVFSCSVMHEALHNHYT QKSLSLSPGa<br><br>Fc Hole (C220S, N297D, K447A, T366S, L368A, Y407V, H435R, Y436F, L234A, L235A, M252Y, S254T, T256E):<br>EPKSSDKTHTCPPCPAPEAAGGPSVFLFPPKPKDTLYITREPEVTCVVVDVSHEDPEVKFNWYVDGVEVHN AKTKPREEQYDSTYRVVSVLTVLHQDWLNGKEYKCKVSNKALPAPIEKTISKAKGQPREPQVYTLPPSRDELT KNQVSLSCAVKGFYPSDIAVEWESNGQPENNYKTTPPVLDSDGSFFLVSKLTVDKSRWQQGNVFSCS VMHEALHNRFTQKSLSLSPGA |
| 25. | PRL (N59D) - GGsGG - Fc A (C220S, K447A, N297D, T350V, L351Y, F405A, Y407V):<br>LPICPGGAARCQVTLRDLFDRAVVLSHYIHDLSSEMFSEFDKRYTHGRGFITKAINSCHTSSLATPEDKEQA QQMNQKDFLSLIVSILRSWNEPLYHLVTEVRGMQEAPEAILSKAVEIEEQTKRLLEGMELIVSQVHPETKENE IYPVWSGLPSLQMADEESRLSAYYNLLHCLRRDSHKIDNYLKLLKCRIIHNNNCGGsGGEPKSsDKTHTCP PCPAPELLGGPSVFLFPPKPKDTLMISRTPEVTCVVVDVSHEDPEVKFNWYVDGVEVHNAKTKPREEQYdST YRVVSVLTVLHQDWLNGKEYKCKVSNKALPAPIEKTISKAKGQPREPQVYVYPPSRDELTKNQVSLTCLVK GFYPSDIAVEWESNGQPENNYKTTPPVLDSDGSFALVSKLTVDKSRWQQGNVFSCSVMHEALHNHYT QKSLSLSPGa<br><br>Fc B (C220S, N297D, K427A, T350V, T366L, K293L, T394W):<br>EPKSsDKTHTCPPCPAPELLGGPSVFLFPPKPKDTLMISRTPEVTCVVVDVSHEDPEVKFNWYVDGVEVHNA KTKPREEQYdSTYRVVSVLTVLHQDWLNGKEYKCKVSNKALPAPIEKTISKAKGQPREPQVYVLPPSRDELTK NQVSLLCLVKGFYPSDIAVEWESNGQPENNYLTWPPVLDSDGSFFLYSKLTVDKSRWQQGNVFSCSV MHEALHNRFTQKSLSLSPGa |
| 26. | PRL (N59D) - GGsGG- Fc A (C220S, K447A, N297D, T350V, L351Y, F405A, Y407V, L234A, L235A, V264E, L309D, Q311H, N434S):<br>LPICPGGAARCQVTLRDLFDRAVVLSHYIHDLSSEMFSEFDKRYTHGRGFITKAINSCHTSSLATPEDKEQA QQMNQKDFLSLIVSILRSWNEPLYHLVTEVRGMQEAPEAILSKAVEIEEQTKRLLEGMELIVSQVHPETKENE IYPVWSGLPSLQMADEESRLSAYYNLLHCLRRDSHKIDNYLKLLKCRIIHNNNCGGsGGEPKSsDKTHTCP PCPAPEAAGGPSVFLFPPKPKDTLMISRTPEVTCVVEDVSHEDPEVKFNWYVDGVEVHNAKTKPREEQYdS TYRVVSVLTVDHHDWLNGKEYKCKVSNKALPAPIEKTISKAKGQPREPQVYVYPPSRDELTKNQVSLTCLVK GFYPSDIAVEWESNGQPENNYKTTPPVLDSDGSFALVSKLTVDKSRWQQGNVFSCSVMHEALHSHYT QKSLSLSPGa<br><br>Fc B (N297D, C220S, K447A, L234A, L235A, V264E, L309D, Q311H, N434S):<br>EPKSsDKTHTCPPCPAPEAAGGPSVFLFPPKPKDTLMISRTPEVTCVVVDVSHEDPEVKFNWYVDGVEVHN AKTKPREEQYdSTYRVVSVLTVDHHDWLNGKEYKCKVSNKALPAPIEKTISKAKGQPREPQVYVLPPSRDEL TKNQVSLLCLVKGFYPSDIAVEWESNGQPENNYLTWPPVLDSDGSFFLYSKLTVDKSRWQQGNVFSCS VMHEALHSHYTQKSLSLSPGa |
| 27. | PRL (N59D) - GGsGG- Fc A (C220S, K447A, N297D, T350V, L351Y, F405A, Y407V, L234A, L235A, V264E, L309D, Q311H, N434S): |

| | |
|---|---|
| | LPICPGGAARCQVTLRDLFDRAVVLSHYIHDLSSEMFSEFDKRYTHGRGFITKAINSCHTSSLATPEDKEQA<br>QQMNQKDFLSLIVSILRSWNEPLYHLVTEVRGMQEAPEAILSKAVEIEEQTKRLLEGMELIVSQVHPETKENE<br>IYPVWSGLPSLQMADEESRLSAYYNLLHCLRRDSHKIDNYLKLLKCRIIHNNNCGGsGGEPKSsDKTHTCP<br>PCPAPEAAGGPSVFLFPPKPKDTLMISRTPEVTCVVEDVSHEDPEVKFNWYVDGVEVHNAKTKPREEQYdS<br>TYRVVSVLTVDHHDWLNGKEYKCKVSNKALPAPIEKTISKAKGQPREPQVYVYPPSRDELTKNQVSLTCLVK<br>GFYPSDIAVEWESNGQPENNYKTTPPVLDSDGSFALVSKLTVDKSRWQQGNVFSCSVMHEALHSHYT<br>QKSLSLSPGa<br><br>Fc B (N297D, C220S, K447A, L234A, L235A, V264E, L309D, Q311H, N434S, H435R, Y436F):<br>EPKSsDKTHTCPPCPAPEAAGGPSVFLFPPKPKDTLMISRTPEVTCVVVDVSHEDPEVKFNWYVDGVEVHN<br>AKTKPREEQYdSTYRVVSVLTVDHHDWLNGKEYKCKVSNKALPAPIEKTISKAKGQPREPQVYVLPPSRDEL<br>TKNQVSLLCLVKGFYPSDIAVEWESNGQPENNYLTWPPVLDSDGSFFLYSKLTVDKSRWQQGNVFSCS<br>VMHEALHSRFTQKSLSLSPGa |
| 28. | PRL N59D - GGsGG- Fc A (C220S, K447A, N297D, T350V, L351Y, F405A, Y407V, L234A, L235A, P329G, V264E, L309D, Q311H, N434S):<br>LPICPGGAARCQVTLRDLFDRAVVLSHYIHDLSSEMFSEFDKRYTHGRGFITKAINSCHTSSLATPEDKEQA<br>QQMNQKDFLSLIVSILRSWNEPLYHLVTEVRGMQEAPEAILSKAVEIEEQTKRLLEGMELIVSQVHPETKENE<br>IYPVWSGLPSLQMADEESRLSAYYNLLHCLRRDSHKIDNYLKLLKCRIIHNNNCGGsGGEPKSsDKTHTCP<br>PCPAPEAAGGPSVFLFPPKPKDTLMISRTPEVTCVVEDVSHEDPEVKFNWYVDGVEVHNAKTKPREEQYdS<br>TYRVVSVLTVDHHDWLNGKEYKCKVSNKALGAPIEKTISKAKGQPREPQVYVYPPSRDELTKNQVSLTCLV<br>KGFYPSDIAVEWESNGQPENNYKTTPPVLDSDGSFALVSKLTVDKSRWQQGNVFSCSVMHEALHSHY<br>TQKSLSLSPGa<br><br>Fc B (N297D, C220S, K447A, L234A, L235A, P329G, V264E, L309D, Q311H, N434S, H435R, Y436F):<br>EPKSsDKTHTCPPCPAPEAAGGPSVFLFPPKPKDTLMISRTPEVTCVVVDVSHEDPEVKFNWYVDGVEVHN<br>AKTKPREEQYdSTYRVVSVLTVDHHDWLNGKEYKCKVSNKALGAPIEKTISKAKGQPREPQVYVLPPSRDEL<br>TKNQVSLLCLVKGFYPSDIAVEWESNGQPENNYLTWPPVLDSDGSFFLYSKLTVDKSRWQQGNVFSCS<br>VMHEALHSRFTQKSLSLSPGa |
| 29. | PRL N59D - GGsGG - Fc A (C220S, K447A, N297D, T350V, L351Y, F405A, Y407V, L234A, L235A, M252Y, S254T, T256E):<br>LPICPGGAARCQVTLRDLFDRAVVLSHYIHDLSSEMFSEFDKRYTHGRGFITKAINSCHTSSLATPEDKEQA<br>QQMNQKDFLSLIVSILRSWNEPLYHLVTEVRGMQEAPEAILSKAVEIEEQTKRLLEGMELIVSQVHPETKENE<br>IYPVWSGLPSLQMADEESRLSAYYNLLHCLRRDSHKIDNYLKLLKCRIIHNNNCGGsGGEPKSsDKTHTCP<br>PCPAPEAAGGPSVFLFPPKPKDTLYITREPEVTCVVVDVSHEDPEVKFNWYVDGVEVHNAKTKPREEQYdST<br>YRVVSVLTVLHQDWLNGKEYKCKVSNKALPAPIEKTISKAKGQPREPQVYVYPPSRDELTKNQVSLTCLVK<br>GFYPSDIAVEWESNGQPENNYKTTPPVLDSDGSFALVSKLTVDKSRWQQGNVFSCSVMHEALHNHYT<br>QKSLSLSPGa<br><br>Fc B (N297D, K447A, H435R, Y436F, L234A, L235A, M252Y, S254T, T256E):<br>EPKSsDKTHTCPPCPAPEAAGGPSVFLFPPKPKDTLYITREPEVTCVVVDVSHEDPEVKFNWYVDGVEVHNA<br>KTKPREEQYdSTYRVVSVLTVLHQDWLNGKEYKCKVSNKALPAPIEKTISKAKGQPREPQVYVLPPSRDELTK<br>NQVSLLCLVKGFYPSDIAVEWESNGQPENNYLTWPPVLDSDGSFFLYSKLTVDKSRWQQGNVFSCSV<br>MHEALHNRFTQKSLSLSPGa |

Human Prolactin WT (P01236 29-227)
Signal sequence
Human Prolactin WT
C-myc tag + 6xHis tag

TTTAAAGCCGCCACCATGGAGACAGACACACTCCTGCTATGGGTACTGCTGCTCTGGGTTCCAGGTG
AGAGCTGCAGCCTGACTGCATaGGGGCTGGGATaGGCATAAGAATAAAGGTCTGTGTGGACAGCCT
TCTGaTTCAGCCACGACCTCTGTGTATcCTTCTcACCCCAcagGTTCCACCGGTCTCCCGATATGTCCGG
GCGGGGCCGCTCGGTGCCAGGTAACTTTGAGGGACCTGTTTGACCGAGCCGTAGTCCTTTCACACTATA
TTCACAACCTCTCATCTGAGATGTTTTCCGAGTTCGACAAGAGATATACCCACGGTCGCGGGTTTATAACTA
AGGCAATAAACAGTTGCCATACCTCAAGTCTCGCTACACCCGAGGACAAGGAACAAGCGCAACAGATG
AATCAGAAGGACTTTTTGTCACTGATAGTGTCCATCCTGCGCAGTTGGAACGAACCCTTGTACCATTTGGTC
ACCGAAGTCAGGGGGATGCAAGAAGCACCGGAGGCTATACTGTCAAAGGCCGTAGAAATCGAAGAAC
AGACGAAGAGACTCCTGGAAGGTATGGAACTCATAGTGTCCCAGGTCCACCCAGAGACAAAAGAGAAC
GAAATATACCCCGTATGGTCTGGCTTGCCTTCCCTGCAAATGGCAGATGAAGAGAGTCGGTTGAGTGCCT
ATTACAACCTTCTCCACTGTCTCAGGAGGGACAGTCACAAGATCGATAACTATCTCAAACTCCTTAAGTGTA
GGATAATTCATAACAATAACTGTGAACAAAAACTCATCTCAGAAGAGGATCTGAATAGCGCCGTCGACC
ATCATCATCATCATCATTGA

Human Prolactin N59D
Signal sequence
Human Prolactin N59D
C-myc tag + 6xHis tag

TTTAAAGCCGCCACCATGGAGACAGACACACTCCTGCTATGGGTACTGCTGCTCTGGGTTCCAGGTG
AGAGCTGCAGCCTGACTGCATaGGGGCTGGGATaGGCATAAGAATAAAGGTCTGTGTGGACAGCCT
TCTGaTTCAGCCACGACCTCTGTGTATcCTTCTcACCCCAcagGTTCCACCGGTCTCCCGATATGTCCGG
GCGGGGCCGCTCGGTGCCAGGTAACTTTGAGGGACCTGTTTGACCGAGCCGTAGTCCTTTCACACTATA
TTCACGACCTCTCATCTGAGATGTTTTCCGAGTTCGACAAGAGATATACCCACGGTCGCGGGTTTATAACTA
AGGCAATAAACAGTTGCCATACCTCAAGTCTCGCTACACCCGAGGACAAGGAACAAGCGCAACAGATG
AATCAGAAGGACTTTTTGTCACTGATAGTGTCCATCCTGCGCAGTTGGAACGAACCCTTGTACCATTTGGTC
ACCGAAGTCAGGGGGATGCAAGAAGCACCGGAGGCTATACTGTCAAAGGCCGTAGAAATCGAAGAAC
AGACGAAGAGACTCCTGGAAGGTATGGAACTCATAGTGTCCCAGGTCCACCCAGAGACAAAAGAGAAC
GAAATATACCCCGTATGGTCTGGCTTGCCTTCCCTGCAAATGGCAGATGAAGAGAGTCGGTTGAGTGCCT
ATTACAACCTTCTCCACTGTCTCAGGAGGGACAGTCACAAGATCGATAACTATCTCAAACTCCTTAAGTGTA
GGATAATTCATAACAATAACTGTGAACAAAAACTCATCTCAGAAGAGGATCTGAATAGCGCCGTCGACC
ATCATCATCATCATCATTGA

Human Prolactin Receptor
Signal sequence
Flag tag
Human Prolactin Receptor

TTTAAAGCCGCCACCATGGAGACAGACACACTCCTGCTATGGGTACTGCTGCTCTGGGTTCCAGGTG
AGAGCTGCAGCCTGACTGCATaGGGGCTGGGATaGGCATAAGAATAAAGGTCTGTGTGGACAGCCT
TCTGaTTCAGCCACGACCTCTGTGTATcCTTCTcACCCCAcagGTTCCACCGGTgactacaaagacgatgacgac
aagCAGCTCCCCCCTGGCAAACCTGAAATCTTTAAGTGCAGAAGTCCAAACAAGGAGACTTTCACATGCT
GGTGGCGACCCGGTACTGACGGTGGCCTCCCAACTAACTACTCTCTTACCTATCATAGGGAGGGCGAA
ACTTTGATGCACGAGTGTCCAGACTATATAACAGGAGGTCCAAATAGTTGTCACTTCGGGAAACAGTACAC
TTCTATGTGGAGAACCTATATCATGATGGTTAACGCAACTAACCAAATGGGAAGCAGTTTTAGCGATGAACT

GTACGTAGATGTAACGTACATAGTGCAGCCAGACCCGCCTCTCGAGCTGGCGGTAGAAGTCAAGCAGC
CCGAAGATAGAAAGCCATATTTGTGGATAAAATGGAGTCCCCCGACCTTGATTGATCTGAAGACGGGCTG
GTTTACGCTTTTGTACGAGATCAGGCTTAAGCCAGAGAAGGCAGCTGAGTGGGAAATCCACTTTGCTGGG
CAGCAGACTGAGTTCAAAATCCTTTCCTTGCATCCCGGCCAAAAATATCTCGTTCAAGTGCGATGTAAACC
GGACCACGGATATTGGTCAGCGTGGTCTCCTGCAACTTTCATCCAAATTCCATCCGACTTTACAATGAACG
ACACTACGGTATGGATCAGCGTAGCAGTCTTGAGCGCCGTCATTTGTTTGATTATCGTTTGGGCCGTCGCC
CTGAAAGGCTATTCAATGGTCACTTGTATATTTCCGCCGGTACCTGGGCCTAAGATCAAGGGTTTCGATGC
TCACCTCTTGGAAAAGGGGAAGTCCGAGGAATTGCTTAGCGCTCTGGGCTGTCAAGATTTCCCTCCAACC
AGTGACTACGAGGACCTGCTTGTCGAGTATCTTGAAGTCGATGACTCTGAAGACCAACACCTGATGTCCG
TCCATTCCAAAGAGCATCCATCACAGGGAATGAAACCAACTTACCTGGACCCTGATACTGACTCCGGTCG
AGGATCTTGCGATTCACCATCCCTTTTGAGTGAGAAGTGCGAGGAACCTCAAGCCAATCCCAGCACCTTTT
ATGACCCTGAAGTAATTGAAAAGCCCGAGAACCCTGAAACAACGCATACATGGGACCCCCAATGTATCA
GTATGGAAGGCAAAATCCCATATTTTCACGCGGGGGGCTCCAAATGCTCTACCTGGCCGCTCCCACAAC
CGAGTCAACACAACCCCCGCAGTTCCTACCACAACATTACCGACGTATGCGAGCTCGCTGTGGGTCCG
GCAGGGGCACCGGCGACCCTTTTGAATGAGGCCGGTAAGGACGCCCTCAAAAGCTCTCAAACTATAAA
ATCCAGAGAGGAGGGGAAAGCCACACAACAGAGAGAGGTCGAATCCTTTCATTCTGAGACTGACCAG
GACACGCCGTGGCTGTTGCCTCAAGAGAAGACGCCGTTCGGGTCTGCGAAGCCCCTTGATTATGTAGA
GATACATAAAGTCAACAAAGACGGCGCCTTGAGTCTTCTCCCTAAACAGAGGGAAAACTCAGGTAAGCC
AAAAAAACCAGGTACACCCGAAAACAATAAGGAATATGCTAAGGTCAGCGGTGTCATGGATAACAACATA
CTGGTCCTCGTTCCTGACCCACACGCAAAAAACGTGGCATGTTTTGAAGAATCTGCTAAGGAGGCACCA
CCTAGTCTTGAACAGAACCAAGCAGAGAAGGCGCTCGCTAACTTCACAGCCACCTCCAGCAAGTGCAG
GCTTCAGCTTGGGGGCCTGGACTATCTCGACCCGGCGTGTTTTACCCACAGTTTTCATTGA

Mouse Prolactin
Signal sequence
Mouse Prolactin
C-myc tag + 6xHis tag

TTTAAAGCCGCCACCATGGAGACAGACACACTCCTGCTATGGGTACTGCTGCTCTGGGTTCCAGGTG
AGAGCTGCAGCCTGACTGCATaGGGGCTGGGATaGGCATAAGAATAAAGGTCTGTGTGGACAGCCT
TCTGaTTCAGCCACGACCTCTGTGTATcCTTCTcACCCCAcagGTTCCACCGGTCTGCCAATTTGTTCCGC
GGGAGATTGTCAAACCAGCCTGAGGGAATTGTTCGATCGAGTGGTGATCCTTAGCCATTACATTCACACG
CTGTACACTGATATGTTTATTGAGTTCGATAAGCAGTATGTTCAGGATCGAGAGTTTATGGTAAAGGTAATAAA
CGATTGCCCAACATCCAGTTTGGCGACCCCAGAAGACAAGGAACAGGCTCTCAAAGTGCCGCCGGAA
GTTCTTCTGAATCTGATTCTTAGTTTGGTCCAATCTTCTAGTGATCCACTGTTTCAGCTCATTACGGGAGTCGG
GGGGATTCAGGAGGCTCCAGAATATATCCTCAGTAGGGCAAAAGAGATAGAGGAACAGAACAAACAGC
TCCTCGAAGGTGTCGAGAAGATCATATCCCAAGCCTACCCGGAAGCTAAGGGGAACGGTATTTATTTTGT
CTGGAGTCAGTTGCCTAGTCTGCAAGGTGTCGACGAAGAATCTAAGATTCTCAGCTTGCGGAACACTATTA
GGTGCTTGCGCCGCGACAGCCATAAAGTCGACAACTTTCTGAAAGTTTTGAGGTGTCAGATCGCACATCA
AAATAATTGCGAACAAAAACTCATCTCAGAAGAGGATCTGAATAGCGCCGTCGACCATCATCATCATCA
TCATTGA

Mouse Prolactin Receptor
Signal sequence
Flag tag
Mouse Prolactin Receptor

TTTAAAGCCGCCACCATGGAGACAGACACACTCCTGCTATGGGTACTGCTGCTCTGGGTTCCAGGTG
AGAGCTGCAGCCTGACTGCATaGGGGCTGGGATaGGCATAAGAATAAAGGTCTGTGTGGACAGCCT
TCTGaTTCAGCCACGACCTCTGTGTATcCTTCTcACCCCAcagGTTCCACCGGTgactacaaagacgatgacgac

aagCAGAGTCCACCCGGCAAACCAGAGATTCATAAATGCCGCTCACCGGACAAAGAGACGTTTACGTG
CTGGTGGAATCCTGGATCTGACGGCGGCCTTCCTACAAACTACTCTCTGACCTACTCTAAGGAGGGCGA
AAAGAACACCTACGAGTGCCCTGACTACAAGACCTCCGGGCCTAATTCTTGCTTTTTCTCAAAACAATACA
CCTCAATTTGGAAAATATACATAATAACAGTGAACGCCACAAACGAGATGGGGTCAAGCACATCCGATCC
GTTGTACGTTGACGTGACCTACATCGTTGAGCCGGAGCCGCCAAGGAATCTGACACTGGAAGTTAAGCA
ACTGAAAGATAAGAAGACCTATCTTTGGGTAAAATGGCTCCCCCCAACAATAACGGACGTTAAAACAGGC
TGGTTCACAATGGAATATGAGATAAGGCTCAAATCTGAAGAGGCGGACGAGTGGGAGATACATTTCACTG
GTCACCAGACGCAGTTCAAGGTATTTGATTTGTATCCAGGGCAAAAATATCTTGTGCAGACTCGGTGTAAA
CCCGACCATGGCTATTGGTCTCGGTGGGGTCAAGAAAAGAGTATAGAAATACCAAACGACTTCACTCTGA
AAGACACTACAGTGTGGATCATAGTCGCGGTTCTTTCCGCTGTCATATGTCTGATCATGGTATGGGCCGTT
GCCCTTAAGGGATACAGCATGATGACTTGCATATTCCCCCCTGTCCCCGGACCGAAGATAAAAGGCTTTG
ACACACATTTGCTTGAGAAGGGCAAAAGTGAGGAGCTGCTGTCAGCGCTGGGATGTCAGGACTTTCCTC
CAACTTCAGACTGTGAAGATCTCTTGGTAGAGTTTCTTGAGGTTGATGATAACGAAGATGAGAGACTGATGC
CGTCACACTCCAAAGAATATCCTGGGCAGGGCGTGAAGCCGACTCACCTCGATCCTGACTCAGACTCA
GGACATGGGTCTTACGACAGTCACTCTCTCTTGTCCGAAAAATGCGAAGAACCCCAGGCTTATCCTCCTG
CTTTTCACATTCCTGAAATAACTGAGAAACCGGAAAATCCGGAGGCAAATATTCCCCCAACTCCAAATCCA
CAGAACAATACACCTAATTGTCACACCGACACAAGTAAATCAACTACATGGCCGCTGCCACCCGGACAA
CATACAAGGAGAAGTCCATATCATTCTATAGCAGACGTCTGTAAGCTGGCAGGGAGTCCTGGAGATACTC
TTGACAGTTTTCTTGATAAAGCGGAAGAAAACGTGCTGAAGTTGAGCGAGGACGCTGGTGAAGAGGAGG
TGGCTGTCCAAGAAGGCGCCAAGTCTTTCCCGTCCGATAAACAAAATACAAGCTGGCCCCCACTGCAA
GAGAAGGGCCCAATTGTATACGCGAAACCGCCGGATTACGTCGAAATCCATAAGGTAAATAAAGACGG
GGTTCTGTCTTTGCTGCCAAAACAGCGAGAAAATCACCAGACAGAGAATCCCGGAGTCCCCGAGACCA
GTAAAGAGTACGCCAAGGTTTCAGGAGTGACCGATAACAATATACTCGTGCTGGTACCGGATAGTCGCG
CTCAAAATACTGCGTTGCTCGAGGAGTCCGCTAAGAAAGTGCCACCATCATTGGAGCAAAACCAGAGTG
AGAAAGATTTGGCTTCCTTCACTGCTACCTCAAGCAATTGTCGGTTGCAACTTGGGCGACTCGATTACCTG
GACCCTACATGCTTCATGCACTCCTTCCATTGA

Human IgG1 Fc WT

gagcccaaatctTGCgacaaaactcacacatgcccaccgtgcccaggtaagccagcccaggcctcgccctccagctcaaggcgggacaggtgcc
ctagagtagcctgcatccagggacaggccccagccgggtgctgacacgtccacctccatctcttcctcagcacctgaactcctggggggaccgtcagtctt
cctcttccccccaaaacccaaggacaccctcatgatctcccggacccctgaggtcacatgcgtggtggtggacgtgagccacgaagaccctgaggtca
agttcaactggtacgtggacggcgtggaggtgcataatgccaagacaaagccgagggaggagcagtacAACagcacgtaccgtgtggtcagcgt
cctcaccgtcctgcaccaggactggctgaatggcaaggagtacaagtgcaaggtctccaacaaagccctcccagcccccatcgagaaaaccatctcc
aaagccaaaggtgggacccgtggggtgcgagggccacatggacagaggccagctcagcccaccctctgccctgagagtgaccgctgtaccaacct
ctgtccctacagggcagccccgagaaccacaggtgtacaccctgcccccatccagggatgagctgaccaagaaccaggtcagcctgACCtgcctg
gtcaaaggcttctatcccagcgacatcgccgtggagtgggagagcaatgggcagccggagaacaactacaagaccacgcctcccgtgctggactcc
gacggctccttcttcctctacagcaagctcaccgtggacaagagcaggtggcagcaggggaacgtcttctcatgctccgtgatgcatgaggctctgcaca
accactacacacagaagagcctctccctgtctcccgggAAA

Human FcɣRI

Signal sequence
Human FcɣRI
C-myc tag + 6xHis tag

TTTAAAGCCGCCACCATGGAGACAGACACACTCCTGCTATGGGTACTGCTGCTCTGGGTTCCAGGTG
AGAGCTGCAGCCTGACTGCATaGGGGCTGGGATaGGCATAAGAATAAAGGTCTGTGTGGACAGCCT
TCTGaTTCAGCCACGACCTCTGTGTATcCTTCTcACCCCAcagGTTCCACCGGTCAAGTGGACACCACAA
AGGCAGTGATCACTTTGCAGCCTCCATGGGTCAGCGTGTTCCAAGAGGAAACCGTAACCTTGCACTGTG
AGGTGCTCCATCTGCCTGGGAGCAGCTCTACACAGTGGTTTCTCAATGGCACAGCCACTCAGACCTCGA
CCCCCAGCTACAGAATCACCTCTGCCAGTGTCAATGACAGTGGTGAATACAGGTGCCAGAGAGGTCTCT

CAGGGCGAAGTGACCCCATACAGCTGGAAATCCACAGAGGCTGGCTACTACTGCAGGTCTCCAGCAG
AGTCTTCACGGAAGGAGAACCTCTGGCCTTGAGGTGTCATGCGTGGAAGGATAAGCTGGTGTACAATGT
GCTTTACTATCGAAATGGCAAAGCCTTTAAGTTTTTCCACTGGAATTCTAACCTCACCATTCTGAAAACCAACA
TAAGTCACAATGGCACCTACCATTGCTCAGGCATGGGAAAGCATCGCTACACATCAGCAGGAATATCTGT
CACTGTGAAAGAGCTATTTCCAGCTCCAGTGCTGAATGCATCTGTGACATCCCCACTCCTGGAGGGGAAT
CTGGTCACCCTGAGCTGTGAAACAAAGTTGCTCTTGCAGAGGCCTGGTTTGCAGCTTTACTTCTCCTTCTAC
ATGGGCAGCAAGACCCTGCGAGGCAGGAACACATCCTCTGAATACCAAATACTAACTGCTAGAAGAGA
AGACTCTGGGTTATACTGGTGCGAGGCTGCCACAGAGGATGGAAATGTCCTTAAGCGCAGCCCTGAGTT
GGAGCTTCAAGTGCTTGGCCTCCAGTTACCAACTCCTGTCTGGTTTCATGAACAAAAACTCATCTCAGAAG
AGGATCTGAATAGCGCCGTCGACCATCATCATCATCATCATTGA

<u>Human FcɣRIIa</u>
Signal sequence
Human FcɣRIIa
C-myc tag + 6xHis tag

TTTAAAGCCGCCACCATGGAGACAGACACACTCCTGCTATGGGTACTGCTGCTCTGGGTTCCAGGTG
AGAGCTGCAGCCTGACTGCATaGGGGCTGGGATaGGCATAAGAATAAAGGTCTGTGTGGACAGCCT
TCTGaTTCAGCCACGACCTCTGTGTATcCTTCTcACCCCAcagGTTCCACCGGTCAAGCTGCAGCTCCCC
CAAAGGCTGTGCTGAAACTTGAGCCCCCGTGGATCAACGTGCTCCAGGAGGACTCTGTGACTCTGACAT
GCCAGGGGGCTCGCAGCCCTGAGAGCGACTCCATTCAGTGGTTCCACAATGGGAATCTCATTCCCACC
CACACGCAGCCCAGCTACAGGTTCAAGGCCAACAACAATGACAGCGGGGAGTACACGTGCCAGACT
GGCCAGACCAGCCTCAGCGACCCTGTGCATCTGACTGTGCTTTCCGAATGGCTGGTGCTCCAGACCCC
TCACCTGGAGTTCCAGGAGGGAGAAACCATCATGCTGAGGTGCCACAGCTGGAAGGACAAGCCTCTG
GTCAAGGTCACATTCTTCCAGAATGGAAAATCCCAGAAATTCTCCCATTTGGATCCCACCTTCTCCATCCCA
CAAGCAAACCACAGTCACAGTGGTGATTACCACTGCACAGGAAACATAGGCTACACGCTGTTCTCATCC
AAGCCTGTGACCATCACTGTCCAAGTGCCCAGCATGGGCAGCTCTTCACCAATGGGGGAACAAAAACT
CATCTCAGAAGAGGATCTGAATAGCGCCGTCGACCATCATCATCATCATCATTGA

<u>Human FcɣRIIb</u>
Signal sequence
Human FcɣRIIb
C-myc tag + 6xHis tag

TTTAAAGCCGCCACCATGGAGACAGACACACTCCTGCTATGGGTACTGCTGCTCTGGGTTCCAGGTG
AGAGCTGCAGCCTGACTGCATaGGGGCTGGGATaGGCATAAGAATAAAGGTCTGTGTGGACAGCCT
TCTGaTTCAGCCACGACCTCTGTGTATcCTTCTcACCCCAcagGTTCCACCGGTACACCTGCAGCTCCCC
CAAAGGCTGTGCTGAAACTCGAGCCCCAGTGGATCAACGTGCTCCAGGAGGACTCTGTGACTCTGACA
TGCCGGGGGACTCACAGCCCTGAGAGCGACTCCATTCAGTGGTTCCACAATGGGAATCTCATTCCCAC
CCACACGCAGCCCAGCTACAGGTTCAAGGCCAACAACAATGACAGCGGGGAGTACACGTGCCAGAC
TGGCCAGACCAGCCTCAGCGACCCTGTGCATCTGACTGTGCTTTCTGAGTGGCTGGTGCTCCAGACCC
CTCACCTGGAGTTCCAGGAGGGAGAAACCATCGTGCTGAGGTGCCACAGCTGGAAGGACAAGCCTCT
GGTCAAGGTCACATTCTTCCAGAATGGAAAATCCAAGAAATTTTCCCGTTCGGATCCCAACTTCTCCATCCC
ACAAGCAAACCACAGTCACAGTGGTGATTACCACTGCACAGGAAACATAGGCTACACGCTGTACTCATC
CAAGCCTGTGACCATCACTGTCCAAGCTCCCGAACAAAAACTCATCTCAGAAGAGGATCTGAATAGCG
CCGTCGACCATCATCATCATCATCATTGA

<u>Human FcɣRIIIa</u>
Signal sequence
Human FcɣRIIIa

C-myc tag + 6xHis tag

TTTAAAGCCGCCACCATGGAGACAGACACACTCCTGCTATGGGTACTGCTGCTCTGGGTTCCAGGTG
AGAGCTGCAGCCTGACTGCATaGGGGCTGGGATaGGCATAAGAATAAAGGTCTGTGTGGACAGCCT
TCTGaTTCAGCCACGACCTCTGTGTATcCTTCTcACCCCAcagGTTCCACCGGTGGCATGCGGACTGAAG
ATCTCCCAAAGGCTGTGGTGTTCCTGGAGCCTCAATGGTACAGGGTGCTCGAGAAGGACAGTGTGACTC
TGAAGTGCCAGGGAGCCTACTCCCCTGAGGACAATTCCACACAGTGGTTTCACAATGAGAGCCTCATCT
CAAGCCAGGCCTCGAGCTACTTCATTGACGCTGCCACAGTCGACGACAGTGGAGAGTACAGGTGCCA
GACAAACCTCTCCACCCTCAGTGACCCGGTGCAGCTAGAAGTCCATATCGGCTGGCTGTTGCTCCAGG
CCCCTCGGTGGGTGTTCAAGGAGGAAGACCCTATTCACCTGAGGTGTCACAGCTGGAAGAACACTGCT
CTGCATAAGGTCACATATTTACAGAATGGCAAAGGCAGGAAGTATTTTCATCATAATTCTGACTTCTACATTCC
AAAAGCCACACTCAAAGACAGCGGCTCCTACTTCTGCAGGGGGCTTTTTGGGAGTAAAAATGTGTCTTCA
GAGACTGTGAACATCACCATCACTCAAGGTTTGGCAGTGTCAACCATCTCATCATTCTTTCCACCTGGGTAC
CAAGAACAAAAACTCATCTCAGAAGAGGATCTGAATAGCGCCGTCGACCATCATCATCATCATCATTGA

Mouse FcγRI
Signal sequence
Mosue FcγRI
C-myc tag + 6xHis tag

TTTAAAGCCGCCACCATGGAGACAGACACACTCCTGCTATGGGTACTGCTGCTCTGGGTTCCAGGTG
AGAGCTGCAGCCTGACTGCATaGGGGCTGGGATaGGCATAAGAATAAAGGTCTGTGTGGACAGCCT
TCTGaTTCAGCCACGACCTCTGTGTATcCTTCTcACCCCAcagGTTCCACCGGTGAAGTGGTTAATGCCAC
CAAGGCTGTGATCACCTTGCAGCCTCCATGGGTCAGTATTTTCCAGAAGGAAAATGTCACTTTATGGTGTG
AGGGGCCTCACCTGCCTGGAGACAGTTCCACACAATGGTTTATCAACGGAACAGCCGTTCAGATCTCCA
CGCCTAGTTATAGCATCCCAGAGGCCAGTTTTCAGGACAGTGGCGAATACAGGTGTCAGATAGGTTCCTC
AATGCCAAGTGACCCTGTGCAGTTGCAAATCCACAATGATTGGCTGCTACTCCAGGCCTCCCGCAGAGT
CCTCACAGAAGGAGAACCCCTGGCCTTGAGGTGTCACGGATGGAAGAATAAACTGGTGTACAATGTGGT
TTTCTATAGAAATGGAAAATCCTTTCAGTTTTCTTCAGATTCGGAGGTCGCCATTCTGAAAACCAACCTGAGTC
ACAGCGGCATCTACCACTGCTCAGGCACGGGAAGACACCGCTACACATCTGCAGGAGTGTCCATCAC
GGTGAAAGAGCTGTTTACCACGCCAGTGCTGAGAGCATCCGTGTCATCTCCCTTCCCGGAGGGGAGTC
TGGTCACCCTGAACTGTGAGACGAATTTGCTCCTGCAGAGACCCGGCTTACAGCTTCACTTCTCCTTCTAC
GTGGGCAGCAAGATCCTGGAGTACAGGAACACATCCTCAGAGTACCATATAGCAAGGGCGGAAAGAG
AAGATGCTGGATTCTACTGGTGTGAGGTAGCCACGGAGGACAGCAGTGTCCTTAAGCGCAGCCCTGAG
TTGGAGCTCCAAGTGCTTGGTCCCCAGTCATCAGCTCCTGAACAAAAACTCATCTCAGAAGAGGATCTG
AATAGCGCCGTCGACCATCATCATCATCATCATTGA

Mouse FcγRIIb
Signal sequence
Mosue FcγRIIb
C-myc tag + 6xHis tag

TTTAAAGCCGCCACCATGGAGACAGACACACTCCTGCTATGGGTACTGCTGCTCTGGGTTCCAGGTG
AGAGCTGCAGCCTGACTGCATaGGGGCTGGGATaGGCATAAGAATAAAGGTCTGTGTGGACAGCCT
TCTGaTTCAGCCACGACCTCTGTGTATcCTTCTcACCCCAcagGTTCCACCGGTactcatgatcttccaaaggctgtggt
caaactcgagcccccgtggatccaggtgctcaaggaagacacggtgacactgacatgcgaagggacccacaaccctgggaactcttctacccagtg
gttccacaatgggaggtccatccggagccaggtccaagccagctacacgtttaaggccacagtcaatgacagtggagaatatcggtgtcaaatggagc
agacccgcctcagcgaccctgtagatctgggagtgatttctgactggctgctgctccagacccctcagctggtgtttctggaaggggaaaccatcacgctaa
ggtgccatagctggaggaacaaactactgaacaggatctcgttcttccataatgaaaaatccgtgaggtatcatcactacagtagtaatttctctatcccaaa
agccaaccacagtcacagtggggactactactgcaaaggaagtctaggaaggacactgcaccagtccaagcctgtcaccatcactgtccaagggcc

caagtccagcaggtctttaccaGAACAAAAACTCATCTCAGAAGAGGATCTGAATAGCGCCGTCGACCATCATCA
TCATCATCATTGA

<u>Mouse FcɣRIII</u>
Signal sequence
Mosue FcɣRIII
C-myc tag + 6xHis tag

TTTAAAGCCGCCACCATGGAGACAGACACACTCCTGCTATGGGTACTGCTGCTCTGGGTTCCAGGTG
AGAGCTGCAGCCTGACTGCATaGGGGCTGGGATaGGCATAAGAATAAAGGTCTGTGTGGACAGCCT
TCTGaTTCAGCCACGACCTCTGTGTATcCTTCTcACCCCAcagGTTCCACCGGTgctcttccgaaggctgtggtgaaac
tggaccccccatggatccaggtgctcaaggaagacatggtgacactgatgtgcgaagggacccacaaccctgggaactcttctacccagtggttccac
aacgggaggtccatccggagccaggtccaagccagttacacgtttaaggccacagtcaatgacagtggagaatatcggtgtcaaatggagcagacc
cgcctcagcgaccctgtagatctgggagtgatttctgactggctgctgctccagacccctcagcgggtgtttctggaaggggaaaccatcacgctaaggtg
ccatagctggaggaacaaactactgaacaggatctcattcttccataatgaaaaatccgtgaggtatcatcactacaaaagtaatttctctatcccaaaagc
caaccacagtcacagtggggactactactgcaaaggaagtctaggaagtacacagcaccagtccaagcctgtcaccatcactgtccaagatccagca
actacatcctccatctctctagtctggtaccacactGAACAAAAACTCATCTCAGAAGAGGATCTGAATAGCGCCGTCGAC
CATCATCATCATCATCATTGAA

<u>Mouse FcɣRIIV</u>
Signal sequence
Mosue FcɣRIIV
C-myc tag + 6xHis tag

TTTAAAGCCGCCACCATGGAGACAGACACACTCCTGCTATGGGTACTGCTGCTCTGGGTTCCAGGTG
AGAGCTGCAGCCTGACTGCATaGGGGCTGGGATaGGCATAAGAATAAAGGTCTGTGTGGACAGCCT
TCTGaTTCAGCCACGACCTCTGTGTATcCTTCTcACCCCAcagGTTCCACCGGTGGTCTCCAAAAGGCTGT
GGTGAACCTAGACCCCAAGTGGGTCAGGGTGCTTGAGGAAGACAGCGTGACCCTCAGATGCCAAGG
CACTTTCTCCCCCGAGGACAATTCTATCAAGTGGTTCCATAACGAAAGCCTCATCCCACACCAGGATGCC
AACTATGTCATCCAAAGTGCCAGAGTTAAGGACAGTGGAATGTACAGGTGCCAGACAGCCCTCTCCACG
ATCAGTGACCCAGTGCAACTAGAGGTCCATATGGGCTGGCTATTGCTTCAGACCACTAAGTGGCTGTTCC
AGGAGGGGGACCCCATTCATCTGAGATGCCACAGTTGGCAAAACAGACCTGTACGGAAGGTCACCTAT
TTACAGAACGGCAAAGGCAAGAAGTATTTCCATGAAAATTCTGAATTACTCATTCCAAAAGCTACACACAAT
GACAGTGGCTCCTACTTCTGCAGAGGGCTCATTGGACACAACAACAAATCTTCAGCATCCTTTCGTATAAG
CCTAGGCGATCCAGGGTCTCCATCCATGTTTCCACCGTGGCATCAAGAACAAAAACTCATCTCAGAAGA
GGATCTGAATAGCGCCGTCGACCATCATCATCATCATCATTGA

Table 3: DNA sequences for Fc-prolactin fusions

| Fc-PRL- # | Sequence |
|---|---|
| 1 | Fc - PRL (N59D):<br>TTTAAAGCCGCCACCATGGAGACAGACACACTCCTGCTATGGGTACTGCTGCTCTGGGTT<br>CCAGGTGAGAGCTGCAGCCTGACTGCATaGGGGCTGGGATaGGCATAAGAATAAAGGT<br>CTGTGTGGACAGCCTTCTGaTTCAGCCACGACCTCTGTGTATcCTTCTcACCCCAcagGTTCC<br>ACCGGTgagcccaaatctTGCgacaaaactcacacatgcccaccgtgcccaggtaagccagcccaggcctcgccctcca<br>gctcaaggcgggacaggtgccctagagtagcctgcatccagggacaggccccagccgggtgctgacacgtccacctccatctct<br>tcctcagcacctgaactcctggggggaccgtcagtcttcctcttccccccaaaacccaaggacaccctcatgatctcccggacccctg<br>aggtcacatgcgtggtggtggacgtgagccacgaagaccctgaggtcaagttcaactggtacgtggacggcgtggaggtgcata<br>atgccaagacaaagccgagggaggagcagtacAACagcacgtaccgtgtggtcagcgtcctcaccgtcctgcaccaggact<br>ggctgaatggcaaggagtacaagtgcaaggtctccaacaaagccctcccagcccccatcgagaaaaccatctccaaagccaa<br>aggtgggacccgtggggtgcgagggccacatggacagaggccagctcagcccaccctctgccctgagagtgaccgctgtacc<br>aacctctgtccctacagggcagccccgagaaccacaggtgtacaccctgcccccatcccgggatgagctgaccaagaaccag<br>gtcagcctgACCtgcctggtcaaaggcttctatcccagcgacatcgccgtggagtgggagagcaatgggcagccggagaaca<br>actacaagaccacgcctcccgtgctggactccgacggctccttcttcctctacagcaagctcaccgtggacaagagcaggtggca<br>gcaggggaacgtcttctcatgctccgtgatgcatgaggctctgcacaaccactacacacagaagagcctctccctgtctcccgggA<br>AATTGCCCATCTGTCCCGGCGGGGCTGCCCGATGCCAGGTGACCCTTCGAGACCTGTTT<br>GACCGCGCCGTCGTCCTGTCCCACTACATCCATGACCTCTCCTCAGAAATGTTCAGCGAA<br>TTCGATAAACGGTATACCCATGGCCGGGGGTTCATTACCAAGGCCATCAACAGCTGCCA<br>CACTTCTTCCCTTGCCACCCCCGAAGACAAGGAGCAAGCCCAACAGATGAATCAAAAAG<br>ACTTTCTGAGCCTGATAGTCAGCATATTGCGATCCTGGAATGAGCCTCTGTATCATCTGGTC<br>ACGGAAGTACGTGGTATGCAAGAAGCCCCGGAGGCTATCCTATCCAAAGCTGTAGAGAT<br>TGAGGAGCAAACCAAACGGCTTCTAGAGGGCATGGAGCTGATAGTCAGCCAGGTTCATC<br>CTGAAACCAAAGAAAATGAGATCTACCCTGTCTGGTCGGGACTTCCATCCCTGCAGATGG<br>CTGATGAAGAGTCTCGCCTTTCTGCTTATTATAACCTGCTCCACTGCCTACGCAGGGATTCA<br>CATAAAATCGACAATTATCTCAAGCTCCTGAAGTGCCGAATCATCCACAACAACAACTGCT<br>GA |
| 2. | Fc (C220S, N297D, K427A) - PRL WT:<br>TTTAAAGCCGCCACCATGGAGACAGACACACTCCTGCTATGGGTACTGCTGCTCTGGGTT<br>CCAGGTGAGAGCTGCAGCCTGACTGCATaGGGGCTGGGATaGGCATAAGAATAAAGGT<br>CTGTGTGGACAGCCTTCTGaTTCAGCCACGACCTCTGTGTATcCTTCTcACCCCAcagGTTCC<br>ACCGGTgagcccaaatctagcgacaaaactcacacatgcccaccgtgcccaggtaagccagcccaggcctcgccctcca<br>gctcaaggcgggacaggtgccctagagtagcctgcatccagggacaggccccagccgggtgctgacacgtccacctccatctct<br>tcctcagcacctgaactcctggggggaccgtcagtcttcctcttccccccaaaacccaaggacaccctcatgatctcccggacccctg<br>aggtcacatgcgtggtggtggacgtgagccacgaagaccctgaggtcaagttcaactggtacgtggacggcgtggaggtgcata<br>atgccaagacaaagccgagggaggagcagtacgacagcacgtaccgtgtggtcagcgtcctcaccgtcctgcaccaggactg<br>gctgaatggcaaggagtacaagtgcaaggtctccaacaaagccctcccagcccccatcgagaaaaccatctccaaagccaaa<br>ggtgggacccgtggggtgcgagggccacatggacagaggccagctcagcccaccctctgccctgagagtgaccgctgtacca<br>acctctgtccctacagggcagccccgagaaccacaggtgtacaccctgcccccatcccgggatgagctgaccaagaaccaggt<br>cagcctgACCtgcctggtcaaaggcttctatcccagcgacatcgccgtggagtgggagagcaatgggcagccggagaacaa<br>ctacaagaccacgcctcccgtgctggactccgacggctccttcttcctctacagcaagctcaccgtggacaagagcaggtggcag<br>caggggaacgtcttctcatgctccgtgatgcatgaggctctgcacaaccactacacacagaagagcctctccctgtctcccggggc<br>aCTCCCGATATGTCCGGGCGGGGCCGCTCGGTGCCAGGTAACTTTGAGGGACCTGTTT<br>GACCGAGCCGTAGTCCTTTCACACTATATTCACAACCTCTCATCTGAGATGTTTTCCGAGTTC<br>GACAAGAGATATACCCACGGTCGCGGGTTTATAACTAAGGCAATAAACAGTTGCCATACC<br>TCAAGTCTCGCTACACCCGAGGACAAGGAACAAGCGCAACAGATGAATCAGAAGGACTT<br>TTTGTCACTGATAGTGTCCATCCTGCGCAGTTGGAACGAACCCTTGTACCATTTGGTCACCG<br>AAGTCAGGGGGATGCAAGAAGCACCGGAGGCTATACTGTCAAAGGCCGTAGAAATCGA<br>AGAACAGACGAAGAGACTCCTGGAAGGTATGGAACTCATAGTGTCCCAGGTCCACCCA |

| | |
|---|---|
| | GAGACAAAAGAGAACGAAATATACCCCGTATGGTCTGGCTTGCCTTCCCTGCAAATGGCA<br>GATGAAGAGAGTCGGTTGAGTGCCTATTACAACCTTCTCCACTGTCTCAGGAGGGACAGT<br>CACAAGATCGATAACTATCTCAAACTCCTTAAGTGTAGGATAATTCATAACAATAACTGTGAA<br>CAAAAACTCATCTCAGAAGAGGATCTGAATAGCGCCGTCGACCATCATCATCATCATCATT<br>GA |
| 3. | Fc (C220S, N297D, K427A) - PRL (N59D):<br>TTTAAAGCCGCCACCATGGAGACAGACACACTCCTGCTATGGGTACTGCTGCTCTGGGTT<br>CCAGGTGAGAGCTGCAGCCTGACTGCATaGGGGCTGGGATaGGCATAAGAATAAAGGT<br>CTGTGTGGACAGCCTTCTGaTTCAGCCACGACCTCTGTGTATcCTTCTcACCCCAcagGTTCC<br>ACCGGTgagcccaaatctagcgacaaaactcacacatgcccaccgtgcccaggtaagccagcccaggcctcgccctcca<br>gctcaaggcgggacaggtgccctagagtagcctgcatccagggacaggccccagccgggtgctgacacgtccacctccatctct<br>tcctcagcacctgaactcctggggggaccgtcagtcttcctcttccccccaaaacccaaggacaccctcatgatctcccggacccctg<br>aggtcacatgcgtggtggtggacgtgagccacgaagaccctgaggtcaagttcaactggtacgtggacggcgtggaggtgcata<br>atgccaagacaaagccgagggaggagcagtacgacagcacgtaccgtgtggtcagcgtcctcaccgtcctgcaccaggactg<br>gctgaatggcaaggagtacaagtgcaaggtctccaacaaagccctcccagcccccatcgagaaaaccatctccaaagccaaa<br>ggtgggacccgtggggtgcgagggccacatggacagaggccagctcagcccaccctctgccctgagagtgaccgctgtacca<br>acctctgtccctacagggcagccccgagaaccacaggtgtacaccctgcccccatccagggatgagctgaccaagaaccaggt<br>cagcctgACCtgcctggtcaaaggcttctatcccagcgacatcgccgtggagtgggagagcaatgggcagccggagaacaa<br>ctacaagaccacgcctcccgtgctggactccgacggctccttcttcctctacagcaagctcaccgtggacaagagcaggtggcag<br>caggggaacgtcttctcatgctccgtgatgcatgaggctctgcacaaccactacacacagaagagcctctccctgtctcccggggc<br>aCTCCCGATATGTCCGGGCGGGGCCGCTCGGTGCCAGGTAACTTTGAGGGACCTGTTT<br>GACCGAGCCGTAGTCCTTTCACACTATATTCACGACCTCTCATCTGAGATGTTTTCCGAGTT<br>CGACAAGAGATATACCCACGGTCGCGGGTTTATAACTAAGGCAATAAACAGTTGCCATAC<br>CTCAAGTCTCGCTACACCCGAGGACAAGGAACAAGCGCAACAGATGAATCAGAAGGAC<br>TTTTTGTCACTGATAGTGTCCATCCTGCGCAGTTGGAACGAACCCTTGTACCATTTGGTCAC<br>CGAAGTCAGGGGGATGCAAGAAGCACCGGAGGCTATACTGTCAAAGGCCGTAGAAATC<br>GAAGAACAGACGAAGAGACTCCTGGAAGGTATGGAACTCATAGTGTCCCAGGTCCACC<br>CAGAGACAAAAGAGAACGAAATATACCCCGTATGGTCTGGCTTGCCTTCCCTGCAAATGG<br>CAGATGAAGAGAGTCGGTTGAGTGCCTATTACAACCTTCTCCACTGTCTCAGGAGGGACA<br>GTCACAAGATCGATAACTATCTCAAACTCCTTAAGTGTAGGATAATTCATAACAATAACTGTTG<br>A |
| 4. | Fc (C220S, N297D, K427A) - GGsGG - PRL (N59D):<br>TTTAAAGCCGCCACCATGGAGACAGACACACTCCTGCTATGGGTACTGCTGCTCTGGGTT<br>CCAGGTGAGAGCTGCAGCCTGACTGCATaGGGGCTGGGATaGGCATAAGAATAAAGGT<br>CTGTGTGGACAGCCTTCTGaTTCAGCCACGACCTCTGTGTATcCTTCTcACCCCAcagGTTCC<br>ACCGGTgagcccaaatctagcgacaaaactcacacatgcccaccgtgcccaggtaagccagcccaggcctcgccctcca<br>gctcaaggcgggacaggtgccctagagtagcctgcatccagggacaggccccagccgggtgctgacacgtccacctccatctct<br>tcctcagcacctgaactcctggggggaccgtcagtcttcctcttccccccaaaacccaaggacaccctcatgatctcccggacccctg<br>aggtcacatgcgtggtggtggacgtgagccacgaagaccctgaggtcaagttcaactggtacgtggacggcgtggaggtgcata<br>atgccaagacaaagccgagggaggagcagtacgacagcacgtaccgtgtggtcagcgtcctcaccgtcctgcaccaggactg<br>gctgaatggcaaggagtacaagtgcaaggtctccaacaaagccctcccagcccccatcgagaaaaccatctccaaagccaaa<br>ggtgggacccgtggggtgcgagggccacatggacagaggccagctcagcccaccctctgccctgagagtgaccgctgtacca<br>acctctgtccctacagggcagccccgagaaccacaggtgtacaccctgcccccatccagggatgagctgaccaagaaccaggt<br>cagcctgACCtgcctggtcaaaggcttctatcccagcgacatcgccgtggagtgggagagcaatgggcagccggagaacaa<br>ctacaagaccacgcctcccgtgctggactccgacggctccttcttcctctacagcaagctcaccgtggacaagagcaggtggcag<br>caggggaacgtcttctcatgctccgtgatgcatgaggctctgcacaaccactacacacagaagagcctctccctgtctcccggggc<br>aGGCGGTAGCGGTGGCCTCCCGATATGTCCGGGCGGGGCCGCTCGGTGCCAGGTAA<br>CTTTGAGGGACCTGTTTGACCGAGCCGTAGTCCTTTCACACTATATTCACGACCTCTCATCT<br>GAGATGTTTTCCGAGTTCGACAAGAGATATACCCACGGTCGCGGGTTTATAACTAAGGCAA<br>TAAACAGTTGCCATACCTCAAGTCTCGCTACACCCGAGGACAAGGAACAAGCGCAACAG |

| | |
|---|---|
| | ATGAATCAGAAGGACTTTTTGTCACTGATAGTGTCCATCCTGCGCAGTTGGAACGAACCCTT<br>GTACCATTTGGTCACCGAAGTCAGGGGGATGCAAGAAGCACCGGAGGCTATACTGTCAA<br>AGGCCGTAGAAATCGAAGAACAGACGAAGAGACTCCTGGAAGGTATGGAACTCATAGT<br>GTCCCAGGTCCACCCAGAGACAAAAGAGAACGAAATATACCCCGTATGGTCTGGCTTGC<br>CTTCCCTGCAAATGGCAGATGAAGAGAGTCGGTTGAGTGCCTATTACAACCTTCTCCACTG<br>TCTCAGGAGGGACAGTCACAAGATCGATAACTATCTCAAACTCCTTAAGTGTAGGATAATTC<br>ATAACAATAACTGTGAACAAAAACTCATCTCAGAAGAGGATCTGAATAGCGCCGTCGACC<br>ATCATCATCATCATCATTGA |
| 5. | Fc (C220S, K447A, N297D, L234A, L235A, V264E, L309D, Q311H, N434S) - PRL (N59D):<br>TTTAAAGCCGCCACCATGGAGACAGACACACTCCTGCTATGGGTACTGCTGCTCTGGGTT<br>CCAGGTGAGAGCTGCAGCCTGACTGCATaGGGGCTGGGATaGGCATAAGAATAAAGGT<br>CTGTGTGGACAGCCTTCTGaTTCAGCCACGACCTCTGTGTATcCTTCTcACCCCAcagGTTCC<br>ACCGGTgagcccaaatctagcgacaaaactcacacatgcccaccgtgcccaggtaagccagcccaggcctcgccctcca<br>gctcaaggcgggacaggtgccctagagtagcctgcatccagggacaggccccagccgggtgctgacacgtccacctccatctct<br>tcctcagcacctgaaGCGGCGgggggaccgtcagtcttcctcttccccccaaaacccaaggacaccctcatgatctcccgga<br>cccctgaggtcacatgcgtggtgGAAgacgtgagccacgaagaccctgaggtcaagttcaactggtacgtggacggcgtgga<br>ggtgcataatgccaagacaaagccgagggaggagcagtacgacagcacgtaccgtgtggtcagcgtcctcaccgtcGATca<br>cCATgactggctgaatggcaaggagtacaagtgcaaggtctccaacaaagccctcccagcccccatcgagaaaaccatctcc<br>aaagccaaaggtgggacccgtggggtgcgagggccacatggacagaggccagctcagcccaccctctgccctgagagtgac<br>cgctgtaccaacctctgtccctacagggcagccccgagaaccacaggtgtacaccctgcccccatccagggatgagctgaccaa<br>gaaccaggtcagcctgACCtgcctggtcaaaggcttctatcccagcgacatcgccgtggagtgggagagcaatgggcagccg<br>gagaacaactacaagaccacgcctcccgtgctggactccgacggctccttcttcctctacagcaagctcaccgtggacaagagca<br>ggtggcagcaggggaacgtcttctcatgctccgtgatgcatgaggctctgcacAGCcactacacacagaagagcctctccctgtc<br>tcccggggcaTTGCCCATCTGTCCCGGCGGGGCTGCCCGATGCCAGGTGACCCTTCGAGA<br>CCTGTTTGACCGCGCCGTCGTCCTGTCCCACTACATCCATGACCTCTCCTCAGAAATGTTC<br>AGCGAATTCGATAAACGGTATACCCATGGCCGGGGGTTCATTACCAAGGCCATCAACAG<br>CTGCCACACTTCTTCCCTTGCCACCCCCGAAGACAAGGAGCAAGCCCAACAGATGAATC<br>AAAAAGACTTTCTGAGCCTGATAGTCAGCATATTGCGATCCTGGAATGAGCCTCTGTATCAT<br>CTGGTCACGGAAGTACGTGGTATGCAAGAAGCCCCGGAGGCTATCCTATCCAAAGCTGT<br>AGAGATTGAGGAGCAAACCAAACGGCTTCTAGAGGGCATGGAGCTGATAGTCAGCCAG<br>GTTCATCCTGAAACCAAAGAAAATGAGATCTACCCTGTCTGGTCGGGACTTCCATCCCTGC<br>AGATGGCTGATGAAGAGTCTCGCCTTTCTGCTTATTATAACCTGCTCCACTGCCTACGCAG<br>GGATTCACATAAAATCGACAATTATCTCAAGCTCCTGAAGTGCCGAATCATCCACAACAAC<br>AACTGCTGA |
| 6. | Fc (C220S, K447A, N297D, L234A, L235A, P329G, V264E, L309D, Q311H, N434S) - PRL N59D:<br>TTTAAAGCCGCCACCATGGAGACAGACACACTCCTGCTATGGGTACTGCTGCTCTGGGTT<br>CCAGGTGAGAGCTGCAGCCTGACTGCATaGGGGCTGGGATaGGCATAAGAATAAAGGT<br>CTGTGTGGACAGCCTTCTGaTTCAGCCACGACCTCTGTGTATcCTTCTcACCCCAcagGTTCC<br>ACCGGTgagcccaaatctagcgacaaaactcacacatgcccaccgtgcccaggtaagccagcccaggcctcgccctcca<br>gctcaaggcgggacaggtgccctagagtagcctgcatccagggacaggccccagccgggtgctgacacgtccacctccatctct<br>tcctcagcacctgaaGCGGCGgggggaccgtcagtcttcctcttccccccaaaacccaaggacaccctcatgatctcccgga<br>cccctgaggtcacatgcgtggtgGAAgacgtgagccacgaagaccctgaggtcaagttcaactggtacgtggacggcgtgga<br>ggtgcataatgccaagacaaagccgagggaggagcagtacgacagcacgtaccgtgtggtcagcgtcctcaccgtcGATca<br>cCATgactggctgaatggcaaggagtacaagtgcaaggtctccaacaaagccctcGGAgcccccatcgagaaaaccatct<br>ccaaagccaaaggtgggacccgtggggtgcgagggccacatggacagaggccagctcagcccaccctctgccctgagagtg<br>accgctgtaccaacctctgtccctacagggcagccccgagaaccacaggtgtacaccctgcccccatccagggatgagctgacc<br>aagaaccaggtcagcctgACCtgcctggtcaaaggcttctatcccagcgacatcgccgtggagtgggagagcaatgggcagc<br>cggagaacaactacaagaccacgcctcccgtgctggactccgacggctccttcttcctctacagcaagctcaccgtggacaagag |

caggtggcagcaggggaacgtcttctcatgctccgtgatgcatgaggctctgcacAGCcactacacacagaagagcctctccct
gtctcccggggcaTTGCCCATCTGTCCCGGCGGGGCTGCCCGATGCCAGGTGACCCTTCGA
GACCTGTTTGACCGCGCCGTCGTCCTGTCCCACTACATCCATGACCTCTCCTCAGAAATGT
TCAGCGAATTCGATAAACGGTATACCCATGGCCGGGGGTTCATTACCAAGGCCATCAACA
GCTGCCACACTTCTTCCCTTGCCACCCCCGAAGACAAGGAGCAAGCCCAACAGATGAAT
CAAAAAGACTTTCTGAGCCTGATAGTCAGCATATTGCGATCCTGGAATGAGCCTCTGTATCA
TCTGGTCACGGAAGTACGTGGTATGCAAGAAGCCCCGGAGGCTATCCTATCCAAAGCTG
TAGAGATTGAGGAGCAAACCAAACGGCTTCTAGAGGGCATGGAGCTGATAGTCAGCCA
GGTTCATCCTGAAACCAAAGAAAATGAGATCTACCCTGTCTGGTCGGGACTTCCATCCCTG
CAGATGGCTGATGAAGAGTCTCGCCTTTCTGCTTATTATAACCTGCTCCACTGCCTACGCA
GGGATTCACATAAAATCGACAATTATCTCAAGCTCCTGAAGTGCCGAATCATCCACAACAA
CAACTGCTGA

7. Fc (C220S, K447A, N297D, L234A, L235A, M252Y, S254T, T256E) - PRL N59D:
TTTAAAGCCGCCACCATGGAGACAGACACACTCCTGCTATGGGTACTGCTGCTCTGGGTT
CCAGGTGAGAGCTGCAGCCTGACTGCATaGGGGCTGGGATaGGCATAAGAATAAAGGT
CTGTGTGGACAGCCTTCTGaTTCAGCCACGACCTCTGTGTATcCTTCTcACCCCAcagGTTCC
ACCGGTgagcccaaatctagcgacaaaactcacacatgcccaccgtgcccaggtaagccagcccaggcctcgccctcca
gctcaaggcgggacaggtgccctagagtagcctgcatccagggacaggccccagccgggtgctgacacgtccacctccatctct
tcctcagcacctgaaGCCGCTgggggaccgtcagtcttcctcttccccccaaaacccaaggacaccctcTATatcACCcgg
GAAcctgaggtcacatgcgtggtggtggacgtgagccacgaagaccctgaggtcaagttcaactggtacgtggacggcgtgg
aggtgcataatgccaagacaaagccgagggaggagcagtacgacagcacgtaccgtgtggtcagcgtcctcaccgtcctgca
ccaggactggctgaatggcaaggagtacaagtgcaaggtctccaacaaagccctcccagcccccatcgagaaaaccatctcc
aaagccaaaggtgggacccgtggggtgcgagggccacatggacagaggccagctcagcccaccctctgccctgagagtgac
cgctgtaccaacctctgtccctacagggcagccccgagaaccacaggtgtacaccctgcccccatccagggatgagctgaccaa
gaaccaggtcagcctgACCtgcctggtcaaaggcttctatcccagcgacatcgccgtggagtgggagagcaatgggcagccg
gagaacaactacaagaccacgcctcccgtgctggactccgacggctccttcttcctctacagcaagctcaccgtggacaagagca
ggtggcagcaggggaacgtcttctcatgctccgtgatgcatgaggctctgcacaaccactacacacagaagagcctctccctgtctc
ccggggcaCTCCCGATATGTCCGGGCGGGGCCGCTCGGTGCCAGGTAACTTTGAGGGAC
CTGTTTGACCGAGCCGTAGTCCTTTCACACTATATTCACGACCTCTCATCTGAGATGTTTTCC
GAGTTCGACAAGAGATATACCCACGGTCGCGGGTTTATAACTAAGGCAATAAACAGTTGC
CATACCTCAAGTCTCGCTACACCCGAGGACAAGGAACAAGCGCAACAGATGAATCAGAA
GGACTTTTTGTCACTGATAGTGTCCATCCTGCGCAGTTGGAACGAACCCTTGTACCATTTGG
TCACCGAAGTCAGGGGGATGCAAGAAGCACCGGAGGCTATACTGTCAAAGGCCGTAG
AAATCGAAGAACAGACGAAGAGACTCCTGGAAGGTATGGAACTCATAGTGTCCCAGGTC
CACCCAGAGACAAAAGAGAACGAAATATACCCCGTATGGTCTGGCTTGCCTTCCCTGCAA
ATGGCAGATGAAGAGAGTCGGTTGAGTGCCTATTACAACCTTCTCCACTGTCTCAGGAGG
GACAGTCACAAGATCGATAACTATCTCAAACTCCTTAAGTGTAGGATAATTCATAACAATAAC
TGTTGA

8. Fc Knob (C220S, N297D, K427A, T366W) - PRL (N59D):
TTTAAAGCCGCCACCATGGAGACAGACACACTCCTGCTATGGGTACTGCTGCTCTGGGTT
CCAGGTGAGAGCTGCAGCCTGACTGCATaGGGGCTGGGATaGGCATAAGAATAAAGGT
CTGTGTGGACAGCCTTCTGaTTCAGCCACGACCTCTGTGTATcCTTCTcACCCCAcagGTTCC
ACCGGTgagcccaaatctagcgacaaaactcacacatgcccaccgtgcccaggtaagccagcccaggcctcgccctcca
gctcaaggcgggacaggtgccctagagtagcctgcatccagggacaggccccagccgggtgctgacacgtccacctccatctct
tcctcagcacctgaactcctggggggaccgtcagtcttcctcttccccccaaaacccaaggacaccctcatgatctcccggacccctg
aggtcacatgcgtggtggtggacgtgagccacgaagaccctgaggtcaagttcaactggtacgtggacggcgtggaggtgcata
atgccaagacaaagccgagggaggagcagtacgacagcacgtaccgtgtggtcagcgtcctcaccgtcctgcaccaggactg
gctgaatggcaaggagtacaagtgcaaggtctccaacaaagccctcccagcccccatcgagaaaaccatctccaaagccaaa
ggtgggacccgtggggtgcgagggccacatggacagaggccagctcagcccaccctctgccctgagagtgaccgctgtacca
acctctgtccctacagggcagccccgagaaccacaggtgtacaccctgcccccatccagggatgagctgaccaagaaccaggt

cagcctgtggtgcctggtcaaaggcttctatcccagcgacatcgccgtggagtgggagagcaatgggcagccggagaacaact
acaagaccacgcctcccgtgctggactccgacggctccttcttcctctacagcaagctcaccgtggacaagagcaggtggcagca
ggggaacgtcttctcatgctccgtgatgcatgaggctctgcacaaccactacacacagaagagcctctccctgtctcccggggcaC
TCCCGATATGTCCGGGCGGGGCCGCTCGGTGCCAGGTAACTTTGAGGGACCTGTTTGA
CCGAGCCGTAGTCCTTTCACACTATATTCACGACCTCTCATCTGAGATGTTTTCCGAGTTCG
ACAAGAGATATACCCACGGTCGCGGGTTTATAACTAAGGCAATAAACAGTTGCCATACCTC
AAGTCTCGCTACACCCGAGGACAAGGAACAAGCGCAACAGATGAATCAGAAGGACTTTT
TGTCACTGATAGTGTCCATCCTGCGCAGTTGGAACGAACCCTTGTACCATTTGGTCACCGA
AGTCAGGGGGATGCAAGAAGCACCGGAGGCTATACTGTCAAAGGCCGTAGAAATCGAA
GAACAGACGAAGAGACTCCTGGAAGGTATGGAACTCATAGTGTCCCAGGTCCACCCAG
AGACAAAAGAGAACGAAATATACCCCGTATGGTCTGGCTTGCCTTCCCTGCAAATGGCAG
ATGAAGAGAGTCGGTTGAGTGCCTATTACAACCTTCTCCACTGTCTCAGGAGGGACAGTC
ACAAGATCGATAACTATCTCAAACTCCTTAAGTGTAGGATAATTCATAACAATAACTGTGAAC
AAAAACTCATCTCAGAAGAGGATCTGAATAGCGCCGTCGACCATCATCATCATCATCATTG
A

Fc Hole (C220S, N297D, K427A, T366S, L368A, Y407V):
atggagacagacacactcctgctatgggtactgctgctctgggttccaggtgagtgctccagcctgactgcatgggggctgggatgg
gcataagaataaaggtctgtgtggacagccttctgcttcagccacgacctctgtgaattcttctaaccccacaggttccaccggtgagc
ccaaatctagcgacaaaactcacacatgcccaccgtgcccaggtaagccagcccaggcctcgccctccagctcaaggcggga
caggtgccctagagtagcctgcatccagggacaggccccagccgggtgctgacacgtccacctccatctcttcctcagcacctgaa
ctcctggggggaccgtcagtcttcctcttccccccaaaacccaaggacaccctcatgatctcccggacccctgaggtcacatgcgtg
gtggtggacgtgagccacgaagaccctgaggtcaagttcaactggtacgtggacggcgtggaggtgcataatgccaagacaa
agccgagggaggagcagtacgacagcacgtaccgtgtggtcagcgtcctcaccgtcctgcaccaggactggctgaatggcaa
ggagtacaagtgcaaggtctccaacaaagccctcccagcccccatcgagaaaaccatctccaaagccaaaggtgggacccgt
ggggtgcgagggccacatggacagaggccagctcagcccaccctctgccctgagagtgaccgctgtaccaacctctgtccctac
agggcagccccgagaaccacaggtgtacaccctgcccccatccagggatgagctgaccaagaaccaggtcagcctgagctg
cgccgtcaaaggcttctatcccagcgacatcgccgtggagtgggagagcaatgggcagccggagaacaactacaagaccac
gcctcccgtgctggactccgacggctccttcttcctcgtcagcaagctcaccgtggacaagagcaggtggcagcaggggaacgtct
tctcatgctccgtgatgcatgaggctctgcacaaccactacacacagaagagcctctccctgtctccggggcataa

9. Fc Knob (C220S, N297D, K427A, T366W) - GGsGG - PRL (N59D):
TTTAAAGCCGCCACCATGGAGACAGACACACTCCTGCTATGGGTACTGCTGCTCTGGGTT
CCAGGTGAGAGCTGCAGCCTGACTGCATaGGGGCTGGGATaGGCATAAGAATAAAGGT
CTGTGTGGACAGCCTTCTGaTTCAGCCACGACCTCTGTGTATcCTTCTcACCCCAcagGTTCC
ACCGGTgagcccaaatctagcgacaaaactcacacatgcccaccgtgcccaggtaagccagcccaggcctcgccctcca
gctcaaggcgggacaggtgccctagagtagcctgcatccagggacaggccccagccgggtgctgacacgtccacctccatctct
tcctcagcacctgaactcctggggggaccgtcagtcttcctcttccccccaaaacccaaggacaccctcatgatctcccggacccctg
aggtcacatgcgtggtggtggacgtgagccacgaagaccctgaggtcaagttcaactggtacgtggacggcgtggaggtgcata
atgccaagacaaagccgagggaggagcagtacgacagcacgtaccgtgtggtcagcgtcctcaccgtcctgcaccaggactg
gctgaatggcaaggagtacaagtgcaaggtctccaacaaagccctcccagcccccatcgagaaaaccatctccaaagccaaa
ggtgggacccgtggggtgcgagggccacatggacagaggccagctcagcccaccctctgccctgagagtgaccgctgtacca
acctctgtccctacagggcagccccgagaaccacaggtgtacaccctgcccccatccagggatgagctgaccaagaaccaggt
cagcctgtggtgcctggtcaaaggcttctatcccagcgacatcgccgtggagtgggagagcaatgggcagccggagaacaact
acaagaccacgcctcccgtgctggactccgacggctccttcttcctctacagcaagctcaccgtggacaagagcaggtggcagca
ggggaacgtcttctcatgctccgtgatgcatgaggctctgcacaaccactacacacagaagagcctctccctgtctcccggggcaG
GCGGTAGCGGTGGCCTCCCGATATGTCCGGGCGGGGCCGCTCGGTGCCAGGTAACTT
TGAGGGACCTGTTTGACCGAGCCGTAGTCCTTTCACACTATATTCACGACCTCTCATCTGA
GATGTTTTCCGAGTTCGACAAGAGATATACCCACGGTCGCGGGTTTATAACTAAGGCAATA
AACAGTTGCCATACCTCAAGTCTCGCTACACCCGAGGACAAGGAACAAGCGCAACAGAT
GAATCAGAAGGACTTTTTGTCACTGATAGTGTCCATCCTGCGCAGTTGGAACGAACCCTTG

| | |
|---|---|
| | TACCATTTGGTCACCGAAGTCAGGGGGATGCAAGAAGCACCGGAGGCTATACTGTCAAA<br>GGCCGTAGAAATCGAAGAACAGACGAAGAGACTCCTGGAAGGTATGGAACTCATAGTGT<br>CCCAGGTCCACCCAGAGACAAAAGAGAACGAAATATACCCCGTATGGTCTGGCTTGCCT<br>TCCCTGCAAATGGCAGATGAAGAGAGTCGGTTGAGTGCCTATTACAACCTTCTCCACTGTC<br>TCAGGAGGGACAGTCACAAGATCGATAACTATCTCAAACTCCTTAAGTGTAGGATAATTCAT<br>AACAATAACTGTTGA<br><br>Fc Hole (C220S, N297D, K427A, T366S, L368A, Y407V):<br>TTTAAAGCCGCCACCATGGAGACAGACACACTCCTGCTATGGGTACTGCTGCTCTGGGTT<br>CCAGGTGAGAGCTGCAGCCTGACTGCATaGGGGCTGGGATaGGCATAAGAATAAAGGT<br>CTGTGTGGACAGCCTTCTGaTTCAGCCACGACCTCTGTGTATcCTTCTcACCCCAcagGTTCC<br>ACCGGTgagcccaaatctagcgacaaaactcacacatgcccaccgtgcccaggtaagccagcccaggcctcgccctcca<br>gctcaaggcgggacaggtgccctagagtagcctgcatccagggacaggccccagccgggtgctgacacgtccacctccatctct<br>tcctcagcacctgaactcctggggggaccgtcagtcttcctcttccccccaaaacccaaggacaccctcatgatctcccggacccctg<br>aggtcacatgcgtggtggtggacgtgagccacgaagaccctgaggtcaagttcaactggtacgtggacggcgtggaggtgcata<br>atgccaagacaaagccgagggaggagcagtacgacagcacgtaccgtgtggtcagcgtcctcaccgtcctgcaccaggactg<br>gctgaatggcaaggagtacaagtgcaaggtctccaacaaagccctcccagcccccatcgagaaaaccatctccaaagccaaa<br>ggtgggacccgtggggtgcgagggccacatggacagaggccagctcagcccaccctctgccctgagagtgaccgctgtacca<br>acctctgtccctacagggcagccccgagaaccacaggtgtacaccctgcccccatccagggatgagctgaccaagaaccaggt<br>cagcctgagctgcgccgtcaaaggcttctatcccagcgacatcgccgtggagtgggagagcaatgggcagccggagaacaac<br>tacaagaccacgcctcccgtgctggactccgacggctccttcttcctcgtcagcaagctcaccgtggacaagagcaggtggcagc<br>aggggaacgtcttctcatgctccgtgatgcatgaggctctgcacaacCGTTTTacacagaagagcctctccctgtctcccggggc<br>aTGA |
| 10. | Fc Knob (C220S, K447A, N297D, T366W, L234A, L235A, V264E, L309D, Q311H, N434S) - GGsGG - PRL (N59D):<br>TTTAAAGCCGCCACCATGGAGACAGACACACTCCTGCTATGGGTACTGCTGCTCTGGGTT<br>CCAGGTGAGAGCTGCAGCCTGACTGCATaGGGGCTGGGATaGGCATAAGAATAAAGGT<br>CTGTGTGGACAGCCTTCTGaTTCAGCCACGACCTCTGTGTATcCTTCTcACCCCAcagGTTCC<br>ACCGGTgagcccaaatctagcgacaaaactcacacatgcccaccgtgcccaggtaagccagcccaggcctcgccctcca<br>gctcaaggcgggacaggtgccctagagtagcctgcatccagggacaggccccagccgggtgctgacacgtccacctccatctct<br>tcctcagcacctgaaGCGGCGgggggaccgtcagtcttcctcttccccccaaaacccaaggacaccctcatgatctcccgga<br>cccctgaggtcacatgcgtggtgGAAgacgtgagccacgaagaccctgaggtcaagttcaactggtacgtggacggcgtgga<br>ggtgcataatgccaagacaaagccgagggaggagcagtacgacagcacgtaccgtgtggtcagcgtcctcaccgtcGATca<br>cCATgactggctgaatggcaaggagtacaagtgcaaggtctccaacaaagccctcccagcccccatcgagaaaaccatctcc<br>aaagccaaaggtgggacccgtggggtgcgagggccacatggacagaggccagctcagcccaccctctgccctgagagtgac<br>cgctgtaccaacctctgtccctacagggcagccccgagaaccacaggtgtacaccctgcccccatccagggatgagctgaccaa<br>gaaccaggtcagcctgTGGtgcctggtcaaaggcttctatcccagcgacatcgccgtggagtgggagagcaatgggcagccg<br>gagaacaactacaagaccacgcctcccgtgctggactccgacggctccttcttcctctacagcaagctcaccgtggacaagagca<br>ggtggcagcaggggaacgtcttctcatgctccgtgatgcatgaggctctgcacAGCcactacacacagaagagcctctccctgtc<br>tcccggggcaGGCGGTAGCGGTGGCTTGCCCATCTGTCCCGGCGGGGCTGCCCGATGCC<br>AGGTGACCCTTCGAGACCTGTTTGACCGCGCCGTCGTCCTGTCCCACTACATCCATGAC<br>CTCTCCTCAGAAATGTTCAGCGAATTCGATAAACGGTATACCCATGGCCGGGGGTTCATTA<br>CCAAGGCCATCAACAGCTGCCACACTTCTTCCCTTGCCACCCCCGAAGACAAGGAGCA<br>AGCCCAACAGATGAATCAAAAAGACTTTCTGAGCCTGATAGTCAGCATATTGCGATCCTGG<br>AATGAGCCTCTGTATCATCTGGTCACGGAAGTACGTGGTATGCAAGAAGCCCCGGAGGC<br>TATCCTATCCAAAGCTGTAGAGATTGAGGAGCAAACCAAACGGCTTCTAGAGGGCATGGA<br>GCTGATAGTCAGCCAGGTTCATCCTGAAACCAAAGAAAATGAGATCTACCCTGTCTGGTC<br>GGGACTTCCATCCCTGCAGATGGCTGATGAAGAGTCTCGCCTTTCTGCTTATTATAACCTGC<br>TCCACTGCCTACGCAGGGATTCACATAAAATCGACAATTATCTCAAGCTCCTGAAGTGCCG<br>AATCATCCACAACAACAACTGCTGA |

Fc Hole (C220S, N297D, K447A, T366S, L368A, Y407V, H435R, Y436F, L234A, L235A, V264E, L309D, Q311H, N434S):
TTTAAAGCCGCCACCATGGAGACAGACACACTCCTGCTATGGGTACTGCTGCTCTGGGTT
CCAGGTGAGAGCTGCAGCCTGACTGCATaGGGGCTGGGATaGGCATAAGAATAAAGGT
CTGTGTGGACAGCCTTCTGaTTCAGCCACGACCTCTGTGTATcCTTCTcACCCCAcagGTTCC
ACCGGTgagcccaaatctagcgacaaaactcacacatgcccaccgtgcccaggtaagccagcccaggcctcgccctcca
gctcaaggcgggacaggtgccctagagtagcctgcatccagggacaggccccagccgggtgctgacacgtccacctccatctct
tcctcagcacctgaaGCGGCGgggggaccgtcagtcttcctcttccccccaaaacccaaggacaccctcatgatctcccgga
cccctgaggtcacatgcgtggtgGAAgacgtgagccacgaagaccctgaggtcaagttcaactggtacgtggacggcgtgga
ggtgcataatgccaagacaaagccgagggaggagcagtacgacagcacgtaccgtgtggtcagcgtcctcaccgtcGATca
cCATgactggctgaatggcaaggagtacaagtgcaaggtctccaacaaagccctcccagcccccatcgagaaaaccatctcc
aaagccaaaggtgggacccgtggggtgcgagggccacatggacagaggccagctcagcccaccctctgccctgagagtgac
cgctgtaccaacctctgtccctacagggcagccccgagaaccacaggtgtacaccctgcccccatccagggatgagctgaccaa
gaaccaggtcagcctgagctgcgccgtcaaaggcttctatcccagcgacatcgccgtggagtgggagagcaatgggcagccg
gagaacaactacaagaccacgcctcccgtgctggactccgacggctccttcttcctcgtcagcaagctcaccgtggacaagagca
ggtggcagcaggggaacgtcttctcatgctccgtgatgcatgaggctctgcacAGCCGTTTTacacagaagagcctctccctgt
ctcccggggcaTGA

11. Fc Knob (C220S, K447A, N297D, T366W, L234A, L235A, P329G, V264E, L309D, Q311H, N434S) - GGsGG - PRL N59D:
TTTAAAGCCGCCACCATGGAGACAGACACACTCCTGCTATGGGTACTGCTGCTCTGGGTT
CCAGGTGAGAGCTGCAGCCTGACTGCATaGGGGCTGGGATaGGCATAAGAATAAAGGT
CTGTGTGGACAGCCTTCTGaTTCAGCCACGACCTCTGTGTATcCTTCTcACCCCAcagGTTCC
ACCGGTgagcccaaatctagcgacaaaactcacacatgcccaccgtgcccaggtaagccagcccaggcctcgccctcca
gctcaaggcgggacaggtgccctagagtagcctgcatccagggacaggccccagccgggtgctgacacgtccacctccatctct
tcctcagcacctgaaGCGGCGgggggaccgtcagtcttcctcttccccccaaaacccaaggacaccctcatgatctcccgga
cccctgaggtcacatgcgtggtgGAAgacgtgagccacgaagaccctgaggtcaagttcaactggtacgtggacggcgtgga
ggtgcataatgccaagacaaagccgagggaggagcagtacgacagcacgtaccgtgtggtcagcgtcctcaccgtcGATca
cCATgactggctgaatggcaaggagtacaagtgcaaggtctccaacaaagccctcGGAgcccccatcgagaaaaccatct
ccaaagccaaaggtgggacccgtggggtgcgagggccacatggacagaggccagctcagcccaccctctgccctgagagtg
accgctgtaccaacctctgtccctacagggcagccccgagaaccacaggtgtacaccctgcccccatccagggatgagctgacc
aagaaccaggtcagcctgTGGtgcctggtcaaaggcttctatcccagcgacatcgccgtggagtgggagagcaatgggcagc
cggagaacaactacaagaccacgcctcccgtgctggactccgacggctccttcttcctctacagcaagctcaccgtggacaagag
caggtggcagcaggggaacgtcttctcatgctccgtgatgcatgaggctctgcacAGCcactacacacagaagagcctctccct
gtctcccggggcaGGCGGTAGCGGTGGCTTGCCCATCTGTCCCGGCGGGGCTGCCCGATG
CCAGGTGACCCTTCGAGACCTGTTTGACCGCGCCGTCGTCCTGTCCCACTACATCCATG
ACCTCTCCTCAGAAATGTTCAGCGAATTCGATAAACGGTATACCCATGGCCGGGGGTTCA
TTACCAAGGCCATCAACAGCTGCCACACTTCTTCCCTTGCCACCCCCGAAGACAAGGAG
CAAGCCCAACAGATGAATCAAAAAGACTTTCTGAGCCTGATAGTCAGCATATTGCGATCCT
GGAATGAGCCTCTGTATCATCTGGTCACGGAAGTACGTGGTATGCAAGAAGCCCCGGAG
GCTATCCTATCCAAAGCTGTAGAGATTGAGGAGCAAACCAAACGGCTTCTAGAGGGCAT
GGAGCTGATAGTCAGCCAGGTTCATCCTGAAACCAAAGAAAATGAGATCTACCCTGTCTG
GTCGGGACTTCCATCCCTGCAGATGGCTGATGAAGAGTCTCGCCTTTCTGCTTATTATAACC
TGCTCCACTGCCTACGCAGGGATTCACATAAAATCGACAATTATCTCAAGCTCCTGAAGTG
CCGAATCATCCACAACAACAACTGCTGA

Fc Hole (C220S, N297D, K447A, T366S, L368A, Y407V, H435R, Y436F, L234A, L235A, P329G, V264E, L309D, Q311H, N434S):
TTTAAAGCCGCCACCATGGAGACAGACACACTCCTGCTATGGGTACTGCTGCTCTGGGTT
CCAGGTGAGAGCTGCAGCCTGACTGCATaGGGGCTGGGATaGGCATAAGAATAAAGGT

CTGTGTGGACAGCCTTCTGaTTCAGCCACGACCTCTGTGTATcCTTCTcACCCCAcagGTTCC
ACCGGTgagcccaaatctagcgacaaaactcacacatgcccaccgtgcccaggtaagccagcccaggcctcgccctcca
gctcaaggcgggacaggtgccctagagtagcctgcatccagggacaggccccagccgggtgctgacacgtccacctccatctct
tcctcagcacctgaaGCGGCGgggggaccgtcagtcttcctcttccccccaaaacccaaggacaccctcatgatctcccgga
cccctgaggtcacatgcgtggtgGAAgacgtgagccacgaagaccctgaggtcaagttcaactggtacgtggacggcgtgga
ggtgcataatgccaagacaaagccgagggaggagcagtacgacagcacgtaccgtgtggtcagcgtcctcaccgtcGATca
cCATgactggctgaatggcaaggagtacaagtgcaaggtctccaacaaagccctcGGAgcccccatcgagaaaaccatct
ccaaagccaaaggtgggacccgtggggtgcgagggccacatggacagaggccagctcagcccaccctctgccctgagagtg
accgctgtaccaacctctgtccctacagggcagccccgagaaccacaggtgtacaccctgcccccatccagggatgagctgacc
aagaaccaggtcagcctgagctgcgccgtcaaaggcttctatcccagcgacatcgccgtggagtgggagagcaatgggcagc
cggagaacaactacaagaccacgcctcccgtgctggactccgacggctccttcttcctcgtcagcaagctcaccgtggacaagag
caggtggcagcaggggaacgtcttctcatgctccgtgatgcatgaggctctgcacAGCCGTTTTacacagaagagcctctccc
tgtctcccggggcaTGA

12. Fc Knob (C220S, K447A, N297D, L234A, L235A, M252Y, S254T, T256E) - GGsGG - PRL N59D:
TTTAAAGCCGCCACCATGGAGACAGACACACTCCTGCTATGGGTACTGCTGCTCTGGGTT
CCAGGTGAGAGCTGCAGCCTGACTGCATaGGGGCTGGGATaGGCATAAGAATAAAGGT
CTGTGTGGACAGCCTTCTGaTTCAGCCACGACCTCTGTGTATcCTTCTcACCCCAcagGTTCC
ACCGGTgagcccaaatctagcgacaaaactcacacatgcccaccgtgcccaggtaagccagcccaggcctcgccctcca
gctcaaggcgggacaggtgccctagagtagcctgcatccagggacaggccccagccgggtgctgacacgtccacctccatctct
tcctcagcacctgaaGCCGCTgggggaccgtcagtcttcctcttccccccaaaacccaaggacaccctcTATatcACCcgg
GAAcctgaggtcacatgcgtggtggtggacgtgagccacgaagaccctgaggtcaagttcaactggtacgtggacggcgtgg
aggtgcataatgccaagacaaagccgagggaggagcagtacgacagcacgtaccgtgtggtcagcgtcctcaccgtcctgca
ccaggactggctgaatggcaaggagtacaagtgcaaggtctccaacaaagccctcccagcccccatcgagaaaaccatctcc
aaagccaaaggtgggacccgtggggtgcgagggccacatggacagaggccagctcagcccaccctctgccctgagagtgac
cgctgtaccaacctctgtccctacagggcagccccgagaaccacaggtgtacaccctgcccccatccagggatgagctgaccaa
gaaccaggtcagcctgtggtgcctggtcaaaggcttctatcccagcgacatcgccgtggagtgggagagcaatgggcagccgg
agaacaactacaagaccacgcctcccgtgctggactccgacggctccttcttcctctacagcaagctcaccgtggacaagagcag
gtggcagcaggggaacgtcttctcatgctccgtgatgcatgaggctctgcacaaccactacacacagaagagcctctccctgtctcc
cggggcaGGCGGTAGCGGTGGCCTCCCGATATGTCCGGGCGGGGCCGCTCGGTGCCA
GGTAACTTTGAGGGACCTGTTTGACCGAGCCGTAGTCCTTTCACACTATATTCACGACCTCT
CATCTGAGATGTTTTCCGAGTTCGACAAGAGATATACCCACGGTCGCGGGTTTATAACTAA
GGCAATAAACAGTTGCCATACCTCAAGTCTCGCTACACCCGAGGACAAGGAACAAGCG
CAACAGATGAATCAGAAGGACTTTTTGTCACTGATAGTGTCCATCCTGCGCAGTTGGAACG
AACCCTTGTACCATTTGGTCACCGAAGTCAGGGGGATGCAAGAAGCACCGGAGGCTATA
CTGTCAAAGGCCGTAGAAATCGAAGAACAGACGAAGAGACTCCTGGAAGGTATGGAACT
CATAGTGTCCCAGGTCCACCCAGAGACAAAAGAGAACGAAATATACCCCGTATGGTCTG
GCTTGCCTTCCCTGCAAATGGCAGATGAAGAGAGTCGGTTGAGTGCCTATTACAACCTTCT
CCACTGTCTCAGGAGGGACAGTCACAAGATCGATAACTATCTCAAACTCCTTAAGTGTAGG
ATAATTCATAACAATAACTGTTGA

Fc Hole (C220S, N297D, K447A, T366S, L368A, Y407V, H435R, Y436F, L234A, L235A, M252Y, S254T, T256E):
TTTAAAGCCGCCACCATGGAGACAGACACACTCCTGCTATGGGTACTGCTGCTCTGGGTT
CCAGGTGAGAGCTGCAGCCTGACTGCATaGGGGCTGGGATaGGCATAAGAATAAAGGT
CTGTGTGGACAGCCTTCTGaTTCAGCCACGACCTCTGTGTATcCTTCTcACCCCAcagGTTCC
ACCGGTgagcccaaatctagcgacaaaactcacacatgcccaccgtgcccaggtaagccagcccaggcctcgccctcca
gctcaaggcgggacaggtgccctagagtagcctgcatccagggacaggccccagccgggtgctgacacgtccacctccatctct
tcctcagcacctgaaGCCGCTgggggaccgtcagtcttcctcttccccccaaaacccaaggacaccctcTATatcACCcgg
GAAcctgaggtcacatgcgtggtggtggacgtgagccacgaagaccctgaggtcaagttcaactggtacgtggacggcgtgg

aggtgcataatgccaagacaaagccgagggaggagcagtacgacagcacgtaccgtgtggtcagcgtcctcaccgtcctgca
ccaggactggctgaatggcaaggagtacaagtgcaaggtctccaacaaagccctcccagcccccatcgagaaaaccatctcc
aaagccaaaggtgggacccgtggggtgcgagggccacatggacagaggccagctcagcccaccctctgccctgagagtgac
cgctgtaccaacctctgtccctacagggcagccccgagaaccacaggtgtacaccctgcccccatccagggatgagctgaccaa
gaaccaggtcagcctgagctgcgccgtcaaaggcttctatcccagcgacatcgccgtggagtgggagagcaatgggcagccg
gagaacaactacaagaccacgcctcccgtgctggactccgacggctccttcttcctcgtcagcaagctcaccgtggacaagagca
ggtggcagcaggggaacgtcttctcatgctccgtgatgcatgaggctctgcacaacCGTTTTacacagaagagcctctccctgtc
tcccggggcaTGA

13.

Fc A (C220S, K447A, N297D, T350V, L351Y, F405A, Y407V) - GGsGG - PRL (N59D):
TTTAAAGCCGCCACCATGGAGACAGACACACTCCTGCTATGGGTACTGCTGCTCTGGGTT
CCAGGTGAGAGCTGCAGCCTGACTGCATaGGGGCTGGGATaGGCATAAGAATAAAGGT
CTGTGTGGACAGCCTTCTGaTTCAGCCACGACCTCTGTGTATcCTTCTcACCCCAcagGTTCC
ACCGGTgagcccaaatctagcgacaaaactcacacatgcccaccgtgcccaggtaagccagcccaggcctcgccctcca
gctcaaggcgggacaggtgccctagagtagcctgcatccagggacaggccccagccgggtgctgacacgtccacctccatctct
tcctcagcacctgaactcctggggggaccgtcagtcttcctcttccccccaaaacccaaggacaccctcatgatctcccggacccctg
aggtcacatgcgtggtggtggacgtgagccacgaagaccctgaggtcaagttcaactggtacgtggacggcgtggaggtgcata
atgccaagacaaagccgagggaggagcagtacgacagcacgtaccgtgtggtcagcgtcctcaccgtcctgcaccaggactg
gctgaatggcaaggagtacaagtgcaaggtctccaacaaagccctcccagcccccatcgagaaaaccatctccaaagccaaa
ggtgggacccgtggggtgcgagggccacatggacagaggccagctcagcccaccctctgccctgagagtgaccgctgtacca
acctctgtccctacagggcagccccgagaaccacaggtgtacGTGTATcccccatccagggatgagctgaccaagaaccag
gtcagcctgCTGtgcctggtcaaaggcttctatcccagcgacatcgccgtggagtgggagagcaatgggcagccggagaaca
actacaagaccacgcctcccgtgctggactccgacggctccttcGCGctcGTGagcaagctcaccgtggacaagagcaggt
ggcagcaggggaacgtcttctcatgctccgtgatgcatgaggctctgcacaaccactacacacagaagagcctctccctgtctccc
ggggcaGGCGGTAGCGGTGGCCTCCCGATATGTCCGGGCGGGGCCGCTCGGTGCCAG
GTAACTTTGAGGGACCTGTTTGACCGAGCCGTAGTCCTTTCACACTATATTCACGACCTCTC
ATCTGAGATGTTTTCCGAGTTCGACAAGAGATATACCCACGGTCGCGGGTTTATAACTAAG
GCAATAAACAGTTGCCATACCTCAAGTCTCGCTACACCCGAGGACAAGGAACAAGCGCA
ACAGATGAATCAGAAGGACTTTTTGTCACTGATAGTGTCCATCCTGCGCAGTTGGAACGAA
CCCTTGTACCATTTGGTCACCGAAGTCAGGGGGATGCAAGAAGCACCGGAGGCTATACT
GTCAAAGGCCGTAGAAATCGAAGAACAGACGAAGAGACTCCTGGAAGGTATGGAACTC
ATAGTGTCCCAGGTCCACCCAGAGACAAAAGAGAACGAAATATACCCCGTATGGTCTGG
CTTGCCTTCCCTGCAAATGGCAGATGAAGAGAGTCGGTTGAGTGCCTATTACAACCTTCTC
CACTGTCTCAGGAGGGACAGTCACAAGATCGATAACTATCTCAAACTCCTTAAGTGTAGGA
TAATTCATAACAATAACTGTTGA

Fc B (C220S, N297D, K427A, T350V, T366L, K293L, T394W, H435R, Y436F):
TTTAAAGCCGCCACCATGGAGACAGACACACTCCTGCTATGGGTACTGCTGCTCTGGGTT
CCAGGTGAGAGCTGCAGCCTGACTGCATaGGGGCTGGGATaGGCATAAGAATAAAGGT
CTGTGTGGACAGCCTTCTGaTTCAGCCACGACCTCTGTGTATcCTTCTcACCCCAcagGTTCC
ACCGGTgagcccaaatctagcgacaaaactcacacatgcccaccgtgcccaggtaagccagcccaggcctcgccctcca
gctcaaggcgggacaggtgccctagagtagcctgcatccagggacaggccccagccgggtgctgacacgtccacctccatctct
tcctcagcacctgaactcctggggggaccgtcagtcttcctcttccccccaaaacccaaggacaccctcatgatctcccggacccctg
aggtcacatgcgtggtggtggacgtgagccacgaagaccctgaggtcaagttcaactggtacgtggacggcgtggaggtgcata
atgccaagacaaagccgagggaggagcagtacgacagcacgtaccgtgtggtcagcgtcctcaccgtcctgcaccaggactg
gctgaatggcaaggagtacaagtgcaaggtctccaacaaagccctcccagcccccatcgagaaaaccatctccaaagccaaa
ggtgggacccgtggggtgcgagggccacatggacagaggccagctcagcccaccctctgccctgagagtgaccgctgtacca
acctctgtccctacagggcagccccgagaaccacaggtgtacGTGctgcccccatccagggatgagctgaccaagaaccag
gtcagcctgCTGtgcctggtcaaaggcttctatcccagcgacatcgccgtggagtgggagagcaatgggcagccggagaaca
actacCTGaccTGGcctcccgtgctggactccgacggctccttcttcctctacagcaagctcaccgtggacaagagcaggtggc

| | |
|---|---|
| | agcaggggaacgtcttctcatgctccgtgatgcatgaggctctgcacaacCGTTTTacacagaagagcctctccctgtctcccgg<br>ggcaTGA |
| 14. | Fc A (C220S, K447A, N297D, T350V, L351Y, F405A, Y407V, L234A, L235A, V264E, L309D, Q311H, N434S) - GGsGG - PRL (N59D):<br>TTTAAAGCCGCCACCATGGAGACAGACACACTCCTGCTATGGGTACTGCTGCTCTGGGTT<br>CCAGGTGAGAGCTGCAGCCTGACTGCATaGGGGCTGGGATaGGCATAAGAATAAAGGT<br>CTGTGTGGACAGCCTTCTGaTTCAGCCACGACCTCTGTGTATcCTTCTcACCCCAcagGTTCC<br>ACCGGTgagcccaaatctagcgacaaaactcacacatgcccaccgtgcccaggtaagccagcccaggcctcgccctcca<br>gctcaaggcgggacaggtgccctagagtagcctgcatccagggacaggccccagccgggtgctgacacgtccacctccatctct<br>tcctcagcacctgaaGCGGCGgggggaccgtcagtcttcctcttccccccaaaacccaaggacaccctcatgatctcccgga<br>cccctgaggtcacatgcgtggtgGAAgacgtgagccacgaagaccctgaggtcaagttcaactggtacgtggacggcgtgga<br>ggtgcataatgccaagacaaagccgagggaggagcagtacgacagcacgtaccgtgtggtcagcgtcctcaccgtcGATca<br>cCATgactggctgaatggcaaggagtacaagtgcaaggtctccaacaaagccctcccagcccccatcgagaaaaccatctcc<br>aaagccaaaggtgggacccgtggggtgcgagggccacatggacagaggccagctcagcccaccctctgccctgagagtgac<br>cgctgtaccaacctctgtccctacagggcagccccgagaaccacaggtgtacGTGTATcccccatccagggatgagctgacc<br>aagaaccaggtcagcctgCTGtgcctggtcaaaggcttctatcccagcgacatcgccgtggagtgggagagcaatgggcagc<br>cggagaacaactacaagaccacgcctcccgtgctggactccgacggctccttcGCGctcGTGagcaagctcaccgtggaca<br>agagcaggtggcagcaggggaacgtcttctcatgctccgtgatgcatgaggctctgcacAGCcactacacacagaagagcctc<br>tccctgtctcccggggcaGGCGGTAGCGGTGGCTTGCCCATCTGTCCCGGCGGGGCTGCCCGA<br>TGCCAGGTGACCCTTCGAGACCTGTTTGACCGCGCCGTCGTCCTGTCCCACTACATCCAT<br>GACCTCTCCTCAGAAATGTTCAGCGAATTCGATAAACGGTATACCCATGGCCGGGGGTTC<br>ATTACCAAGGCCATCAACAGCTGCCACACTTCTTCCCTTGCCACCCCCGAAGACAAGGA<br>GCAAGCCCAACAGATGAATCAAAAAGACTTTCTGAGCCTGATAGTCAGCATATTGCGATCC<br>TGGAATGAGCCTCTGTATCATCTGGTCACGGAAGTACGTGGTATGCAAGAAGCCCCGGA<br>GGCTATCCTATCCAAAGCTGTAGAGATTGAGGAGCAAACCAAACGGCTTCTAGAGGGCA<br>TGGAGCTGATAGTCAGCCAGGTTCATCCTGAAACCAAAGAAAATGAGATCTACCCTGTCT<br>GGTCGGGACTTCCATCCCTGCAGATGGCTGATGAAGAGTCTCGCCTTTCTGCTTATTATAA<br>CCTGCTCCACTGCCTACGCAGGGATTCACATAAAATCGACAATTATCTCAAGCTCCTGAAG<br>TGCCGAATCATCCACAACAACAACTGCTGA<br><br>Fc A (N297D, C220S, K447A, L234A, L235A, V264E, L309D, Q311H, N434S):<br>TTTAAAGCCGCCACCATGGAGACAGACACACTCCTGCTATGGGTACTGCTGCTCTGGGTT<br>CCAGGTGAGAGCTGCAGCCTGACTGCATaGGGGCTGGGATaGGCATAAGAATAAAGGT<br>CTGTGTGGACAGCCTTCTGaTTCAGCCACGACCTCTGTGTATcCTTCTcACCCCAcagGTTCC<br>ACCGGTgagcccaaatctagcgacaaaactcacacatgcccaccgtgcccaggtaagccagcccaggcctcgccctcca<br>gctcaaggcgggacaggtgccctagagtagcctgcatccagggacaggccccagccgggtgctgacacgtccacctccatctct<br>tcctcagcacctgaaGCGGCGgggggaccgtcagtcttcctcttccccccaaaacccaaggacaccctcatgatctcccgga<br>cccctgaggtcacatgcgtggtgGAAgacgtgagccacgaagaccctgaggtcaagttcaactggtacgtggacggcgtgga<br>ggtgcataatgccaagacaaagccgagggaggagcagtacgacagcacgtaccgtgtggtcagcgtcctcaccgtcGATca<br>cCATgactggctgaatggcaaggagtacaagtgcaaggtctccaacaaagccctcccagcccccatcgagaaaaccatctcc<br>aaagccaaaggtgggacccgtggggtgcgagggccacatggacagaggccagctcagcccaccctctgccctgagagtgac<br>cgctgtaccaacctctgtccctacagggcagccccgagaaccacaggtgtacGTGctgcccccatccagggatgagctgacca<br>agaaccaggtcagcctgCTGtgcctggtcaaaggcttctatcccagcgacatcgccgtggagtgggagagcaatgggcagcc<br>ggagaacaactacCTGaccTGGcctcccgtgctggactccgacggctccttcttcctctacagcaagctcaccgtggacaaga<br>gcaggtggcagcaggggaacgtcttctcatgctccgtgatgcatgaggctctgcacAGCcactacacacagaagagcctctccc<br>tgtctcccggggcaTGA |
| 15. | Fc A (C220S, K447A, N297D, T350V, L351Y, F405A, Y407V, L234A, L235A, V264E, L309D, Q311H, N434S) - GGsGG - PRL (N59D): |

| | |
|---|---|
| | TTTAAAGCCGCCACCATGGAGACAGACACACTCCTGCTATGGGTACTGCTGCTCTGGGTT<br>CCAGGTGAGAGCTGCAGCCTGACTGCATaGGGGCTGGGATaGGCATAAGAATAAAGGT<br>CTGTGTGGACAGCCTTCTGaTTCAGCCACGACCTCTGTGTATcCTTCTcACCCCAcagGTTCC<br>ACCGGTgagcccaaatctagcgacaaaactcacacatgcccaccgtgcccaggtaagccagcccaggcctcgccctcca<br>gctcaaggcgggacaggtgccctagagtagcctgcatccagggacaggccccagccgggtgctgacacgtccacctccatctct<br>tcctcagcacctgaaGCGGCGgggggaccgtcagtcttcctcttccccccaaaacccaaggacaccctcatgatctcccgga<br>cccctgaggtcacatgcgtggtgGAAgacgtgagccacgaagaccctgaggtcaagttcaactggtacgtggacggcgtgga<br>ggtgcataatgccaagacaaagccgagggaggagcagtacgacagcacgtaccgtgtggtcagcgtcctcaccgtcGATca<br>cCATgactggctgaatggcaaggagtacaagtgcaaggtctccaacaaagccctcccagcccccatcgagaaaaccatctcc<br>aaagccaaaggtgggacccgtggggtgcgagggccacatggacagaggccagctcagcccaccctctgccctgagagtgac<br>cgctgtaccaacctctgtccctacagggcagccccgagaaccacaggtgtacGTGTATcccccatccagggatgagctgacc<br>aagaaccaggtcagcctgCTGtgcctggtcaaaggcttctatcccagcgacatcgccgtggagtgggagagcaatgggcagc<br>cggagaacaactacaagaccacgcctcccgtgctggactccgacggctccttcGCGctcGTGagcaagctcaccgtggaca<br>agagcaggtggcagcaggggaacgtcttctcatgctccgtgatgcatgaggctctgcacAGCcactacacacagaagagcctc<br>tccctgtctcccggggcaGGCGGTAGCGGTGGCTTGCCCATCTGTCCCGGCGGGGCTGCCCGA<br>TGCCAGGTGACCCTTCGAGACCTGTTTGACCGCGCCGTCGTCCTGTCCCACTACATCCAT<br>GACCTCTCCTCAGAAATGTTCAGCGAATTCGATAAACGGTATACCCATGGCCGGGGGTTC<br>ATTACCAAGGCCATCAACAGCTGCCACACTTCTTCCCTTGCCACCCCCGAAGACAAGGA<br>GCAAGCCCAACAGATGAATCAAAAAGACTTTCTGAGCCTGATAGTCAGCATATTGCGATCC<br>TGGAATGAGCCTCTGTATCATCTGGTCACGGAAGTACGTGGTATGCAAGAAGCCCCGGA<br>GGCTATCCTATCCAAAGCTGTAGAGATTGAGGAGCAAACCAAACGGCTTCTAGAGGGCA<br>TGGAGCTGATAGTCAGCCAGGTTCATCCTGAAACCAAAGAAAATGAGATCTACCCTGTCT<br>GGTCGGGACTTCCATCCCTGCAGATGGCTGATGAAGAGTCTCGCCTTTCTGCTTATTATAA<br>CCTGCTCCACTGCCTACGCAGGGATTCACATAAAATCGACAATTATCTCAAGCTCCTGAAG<br>TGCCGAATCATCCACAACAACAACTGCTGA<br><br>Fc A (N297D, C220S, K447A, L234A, L235A, V264E, L309D, Q311H, N434S, H435R, Y436F):<br>TTTAAAGCCGCCACCATGGAGACAGACACACTCCTGCTATGGGTACTGCTGCTCTGGGTT<br>CCAGGTGAGAGCTGCAGCCTGACTGCATaGGGGCTGGGATaGGCATAAGAATAAAGGT<br>CTGTGTGGACAGCCTTCTGaTTCAGCCACGACCTCTGTGTATcCTTCTcACCCCAcagGTTCC<br>ACCGGTgagcccaaatctagcgacaaaactcacacatgcccaccgtgcccaggtaagccagcccaggcctcgccctcca<br>gctcaaggcgggacaggtgccctagagtagcctgcatccagggacaggccccagccgggtgctgacacgtccacctccatctct<br>tcctcagcacctgaaGCGGCGgggggaccgtcagtcttcctcttccccccaaaacccaaggacaccctcatgatctcccgga<br>cccctgaggtcacatgcgtggtgGAAgacgtgagccacgaagaccctgaggtcaagttcaactggtacgtggacggcgtgga<br>ggtgcataatgccaagacaaagccgagggaggagcagtacgacagcacgtaccgtgtggtcagcgtcctcaccgtcGATca<br>cCATgactggctgaatggcaaggagtacaagtgcaaggtctccaacaaagccctcccagcccccatcgagaaaaccatctcc<br>aaagccaaaggtgggacccgtggggtgcgagggccacatggacagaggccagctcagcccaccctctgccctgagagtgac<br>cgctgtaccaacctctgtccctacagggcagccccgagaaccacaggtgtacGTGctgcccccatccagggatgagctgacca<br>agaaccaggtcagcctgCTGtgcctggtcaaaggcttctatcccagcgacatcgccgtggagtgggagagcaatgggcagcc<br>ggagaacaactacCTGaccTGGcctcccgtgctggactccgacggctccttcttcctctacagcaagctcaccgtggacaaga<br>gcaggtggcagcaggggaacgtcttctcatgctccgtgatgcatgaggctctgcacAGCCGTTTTacacagaagagcctctc<br>cctgtctcccggggcaTGA |
| 16. | Fc A (C220S, K447A, N297D, T350V, L351Y, F405A, Y407V, L234A, L235A, P329G, V264E, L309D, Q311H, N434S) - GGsGG - PRL N59D:<br>TTTAAAGCCGCCACCATGGAGACAGACACACTCCTGCTATGGGTACTGCTGCTCTGGGTT<br>CCAGGTGAGAGCTGCAGCCTGACTGCATaGGGGCTGGGATaGGCATAAGAATAAAGGT<br>CTGTGTGGACAGCCTTCTGaTTCAGCCACGACCTCTGTGTATcCTTCTcACCCCAcagGTTCC<br>ACCGGTgagcccaaatctagcgacaaaactcacacatgcccaccgtgcccaggtaagccagcccaggcctcgccctcca<br>gctcaaggcgggacaggtgccctagagtagcctgcatccagggacaggccccagccgggtgctgacacgtccacctccatctct |

tcctcagcacctgaaGCGGCGgggggaccgtcagtcttcctcttccccccaaaacccaaggacaccctcatgatctcccgga
cccctgaggtcacatgcgtggtgGAAgacgtgagccacgaagaccctgaggtcaagttcaactggtacgtggacggcgtgga
ggtgcataatgccaagacaaagccgagggaggagcagtacgacagcacgtaccgtgtggtcagcgtcctcaccgtcGATca
cCATgactggctgaatggcaaggagtacaagtgcaaggtctccaacaaagccctcGGAgcccccatcgagaaaaccatct
ccaaagccaaaggtgggacccgtggggtgcgagggccacatggacagaggccagctcagcccaccctctgccctgagagtg
accgctgtaccaacctctgtccctacagggcagccccgagaaccacaggtgtacGTGTATcccccatccagggatgagctga
ccaagaaccaggtcagcctgCTGtgcctggtcaaaggcttctatcccagcgacatcgccgtggagtgggagagcaatgggca
gccggagaacaactacaagaccacgcctcccgtgctggactccgacggctccttcGCGctcGTGagcaagctcaccgtgga
caagagcaggtggcagcaggggaacgtcttctcatgctccgtgatgcatgaggctctgcacAGCcactacacacagaagagc
ctctccctgtctcccggggcaGGCGGTAGCGGTGGCTTGCCCATCTGTCCCGGCGGGGCTGCCC
GATGCCAGGTGACCCTTCGAGACCTGTTTGACCGCGCCGTCGTCCTGTCCCACTACATC
CATGACCTCTCCTCAGAAATGTTCAGCGAATTCGATAAACGGTATACCCATGGCCGGGGG
TTCATTACCAAGGCCATCAACAGCTGCCACACTTCTTCCCTTGCCACCCCCGAAGACAAG
GAGCAAGCCCAACAGATGAATCAAAAAGACTTTCTGAGCCTGATAGTCAGCATATTGCGAT
CCTGGAATGAGCCTCTGTATCATCTGGTCACGGAAGTACGTGGTATGCAAGAAGCCCCG
GAGGCTATCCTATCCAAAGCTGTAGAGATTGAGGAGCAAACCAAACGGCTTCTAGAGGG
CATGGAGCTGATAGTCAGCCAGGTTCATCCTGAAACCAAAGAAAATGAGATCTACCCTGT
CTGGTCGGGACTTCCATCCCTGCAGATGGCTGATGAAGAGTCTCGCCTTTCTGCTTATTATA
ACCTGCTCCACTGCCTACGCAGGGATTCACATAAAATCGACAATTATCTCAAGCTCCTGAA
GTGCCGAATCATCCACAACAACAACTGCTGA

Fc B (N297D, C220S, K447A, L234A, L235A, P329G, V264E, L309D, Q311H, N434S, H435R, Y436F):
TTTAAAGCCGCCACCATGGAGACAGACACACTCCTGCTATGGGTACTGCTGCTCTGGGTT
CCAGGTGAGAGCTGCAGCCTGACTGCATaGGGGCTGGGATaGGCATAAGAATAAAGGT
CTGTGTGGACAGCCTTCTGaTTCAGCCACGACCTCTGTGTATcCTTCTcACCCCAcagGTTCC
ACCGGTgagcccaaatctagcgacaaaactcacacatgcccaccgtgcccaggtaagccagcccaggcctcgccctcca
gctcaaggcgggacaggtgccctagagtagcctgcatccagggacaggccccagccgggtgctgacacgtccacctccatctct
tcctcagcacctgaaGCGGCGgggggaccgtcagtcttcctcttccccccaaaacccaaggacaccctcatgatctcccgga
cccctgaggtcacatgcgtggtgGAAgacgtgagccacgaagaccctgaggtcaagttcaactggtacgtggacggcgtgga
ggtgcataatgccaagacaaagccgagggaggagcagtacgacagcacgtaccgtgtggtcagcgtcctcaccgtcGATca
cCATgactggctgaatggcaaggagtacaagtgcaaggtctccaacaaagccctcGGAgcccccatcgagaaaaccatct
ccaaagccaaaggtgggacccgtggggtgcgagggccacatggacagaggccagctcagcccaccctctgccctgagagtg
accgctgtaccaacctctgtccctacagggcagccccgagaaccacaggtgtacGTGctgcccccatccagggatgagctgac
caagaaccaggtcagcctgCTGtgcctggtcaaaggcttctatcccagcgacatcgccgtggagtgggagagcaatgggcag
ccggagaacaactacCTGaccTGGcctcccgtgctggactccgacggctccttcttcctctacagcaagctcaccgtggacaa
gagcaggtggcagcaggggaacgtcttctcatgctccgtgatgcatgaggctctgcacAGCCGTTTTacacagaagagcct
ctccctgtctcccggggcaTGA

17. Fc A (C220S, K447A, N297D, T350V, L351Y, F405A, Y407V, L234A, L235A, M252Y, S254T, T256E) - GGsGG - PRL N59D:
TTTAAAGCCGCCACCATGGAGACAGACACACTCCTGCTATGGGTACTGCTGCTCTGGGTT
CCAGGTGAGAGCTGCAGCCTGACTGCATaGGGGCTGGGATaGGCATAAGAATAAAGGT
CTGTGTGGACAGCCTTCTGaTTCAGCCACGACCTCTGTGTATcCTTCTcACCCCAcagGTTCC
ACCGGTgagcccaaatctagcgacaaaactcacacatgcccaccgtgcccaggtaagccagcccaggcctcgccctcca
gctcaaggcgggacaggtgccctagagtagcctgcatccagggacaggccccagccgggtgctgacacgtccacctccatctct
tcctcagcacctgaaGCCGCTgggggaccgtcagtcttcctcttccccccaaaacccaaggacaccctcTATatcACCcgg
GAAcctgaggtcacatgcgtggtggtggacgtgagccacgaagaccctgaggtcaagttcaactggtacgtggacggcgtgg
aggtgcataatgccaagacaaagccgagggaggagcagtacgacagcacgtaccgtgtggtcagcgtcctcaccgtcctgca
ccaggactggctgaatggcaaggagtacaagtgcaaggtctccaacaaagccctcccagcccccatcgagaaaaccatctcc
aaagccaaaggtgggacccgtggggtgcgagggccacatggacagaggccagctcagcccaccctctgccctgagagtgac

cgctgtaccaacctctgtccctacagggcagccccgagaaccacaggtgtacGTGTATcccccatccagggatgagctgacc
aagaaccaggtcagcctgCTGtgcctggtcaaaggcttctatcccagcgacatcgccgtggagtgggagagcaatgggcagc
cggagaacaactacaagaccacgcctcccgtgctggactccgacggctccttcGCGctcGTGagcaagctcaccgtggaca
agagcaggtggcagcaggggaacgtcttctcatgctccgtgatgcatgaggctctgcacaaccactacacacagaagagcctct
ccctgtctcccgggcaGGCGGTAGCGGTGGCCTCCCGATATGTCCGGGCGGGGCCGCTCG
GTGCCAGGTAACTTTGAGGGACCTGTTTGACCGAGCCGTAGTCCTTTCACACTATATTCAC
GACCTCTCATCTGAGATGTTTTCCGAGTTCGACAAGAGATATACCCACGGTCGCGGGTTTA
TAACTAAGGCAATAAACAGTTGCCATACCTCAAGTCTCGCTACACCCGAGGACAAGGAAC
AAGCGCAACAGATGAATCAGAAGGACTTTTTGTCACTGATAGTGTCCATCCTGCGCAGTTG
GAACGAACCCTTGTACCATTTGGTCACCGAAGTCAGGGGGATGCAAGAAGCACCGGAG
GCTATACTGTCAAAGGCCGTAGAAATCGAAGAACAGACGAAGAGACTCCTGGAAGGTAT
GGAACTCATAGTGTCCCAGGTCCACCCAGAGACAAAAGAGAACGAAATATACCCCGTAT
GGTCTGGCTTGCCTTCCCTGCAAATGGCAGATGAAGAGAGTCGGTTGAGTGCCTATTACA
ACCTTCTCCACTGTCTCAGGAGGGACAGTCACAAGATCGATAACTATCTCAAACTCCTTAA
GTGTAGGATAATTCATAACAATAACTGTTGA

Fc B (N297D, K447A, H435R, Y436F, L234A, L235A, M252Y, S254T, T256E):
TTTAAAGCCGCCACCATGGAGACAGACACACTCCTGCTATGGGTACTGCTGCTCTGGGTT
CCAGGTGAGAGCTGCAGCCTGACTGCATaGGGGCTGGGATaGGCATAAGAATAAAGGT
CTGTGTGGACAGCCTTCTGaTTCAGCCACGACCTCTGTGTATcCTTCTcACCCCAcagGTTCC
ACCGGTgagcccaaatctagcgacaaaactcacacatgcccaccgtgcccaggtaagccagcccaggcctcgccctcca
gctcaaggcgggacaggtgccctagagtagcctgcatccagggacaggccccagccgggtgctgacacgtccacctccatctct
tcctcagcacctgaaGCCGCTgggggaccgtcagtcttcctcttccccccaaaacccaaggacaccctcTATatcACCcgg
GAAcctgaggtcacatgcgtggtggtggacgtgagccacgaagaccctgaggtcaagttcaactggtacgtggacggcgtgg
aggtgcataatgccaagacaaagccgagggaggagcagtacgacagcacgtaccgtgtggtcagcgtcctcaccgtcctgca
ccaggactggctgaatggcaaggagtacaagtgcaaggtctccaacaaagccctcccagcccccatcgagaaaaccatctcc
aaagccaaaggtgggacccgtggggtgcgagggccacatggacagaggccagctcagcccaccctctgccctgagagtgac
cgctgtaccaacctctgtccctacagggcagccccgagaaccacaggtgtacGTGctgcccccatccagggatgagctgacca
agaaccaggtcagcctgCTGtgcctggtcaaaggcttctatcccagcgacatcgccgtggagtgggagagcaatgggcagcc
ggagaacaactacCTGaccTGGcctcccgtgctggactccgacggctccttcttcctctacagcaagctcaccgtggacaaga
gcaggtggcagcaggggaacgtcttctcatgctccgtgatgcatgaggctctgcacaacCGTTTTacacagaagagcctctccc
tgtctcccgggcaTGA

18. PRL (N59D) - Fc Knob (C220S, N297D, K427A, T366W):
TTTAAAGCCGCCACCATGGAGACAGACACACTCCTGCTATGGGTACTGCTGCTCTGGGTT
CCAGGTGAGAGCTGCAGCCTGACTGCATaGGGGCTGGGATaGGCATAAGAATAAAGGT
CTGTGTGGACAGCCTTCTGaTTCAGCCACGACCTCTGTGTATcCTTCTcACCCCAcagGTTCC
ACCGGTCTCCCGATATGTCCGGGCGGGGCCGCTCGGTGCCAGGTAACTTTGAGGGAC
CTGTTTGACCGAGCCGTAGTCCTTTCACACTATATTCACGACCTCTCATCTGAGATGTTTTCC
GAGTTCGACAAGAGATATACCCACGGTCGCGGGTTTATAACTAAGGCAATAAACAGTTGC
CATACCTCAAGTCTCGCTACACCCGAGGACAAGGAACAAGCGCAACAGATGAATCAGAA
GGACTTTTTGTCACTGATAGTGTCCATCCTGCGCAGTTGGAACGAACCCTTGTACCATTTGG
TCACCGAAGTCAGGGGGATGCAAGAAGCACCGGAGGCTATACTGTCAAAGGCCGTAG
AAATCGAAGAACAGACGAAGAGACTCCTGGAAGGTATGGAACTCATAGTGTCCCAGGTC
CACCCAGAGACAAAAGAGAACGAAATATACCCCGTATGGTCTGGCTTGCCTTCCCTGCAA
ATGGCAGATGAAGAGAGTCGGTTGAGTGCCTATTACAACCTTCTCCACTGTCTCAGGAGG
GACAGTCACAAGATCGATAACTATCTCAAACTCCTTAAGTGTAGGATAATTCATAACAATAAC
TGTgagcccaaatctagcgacaaaactcacacatgcccaccgtgcccaggtaagccagcccaggcctcgccctccagctcaa
ggcgggacaggtgccctagagtagcctgcatccagggacaggccccagccgggtgctgacacgtccacctccatctcttcctcag
cacctgaactcctggggggaccgtcagtcttcctcttccccccaaaacccaaggacaccctcatgatctcccggacccctgaggtca
catgcgtggtggtggacgtgagccacgaagaccctgaggtcaagttcaactggtacgtggacggcgtggaggtgcataatgcca

agacaaagccgagggaggagcagtacgacagcacgtaccgtgtggtcagcgtcctcaccgtcctgcaccaggactggctgaa
tggcaaggagtacaagtgcaaggtctccaacaaagccctcccagcccccatcgagaaaaccatctccaaagccaaaggtgg
gacccgtggggtgcgagggccacatggacagaggccagctcagcccaccctctgccctgagagtgaccgctgtaccaacctct
gtccctacagggcagccccgagaaccacaggtgtacaccctgcccccatccagggatgagctgaccaagaaccaggtcagcc
tgtggtgcctggtcaaaggcttctatcccagcgacatcgccgtggagtgggagagcaatgggcagccggagaacaactacaag
accacgcctcccgtgctggactccgacggctccttcttcctctacagcaagctcaccgtggacaagagcaggtggcagcagggg
aacgtcttctcatgctccgtgatgcatgaggctctgcacaaccactacacacagaagagcctctccctgtctccggggcaGAAC
AAAAACTCATCTCAGAAGAGGATCTGAATAGCGCCGTCGACCATCATCATCATCATCATTG
A

Fc Hole (C220S, N297D, K427A, T366S, L368A, Y407V):
TTTAAAGCCGCCACCATGGAGACAGACACACTCCTGCTATGGGTACTGCTGCTCTGGGTT
CCAGGTGAGAGCTGCAGCCTGACTGCATaGGGGCTGGGATaGGCATAAGAATAAAGGT
CTGTGTGGACAGCCTTCTGaTTCAGCCACGACCTCTGTGTATcCTTCTcACCCCAcagGTTCC
ACCGGTgagcccaaatctagcgacaaaactcacacatgcccaccgtgcccaggtaagccagcccaggcctcgccctcca
gctcaaggcgggacaggtgccctagagtagcctgcatccagggacaggccccagccgggtgctgacacgtccacctccatctct
tcctcagcacctgaactcctggggggaccgtcagtcttcctcttccccccaaaacccaaggacaccctcatgatctcccggacccctg
aggtcacatgcgtggtggtggacgtgagccacgaagaccctgaggtcaagttcaactggtacgtggacggcgtggaggtgcata
atgccaagacaaagccgagggaggagcagtacgacagcacgtaccgtgtggtcagcgtcctcaccgtcctgcaccaggactg
gctgaatggcaaggagtacaagtgcaaggtctccaacaaagccctcccagcccccatcgagaaaaccatctccaaagccaaa
ggtgggacccgtggggtgcgagggccacatggacagaggccagctcagcccaccctctgccctgagagtgaccgctgtacca
acctctgtccctacagggcagccccgagaaccacaggtgtacaccctgcccccatccagggatgagctgaccaagaaccaggt
cagcctgagctgcgccgtcaaaggcttctatcccagcgacatcgccgtggagtgggagagcaatgggcagccggagaacaac
tacaagaccacgcctcccgtgctggactccgacggctccttcttcctcgtcagcaagctcaccgtggacaagagcaggtggcagc
aggggaacgtcttctcatgctccgtgatgcatgaggctctgcacaacCGTTTTacacagaagagcctctccctgtctccggggc
aTGA

19. PRL (N59D) - GGsGG - Fc Knob (C220S, N297D, K427A, T366W):
TTTAAAGCCGCCACCATGGAGACAGACACACTCCTGCTATGGGTACTGCTGCTCTGGGTT
CCAGGTGAGAGCTGCAGCCTGACTGCATaGGGGCTGGGATaGGCATAAGAATAAAGGT
CTGTGTGGACAGCCTTCTGaTTCAGCCACGACCTCTGTGTATcCTTCTcACCCCAcagGTTCC
ACCGGTCTCCCGATATGTCCGGGCGGGGCCGCTCGGTGCCAGGTAACTTTGAGGGAC
CTGTTTGACCGAGCCGTAGTCCTTTCACACTATATTCACGACCTCTCATCTGAGATGTTTTCC
GAGTTCGACAAGAGATATACCCACGGTCGCGGGTTTATAACTAAGGCAATAAACAGTTGC
CATACCTCAAGTCTCGCTACACCCGAGGACAAGGAACAAGCGCAACAGATGAATCAGAA
GGACTTTTTGTCACTGATAGTGTCCATCCTGCGCAGTTGGAACGAACCCTTGTACCATTTGG
TCACCGAAGTCAGGGGGATGCAAGAAGCACCGGAGGCTATACTGTCAAAGGCCGTAG
AAATCGAAGAACAGACGAAGAGACTCCTGGAAGGTATGGAACTCATAGTGTCCCAGGTC
CACCCAGAGACAAAAGAGAACGAAATATACCCCGTATGGTCTGGCTTGCCTTCCCTGCAA
ATGGCAGATGAAGAGAGTCGGTTGAGTGCCTATTACAACCTTCTCCACTGTCTCAGGAGG
GACAGTCACAAGATCGATAACTATCTCAAACTCCTTAAGTGTAGGATAATTCATAACAATAAC
TGTGGCGGTAGCGGTGGCgagcccaaatctagcgacaaaactcacacatgcccaccgtgcccaggtaagccag
cccaggcctcgccctccagctcaaggcgggacaggtgccctagagtagcctgcatccagggacaggccccagccgggtgctg
acacgtccacctccatctcttcctcagcacctgaactcctggggggaccgtcagtcttcctcttccccccaaaacccaaggacaccctc
atgatctcccggacccctgaggtcacatgcgtggtggtggacgtgagccacgaagaccctgaggtcaagttcaactggtacgtgg
acggcgtggaggtgcataatgccaagacaaagccgagggaggagcagtacgacagcacgtaccgtgtggtcagcgtcctca
ccgtcctgcaccaggactggctgaatggcaaggagtacaagtgcaaggtctccaacaaagccctcccagcccccatcgagaa
aaccatctccaaagccaaaggtgggacccgtggggtgcgagggccacatggacagaggccagctcagcccaccctctgccct
gagagtgaccgctgtaccaacctctgtccctacagggcagccccgagaaccacaggtgtacaccctgcccccatccagggatg
agctgaccaagaaccaggtcagcctgtggtgcctggtcaaaggcttctatcccagcgacatcgccgtggagtgggagagcaatg
ggcagccggagaacaactacaagaccacgcctcccgtgctggactccgacggctccttcttcctctacagcaagctcaccgtgga

caagagcaggtggcagcaggggaacgtcttctcatgctccgtgatgcatgaggctctgcacaaccactacacacagaagagcct
ctccctgtctcccggggcaTGA

Fc Hole (C220S, N297D, K427A, T366S, L368A, Y407V):
TTTAAAGCCGCCACCATGGAGACAGACACACTCCTGCTATGGGTACTGCTGCTCTGGGTT
CCAGGTGAGAGCTGCAGCCTGACTGCATaGGGGCTGGGATaGGCATAAGAATAAAGGT
CTGTGTGGACAGCCTTCTGaTTCAGCCACGACCTCTGTGTATcCTTCTcACCCCAcagGTTCC
ACCGGTgagcccaaatctagcgacaaaactcacacatgcccaccgtgcccaggtaagccagcccaggcctcgccctcca
gctcaaggcgggacaggtgccctagagtagcctgcatccagggacaggccccagccgggtgctgacacgtccacctccatctct
tcctcagcacctgaactcctggggggaccgtcagtcttcctcttccccccaaaacccaaggacaccctcatgatctcccggacccctg
aggtcacatgcgtggtggtggacgtgagccacgaagaccctgaggtcaagttcaactggtacgtggacggcgtggaggtgcata
atgccaagacaaagccgagggaggagcagtacgacagcacgtaccgtgtggtcagcgtcctcaccgtcctgcaccaggactg
gctgaatggcaaggagtacaagtgcaaggtctccaacaaagccctcccagcccccatcgagaaaaccatctccaaagccaaa
ggtgggacccgtggggtgcgagggccacatggacagaggccagctcagcccaccctctgccctgagagtgaccgctgtacca
acctctgtccctacagggcagccccgagaaccacaggtgtacaccctgcccccatccagggatgagctgaccaagaaccaggt
cagcctgagctgcgccgtcaaaggcttctatcccagcgacatcgccgtggagtgggagagcaatgggcagccggagaacaac
tacaagaccacgcctcccgtgctggactccgacggctccttcttcctcgtcagcaagctcaccgtggacaagagcaggtggcagc
aggggaacgtcttctcatgctccgtgatgcatgaggctctgcacaacCGTTTTacacagaagagcctctccctgtctcccggggc
aTGA

20.

PRL (N59D, C191S, C199S) - Fc Knob (C220S, N297D, K427A, T366W):
TTTAAAGCCGCCACCATGGAGACAGACACACTCCTGCTATGGGTACTGCTGCTCTGGGTT
CCAGGTGAGAGCTGCAGCCTGACTGCATaGGGGCTGGGATaGGCATAAGAATAAAGGT
CTGTGTGGACAGCCTTCTGaTTCAGCCACGACCTCTGTGTATcCTTCTcACCCCAcagGTTCC
ACCGGTCTCCCGATATGTCCGGGCGGGGCCGCTCGGTGCCAGGTAACTTTGAGGGAC
CTGTTTGACCGAGCCGTAGTCCTTTCACACTATATTCACGACCTCTCATCTGAGATGTTTTCC
GAGTTCGACAAGAGATATACCCACGGTCGCGGGTTTATAACTAAGGCAATAAACAGTTGC
CATACCTCAAGTCTCGCTACACCCGAGGACAAGGAACAAGCGCAACAGATGAATCAGAA
GGACTTTTTGTCACTGATAGTGTCCATCCTGCGCAGTTGGAACGAACCCTTGTACCATTTGG
TCACCGAAGTCAGGGGGATGCAAGAAGCACCGGAGGCTATACTGTCAAAGGCCGTAG
AAATCGAAGAACAGACGAAGAGACTCCTGGAAGGTATGGAACTCATAGTGTCCCAGGTC
CACCCAGAGACAAAAGAGAACGAAATATACCCCGTATGGTCTGGCTTGCCTTCCCTGCAA
ATGGCAGATGAAGAGAGTCGGTTGAGTGCCTATTACAACCTTCTCCACTGTCTCAGGAGG
GACAGTCACAAGATCGATAACTATCTCAAACTCCTTAAGAGCAGGATAATTCATAACAATAA
CAGCgagcccaaatctagcgacaaaactcacacatgcccaccgtgcccaggtaagccagcccaggcctcgccctccagctc
aaggcgggacaggtgccctagagtagcctgcatccagggacaggccccagccgggtgctgacacgtccacctccatctcttcctc
agcacctgaactcctggggggaccgtcagtcttcctcttccccccaaaacccaaggacaccctcatgatctcccggacccctgaggt
cacatgcgtggtggtggacgtgagccacgaagaccctgaggtcaagttcaactggtacgtggacggcgtggaggtgcataatgc
caagacaaagccgagggaggagcagtacgacagcacgtaccgtgtggtcagcgtcctcaccgtcctgcaccaggactggctg
aatggcaaggagtacaagtgcaaggtctccaacaaagccctcccagcccccatcgagaaaaccatctccaaagccaaaggt
gggacccgtggggtgcgagggccacatggacagaggccagctcagcccaccctctgccctgagagtgaccgctgtaccaacc
tctgtccctacagggcagccccgagaaccacaggtgtacaccctgcccccatccagggatgagctgaccaagaaccaggtcag
cctgtggtgcctggtcaaaggcttctatcccagcgacatcgccgtggagtgggagagcaatgggcagccggagaacaactaca
agaccacgcctcccgtgctggactccgacggctccttcttcctctacagcaagctcaccgtggacaagagcaggtggcagcagg
ggaacgtcttctcatgctccgtgatgcatgaggctctgcacaaccactacacacagaagagcctctccctgtctcccggggcaGAA
CAAAAACTCATCTCAGAAGAGGATCTGAATAGCGCCGTCGACCATCATCATCATCATCATT
GA

Fc Hole (C220S, N297D, K427A, T366S, L368A, Y407V):
TTTAAAGCCGCCACCATGGAGACAGACACACTCCTGCTATGGGTACTGCTGCTCTGGGTT
CCAGGTGAGAGCTGCAGCCTGACTGCATaGGGGCTGGGATaGGCATAAGAATAAAGGT

CTGTGTGGACAGCCTTCTGaTTCAGCCACGACCTCTGTGTATcCTTCTcACCCCAcagGTTCC
ACCGGTgagcccaaatctagcgacaaaactcacacatgcccaccgtgcccaggtaagccagcccaggcctcgccctcca
gctcaaggcgggacaggtgccctagagtagcctgcatccagggacaggccccagccgggtgctgacacgtccacctccatctct
tcctcagcacctgaactcctggggggaccgtcagtcttcctcttccccccaaaacccaaggacaccctcatgatctcccggacccctg
aggtcacatgcgtggtggtggacgtgagccacgaagaccctgaggtcaagttcaactggtacgtggacggcgtggaggtgcata
atgccaagacaaagccgagggaggagcagtacgacagcacgtaccgtgtggtcagcgtcctcaccgtcctgcaccaggactg
gctgaatggcaaggagtacaagtgcaaggtctccaacaaagccctcccagcccccatcgagaaaaccatctccaaagccaaa
ggtgggacccgtggggtgcgagggccacatggacagaggccagctcagcccaccctctgccctgagagtgaccgctgtacca
acctctgtccctacagggcagccccgagaaccacaggtgtacaccctgcccccatccagggatgagctgaccaagaaccaggt
cagcctgagctgcgccgtcaaaggcttctatcccagcgacatcgccgtggagtgggagagcaatgggcagccggagaacaac
tacaagaccacgcctcccgtgctggactccgacggctccttcttcctcgtcagcaagctcaccgtggacaagagcaggtggcagc
aggggaacgtcttctcatgctccgtgatgcatgaggctctgcacaacCGTTTTacacagaagagcctctccctgtctcccggggc
aTGA

21. PRL (N59D, C191S, C199S) - Fc Knob (C220S, C226S, C229S, N297D, K427A, T366W):

TTTAAAGCCGCCACCATGGAGACAGACACACTCCTGCTATGGGTACTGCTGCTCTGGGTT
CCAGGTGAGAGCTGCAGCCTGACTGCATaGGGGCTGGGATaGGCATAAGAATAAAGGT
CTGTGTGGACAGCCTTCTGaTTCAGCCACGACCTCTGTGTATcCTTCTcACCCCAcagGTTCC
ACCGGTCTCCCGATATGTCCGGGCGGGGCCGCTCGGTGCCAGGTAACTTTGAGGGAC
CTGTTTGACCGAGCCGTAGTCCTTTCACACTATATTCACGACCTCTCATCTGAGATGTTTTCC
GAGTTCGACAAGAGATATACCCACGGTCGCGGGTTTATAACTAAGGCAATAAACAGTTGC
CATACCTCAAGTCTCGCTACACCCGAGGACAAGGAACAAGCGCAACAGATGAATCAGAA
GGACTTTTTGTCACTGATAGTGTCCATCCTGCGCAGTTGGAACGAACCCTTGTACCATTTGG
TCACCGAAGTCAGGGGGATGCAAGAAGCACCGGAGGCTATACTGTCAAAGGCCGTAG
AAATCGAAGAACAGACGAAGAGACTCCTGGAAGGTATGGAACTCATAGTGTCCCAGGTC
CACCCAGAGACAAAAGAGAACGAAATATACCCCGTATGGTCTGGCTTGCCTTCCCTGCAA
ATGGCAGATGAAGAGAGTCGGTTGAGTGCCTATTACAACCTTCTCCACTGTCTCAGGAGG
GACAGTCACAAGATCGATAACTATCTCAAACTCCTTAAGAGCAGGATAATTCATAACAATAA
CAGCgagcccaaatctagcgacaaaactcacacaAGCccaccgAGCccaggtaagccagcccaggcctcgccctcc
agctcaaggcgggacaggtgccctagagtagcctgcatccagggacaggccccagccgggtgctgacacgtccacctccatct
cttcctcagcacctgaactcctggggggaccgtcagtcttcctcttccccccaaaacccaaggacaccctcatgatctcccggacccc
tgaggtcacatgcgtggtggtggacgtgagccacgaagaccctgaggtcaagttcaactggtacgtggacggcgtggaggtgca
taatgccaagacaaagccgagggaggagcagtacgacagcacgtaccgtgtggtcagcgtcctcaccgtcctgcaccaggac
tggctgaatggcaaggagtacaagtgcaaggtctccaacaaagccctcccagcccccatcgagaaaaccatctccaaagcca
aaggtgggacccgtggggtgcgagggccacatggacagaggccagctcagcccaccctctgccctgagagtgaccgctgtac
caacctctgtccctacagggcagccccgagaaccacaggtgtacaccctgcccccatccagggatgagctgaccaagaacca
ggtcagcctgtggtgcctggtcaaaggcttctatcccagcgacatcgccgtggagtgggagagcaatgggcagccggagaaca
actacaagaccacgcctcccgtgctggactccgacggctccttcttcctctacagcaagctcaccgtggacaagagcaggtggca
gcaggggaacgtcttctcatgctccgtgatgcatgaggctctgcacaaccactacacacagaagagcctctccctgtctcccgggg
caGAACAAAAACTCATCTCAGAAGAGGATCTGAATAGCGCCGTCGACCATCATCATCATCA
TCATTGA

Fc Hole (C220S, N297D, K427A, T366S, L368A, Y407V):

TTTAAAGCCGCCACCATGGAGACAGACACACTCCTGCTATGGGTACTGCTGCTCTGGGTT
CCAGGTGAGAGCTGCAGCCTGACTGCATaGGGGCTGGGATaGGCATAAGAATAAAGGT
CTGTGTGGACAGCCTTCTGaTTCAGCCACGACCTCTGTGTATcCTTCTcACCCCAcagGTTCC
ACCGGTgagcccaaatctagcgacaaaactcacacatgcccaccgtgcccaggtaagccagcccaggcctcgccctcca
gctcaaggcgggacaggtgccctagagtagcctgcatccagggacaggccccagccgggtgctgacacgtccacctccatctct
tcctcagcacctgaactcctggggggaccgtcagtcttcctcttccccccaaaacccaaggacaccctcatgatctcccggacccctg
aggtcacatgcgtggtggtggacgtgagccacgaagaccctgaggtcaagttcaactggtacgtggacggcgtggaggtgcata

atgccaagacaaagccgagggaggagcagtacgacagcacgtaccgtgtggtcagcgtcctcaccgtcctgcaccaggactg
gctgaatggcaaggagtacaagtgcaaggtctccaacaaagccctcccagcccccatcgagaaaaccatctccaaagccaaa
ggtgggacccgtggggtgcgagggccacatggacagaggccagctcagcccaccctctgccctgagagtgaccgctgtacca
acctctgtccctacagggcagccccgagaaccacaggtgtacaccctgcccccatccagggatgagctgaccaagaaccaggt
cagcctgagctgcgccgtcaaaggcttctatcccagcgacatcgccgtggagtgggagagcaatgggcagccggagaacaac
tacaagaccacgcctcccgtgctggactccgacggctccttcttcctcgtcagcaagctcaccgtggacaagagcaggtggcagc
aggggaacgtcttctcatgctccgtgatgcatgaggctctgcacaacCGTTTTacacagaagagcctctccctgtctcccggggc
aTGA

22. PRL (N59D) - GGsGG- Fc Knob (C220S, K447A, N297D, T366W, L234A, L235A, V264E, L309D, Q311H, N434S):

TTTAAAGCCGCCACCATGGAGACAGACACACTCCTGCTATGGGTACTGCTGCTCTGGGTT
CCAGGTGAGAGCTGCAGCCTGACTGCATaGGGGCTGGGATaGGCATAAGAATAAAGGT
CTGTGTGGACAGCCTTCTGaTTCAGCCACGACCTCTGTGTATcCTTCTcACCCCAcagGTTCC
ACCGGTTTGCCCATCTGTCCCGGCGGGGCTGCCCGATGCCAGGTGACCCTTCGAGAC
CTGTTTGACCGCGCCGTCGTCCTGTCCCACTACATCCATGACCTCTCCTCAGAAATGTTCA
GCGAATTCGATAAACGGTATACCCATGGCCGGGGGTTCATTACCAAGGCCATCAACAGC
TGCCACACTTCTTCCCTTGCCACCCCCGAAGACAAGGAGCAAGCCCAACAGATGAATCA
AAAAGACTTTCTGAGCCTGATAGTCAGCATATTGCGATCCTGGAATGAGCCTCTGTATCATC
TGGTCACGGAAGTACGTGGTATGCAAGAAGCCCCGGAGGCTATCCTATCCAAAGCTGTA
GAGATTGAGGAGCAAACCAAACGGCTTCTAGAGGGCATGGAGCTGATAGTCAGCCAGG
TTCATCCTGAAACCAAAGAAAATGAGATCTACCCTGTCTGGTCGGGACTTCCATCCCTGCA
GATGGCTGATGAAGAGTCTCGCCTTTCTGCTTATTATAACCTGCTCCACTGCCTACGCAGG
GATTCACATAAAATCGACAATTATCTCAAGCTCCTGAAGTGCCGAATCATCCACAACAACAA
CTGCGGCGGTAGCGGTGGCgagcccaaatctagcgacaaaactcacacatgcccaccgtgcccaggtaagcc
agcccaggcctcgccctccagctcaaggcgggacaggtgccctagagtagcctgcatccagggacaggccccagccgggtg
ctgacacgtccacctccatctcttcctcagcacctgaaGCGGCGgggggaccgtcagtcttcctcttccccccaaaacccaagg
acaccctcatgatctcccggacccctgaggtcacatgcgtggtgGAAgacgtgagccacgaagaccctgaggtcaagttcaact
ggtacgtggacggcgtggaggtgcataatgccaagacaaagccgagggaggagcagtacgacagcacgtaccgtgtggtca
gcgtcctcaccgtcGATcacCATgactggctgaatggcaaggagtacaagtgcaaggtctccaacaaagccctcccagcccc
catcgagaaaaccatctccaaagccaaaggtgggacccgtggggtgcgagggccacatggacagaggccagctcagccca
ccctctgccctgagagtgaccgctgtaccaacctctgtccctacagggcagccccgagaaccacaggtgtacaccctgcccccatc
cagggatgagctgaccaagaaccaggtcagcctgTGGtgcctggtcaaaggcttctatcccagcgacatcgccgtggagtggg
agagcaatgggcagccggagaacaactacaagaccacgcctcccgtgctggactccgacggctccttcttcctctacagcaagct
caccgtggacaagagcaggtggcagcaggggaacgtcttctcatgctccgtgatgcatgaggctctgcacAGCcactacacac
agaagagcctctccctgtctcccggggcaTGA

Fc Hole (C220S, N297D, K447A, T366S, L368A, Y407V, H435R, Y436F, L234A, L235A, V264E, L309D, Q311H, N434S):

TTTAAAGCCGCCACCATGGAGACAGACACACTCCTGCTATGGGTACTGCTGCTCTGGGTT
CCAGGTGAGAGCTGCAGCCTGACTGCATaGGGGCTGGGATaGGCATAAGAATAAAGGT
CTGTGTGGACAGCCTTCTGaTTCAGCCACGACCTCTGTGTATcCTTCTcACCCCAcagGTTCC
ACCGGTgagcccaaatctagcgacaaaactcacacatgcccaccgtgcccaggtaagccagcccaggcctcgccctcca
gctcaaggcgggacaggtgccctagagtagcctgcatccagggacaggccccagccgggtgctgacacgtccacctccatctct
tcctcagcacctgaaGCGGCGgggggaccgtcagtcttcctcttccccccaaaacccaaggacaccctcatgatctcccgga
cccctgaggtcacatgcgtggtgGAAgacgtgagccacgaagaccctgaggtcaagttcaactggtacgtggacggcgtgga
ggtgcataatgccaagacaaagccgagggaggagcagtacgacagcacgtaccgtgtggtcagcgtcctcaccgtcGATca
cCATgactggctgaatggcaaggagtacaagtgcaaggtctccaacaaagccctcccagcccccatcgagaaaaccatctcc
aaagccaaaggtgggacccgtggggtgcgagggccacatggacagaggccagctcagcccaccctctgccctgagagtgac
cgctgtaccaacctctgtccctacagggcagccccgagaaccacaggtgtacaccctgcccccatccagggatgagctgaccaa
gaaccaggtcagcctgagctgcgccgtcaaaggcttctatcccagcgacatcgccgtggagtgggagagcaatgggcagccg

gagaacaactacaagaccacgcctcccgtgctggactccgacggctccttcttcctcgtcagcaagctcaccgtggacaagagca
ggtggcagcaggggaacgtcttctcatgctccgtgatgcatgaggctctgcacAGCCGTTTTacacagaagagcctctccctgt
ctcccggggcaTGA

23. PRL N59D - GGsGG- Fc Knob (C220S, K447A, N297D, T366W, L234A, L235A, P329G, V264E, L309D, Q311H, N434S):

TTTAAAGCCGCCACCATGGAGACAGACACACTCCTGCTATGGGTACTGCTGCTCTGGGTT
CCAGGTGAGAGCTGCAGCCTGACTGCATaGGGGCTGGGATaGGCATAAGAATAAAGGT
CTGTGTGGACAGCCTTCTGaTTCAGCCACGACCTCTGTGTATcCTTCTcACCCCAcagGTTCC
ACCGGTTTGCCCATCTGTCCCGGCGGGGCTGCCCGATGCCAGGTGACCCTTCGAGAC
CTGTTTGACCGCGCCGTCGTCCTGTCCCACTACATCCATGACCTCTCCTCAGAAATGTTCA
GCGAATTCGATAAACGGTATACCCATGGCCGGGGGTTCATTACCAAGGCCATCAACAGC
TGCCACACTTCTTCCCTTGCCACCCCCGAAGACAAGGAGCAAGCCCAACAGATGAATCA
AAAAGACTTTCTGAGCCTGATAGTCAGCATATTGCGATCCTGGAATGAGCCTCTGTATCATC
TGGTCACGGAAGTACGTGGTATGCAAGAAGCCCCGGAGGCTATCCTATCCAAAGCTGTA
GAGATTGAGGAGCAAACCAAACGGCTTCTAGAGGGCATGGAGCTGATAGTCAGCCAGG
TTCATCCTGAAACCAAAGAAAATGAGATCTACCCTGTCTGGTCGGGACTTCCATCCCTGCA
GATGGCTGATGAAGAGTCTCGCCTTTCTGCTTATTATAACCTGCTCCACTGCCTACGCAGG
GATTCACATAAAATCGACAATTATCTCAAGCTCCTGAAGTGCCGAATCATCCACAACAACAA
CTGCGGCGGTAGCGGTGGCgagcccaaatctagcgacaaaactcacacatgcccaccgtgcccaggtaagcc
agcccaggcctcgccctccagctcaaggcgggacaggtgccctagagtagcctgcatccagggacaggccccagccgggtg
ctgacacgtccacctccatctcttcctcagcacctgaaGCGGCGgggggaccgtcagtcttcctcttccccccaaaacccaagg
acaccctcatgatctcccggacccctgaggtcacatgcgtggtgGAAgacgtgagccacgaagaccctgaggtcaagttcaact
ggtacgtggacggcgtggaggtgcataatgccaagacaaagccgagggaggagcagtacgacagcacgtaccgtgtggtca
gcgtcctcaccgtcGATcacCATgactggctgaatggcaaggagtacaagtgcaaggtctccaacaaagccctcGGAgcc
cccatcgagaaaaccatctccaaagccaaaggtgggacccgtggggtgcgagggccacatggacagaggccagctcagcc
caccctctgccctgagagtgaccgctgtaccaacctctgtccctacagggcagccccgagaaccacaggtgtacaccctgcccc
atccagggatgagctgaccaagaaccaggtcagcctgTGGtgcctggtcaaaggcttctatcccagcgacatcgccgtggagt
gggagagcaatgggcagccggagaacaactacaagaccacgcctcccgtgctggactccgacggctccttcttcctctacagca
agctcaccgtggacaagagcaggtggcagcaggggaacgtcttctcatgctccgtgatgcatgaggctctgcacAGCcactac
acacagaagagcctctccctgtctcccggggcaTGA

Fc Hole (C220S, N297D, K447A, T366S, L368A, Y407V, H435R, Y436F, L234A, L235A, P329G, V264E, L309D, Q311H, N434S):

TTTAAAGCCGCCACCATGGAGACAGACACACTCCTGCTATGGGTACTGCTGCTCTGGGTT
CCAGGTGAGAGCTGCAGCCTGACTGCATaGGGGCTGGGATaGGCATAAGAATAAAGGT
CTGTGTGGACAGCCTTCTGaTTCAGCCACGACCTCTGTGTATcCTTCTcACCCCAcagGTTCC
ACCGGTgagcccaaatctagcgacaaaactcacacatgcccaccgtgcccaggtaagccagcccaggcctcgccctcca
gctcaaggcgggacaggtgccctagagtagcctgcatccagggacaggccccagccgggtgctgacacgtccacctccatctct
tcctcagcacctgaaGCGGCGgggggaccgtcagtcttcctcttccccccaaaacccaaggacaccctcatgatctcccgga
cccctgaggtcacatgcgtggtgGAAgacgtgagccacgaagaccctgaggtcaagttcaactggtacgtggacggcgtgga
ggtgcataatgccaagacaaagccgagggaggagcagtacgacagcacgtaccgtgtggtcagcgtcctcaccgtcGATca
cCATgactggctgaatggcaaggagtacaagtgcaaggtctccaacaaagccctcGGAgcccccatcgagaaaaccatct
ccaaagccaaaggtgggacccgtggggtgcgagggccacatggacagaggccagctcagcccaccctctgccctgagagtg
accgctgtaccaacctctgtccctacagggcagccccgagaaccacaggtgtacaccctgcccccatccagggatgagctgacc
aagaaccaggtcagcctgagctgcgccgtcaaaggcttctatcccagcgacatcgccgtggagtgggagagcaatgggcagc
cggagaacaactacaagaccacgcctcccgtgctggactccgacggctccttcttcctcgtcagcaagctcaccgtggacaagag
caggtggcagcaggggaacgtcttctcatgctccgtgatgcatgaggctctgcacAGCCGTTTTacacagaagagcctctccc
tgtctcccggggcaTGA

24. PRL N59D - GGsGG - Fc Knob (C220S, K447A, N297D, L234A, L235A, M252Y, S254T, T256E):
TTTAAAGCCGCCACCATGGAGACAGACACACTCCTGCTATGGGTACTGCTGCTCTGGGTT
CCAGGTGAGAGCTGCAGCCTGACTGCATaGGGGCTGGGATaGGCATAAGAATAAAGGT
CTGTGTGGACAGCCTTCTGaTTCAGCCACGACCTCTGTGTATcCTTCTcACCCCAcagGTTCC
ACCGGTCTCCCGATATGTCCGGGCGGGGCCGCTCGGTGCCAGGTAACTTTGAGGGAC
CTGTTTGACCGAGCCGTAGTCCTTTCACACTATATTCACGACCTCTCATCTGAGATGTTTTCC
GAGTTCGACAAGAGATATACCCACGGTCGCGGGTTTATAACTAAGGCAATAAACAGTTGC
CATACCTCAAGTCTCGCTACACCCGAGGACAAGGAACAAGCGCAACAGATGAATCAGAA
GGACTTTTTGTCACTGATAGTGTCCATCCTGCGCAGTTGGAACGAACCCTTGTACCATTTGG
TCACCGAAGTCAGGGGGATGCAAGAAGCACCGGAGGCTATACTGTCAAAGGCCGTAG
AAATCGAAGAACAGACGAAGAGACTCCTGGAAGGTATGGAACTCATAGTGTCCCAGGTC
CACCCAGAGACAAAAGAGAACGAAATATACCCCGTATGGTCTGGCTTGCCTTCCCTGCAA
ATGGCAGATGAAGAGAGTCGGTTGAGTGCCTATTACAACCTTCTCCACTGTCTCAGGAGG
GACAGTCACAAGATCGATAACTATCTCAAACTCCTTAAGTGTAGGATAATTCATAACAATAAC
TGTGGCGGTAGCGGTGGCgagcccaaatctagcgacaaaactcacacatgcccaccgtgcccaggtaagccag
cccaggcctcgccctccagctcaaggcgggacaggtgccctagagtagcctgcatccagggacaggccccagccgggtgctg
acacgtccacctccatctcttcctcagcacctgaaGCCGCTgggggaccgtcagtcttcctcttccccccaaaacccaaggaca
ccctcTATatcACCcggGAAcctgaggtcacatgcgtggtggtggacgtgagccacgaagaccctgaggtcaagttcaactg
gtacgtggacggcgtggaggtgcataatgccaagacaaagccgagggaggagcagtacgacagcacgtaccgtgtggtcag
cgtcctcaccgtcctgcaccaggactggctgaatggcaaggagtacaagtgcaaggtctccaacaaagccctcccagcccccat
cgagaaaaccatctccaaagccaaaggtgggacccgtggggtgcgagggccacatggacagaggccagctcagcccacc
ctctgccctgagagtgaccgctgtaccaacctctgtccctacagggcagccccgagaaccacaggtgtacaccctgcccccatcc
agggatgagctgaccaagaaccaggtcagcctgtggtgcctggtcaaaggcttctatcccagcgacatcgccgtggagtgggag
agcaatgggcagccggagaacaactacaagaccacgcctcccgtgctggactccgacggctccttcttcctctacagcaagctca
ccgtggacaagagcaggtggcagcaggggaacgtcttctcatgctccgtgatgcatgaggctctgcacaaccactacacacaga
agagcctctccctgtctcccggggcaTGA

Fc Hole (C220S, N297D, K447A, T366S, L368A, Y407V, H435R, Y436F, L234A, L235A, M252Y, S254T, T256E):
TTTAAAGCCGCCACCATGGAGACAGACACACTCCTGCTATGGGTACTGCTGCTCTGGGTT
CCAGGTGAGAGCTGCAGCCTGACTGCATaGGGGCTGGGATaGGCATAAGAATAAAGGT
CTGTGTGGACAGCCTTCTGaTTCAGCCACGACCTCTGTGTATcCTTCTcACCCCAcagGTTCC
ACCGGTgagcccaaatctagcgacaaaactcacacatgcccaccgtgcccaggtaagccagcccaggcctcgccctcca
gctcaaggcgggacaggtgccctagagtagcctgcatccagggacaggccccagccgggtgctgacacgtccacctccatctct
tcctcagcacctgaaGCCGCTgggggaccgtcagtcttcctcttccccccaaaacccaaggacaccctcTATatcACCcgg
GAAcctgaggtcacatgcgtggtggtggacgtgagccacgaagaccctgaggtcaagttcaactggtacgtggacggcgtgg
aggtgcataatgccaagacaaagccgagggaggagcagtacgacagcacgtaccgtgtggtcagcgtcctcaccgtcctgca
ccaggactggctgaatggcaaggagtacaagtgcaaggtctccaacaaagccctcccagcccccatcgagaaaaccatctcc
aaagccaaaggtgggacccgtggggtgcgagggccacatggacagaggccagctcagcccaccctctgccctgagagtgac
cgctgtaccaacctctgtccctacagggcagccccgagaaccacaggtgtacaccctgcccccatccagggatgagctgaccaa
gaaccaggtcagcctgagctgcgccgtcaaaggcttctatcccagcgacatcgccgtggagtgggagagcaatgggcagccg
gagaacaactacaagaccacgcctcccgtgctggactccgacggctccttcttcctcgtcagcaagctcaccgtggacaagagca
ggtggcagcaggggaacgtcttctcatgctccgtgatgcatgaggctctgcacaacCGTTTTacacagaagagcctctccctgtc
tcccggggcaTGA

25. PRL (N59D) - GGsGG - Fc A (C220S, K447A, N297D, T350V, L351Y, F405A, Y407V):
TTTAAAGCCGCCACCATGGAGACAGACACACTCCTGCTATGGGTACTGCTGCTCTGGGTT
CCAGGTGAGAGCTGCAGCCTGACTGCATaGGGGCTGGGATaGGCATAAGAATAAAGGT
CTGTGTGGACAGCCTTCTGaTTCAGCCACGACCTCTGTGTATcCTTCTcACCCCAcagGTTCC
ACCGGTCTCCCGATATGTCCGGGCGGGGCCGCTCGGTGCCAGGTAACTTTGAGGGAC

| | |
|---|---|
| | CTGTTTGACCGAGCCGTAGTCCTTTCACACTATATTCACGACCTCTCATCTGAGATGTTTTCC<br>GAGTTCGACAAGAGATATACCCACGGTCGCGGGTTTATAACTAAGGCAATAAACAGTTGC<br>CATACCTCAAGTCTCGCTACACCCGAGGACAAGGAACAAGCGCAACAGATGAATCAGAA<br>GGACTTTTTGTCACTGATAGTGTCCATCCTGCGCAGTTGGAACGAACCCTTGTACCATTTGG<br>TCACCGAAGTCAGGGGGATGCAAGAAGCACCGGAGGCTATACTGTCAAAGGCCGTAG<br>AAATCGAAGAACAGACGAAGAGACTCCTGGAAGGTATGGAACTCATAGTGTCCCAGGTC<br>CACCCAGAGACAAAAGAGAACGAAATATACCCCGTATGGTCTGGCTTGCCTTCCCTGCAA<br>ATGGCAGATGAAGAGAGTCGGTTGAGTGCCTATTACAACCTTCTCCACTGTCTCAGGAGG<br>GACAGTCACAAGATCGATAACTATCTCAAACTCCTTAAGTGTAGGATAATTCATAACAATAAC<br>TGTGGCGGTAGCGGTGGCgagcccaaatctagcgacaaaactcacacatgcccaccgtgcccaggtaagccag<br>cccaggcctcgccctccagctcaaggcgggacaggtgccctagagtagcctgcatccagggacaggccccagccgggtgctg<br>acacgtccacctccatctcttcctcagcacctgaactcctggggggaccgtcagtcttcctcttccccccaaaacccaaggacaccctc<br>atgatctcccggacccctgaggtcacatgcgtggtggtggacgtgagccacgaagaccctgaggtcaagttcaactggtacgtgg<br>acggcgtggaggtgcataatgccaagacaaagccgagggaggagcagtacgacagcacgtaccgtgtggtcagcgtcctca<br>ccgtcctgcaccaggactggctgaatggcaaggagtacaagtgcaaggtctccaacaaagccctcccagcccccatcgagaa<br>aaccatctccaaagccaaaggtgggacccgtggggtgcgagggccacatggacagaggccagctcagcccaccctctgccct<br>gagagtgaccgctgtaccaacctctgtccctacagggcagccccgagaaccacaggtgtacGTGTATccccatccagggat<br>gagctgaccaagaaccaggtcagcctgCTGtgcctggtcaaaggcttctatcccagcgacatcgccgtggagtgggagagca<br>atgggcagccggagaacaactacaagaccacgcctcccgtgctggactccgacggctccttcGCGctcGTGagcaagctc<br>accgtggacaagagcaggtggcagcaggggaacgtcttctcatgctccgtgatgcatgaggctctgcacaaccactacacacag<br>aagagcctctccctgtctcccggggcaTGA<br><br>Fc B (C220S, N297D, K427A, T350V, T366L, K293L, T394W):<br>TTTAAAGCCGCCACCATGGAGACAGACACACTCCTGCTATGGGTACTGCTGCTCTGGGTT<br>CCAGGTGAGAGCTGCAGCCTGACTGCATaGGGGCTGGGATaGGCATAAGAATAAAGGT<br>CTGTGTGGACAGCCTTCTGaTTCAGCCACGACCTCTGTGTATcCTTCTcACCCCAcagGTTCC<br>ACCGGTgagcccaaatctagcgacaaaactcacacatgcccaccgtgcccaggtaagccagcccaggcctcgccctcca<br>gctcaaggcgggacaggtgccctagagtagcctgcatccagggacaggccccagccgggtgctgacacgtccacctccatctct<br>tcctcagcacctgaactcctggggggaccgtcagtcttcctcttccccccaaaacccaaggacaccctcatgatctcccggacccctg<br>aggtcacatgcgtggtggtggacgtgagccacgaagaccctgaggtcaagttcaactggtacgtggacggcgtggaggtgcata<br>atgccaagacaaagccgagggaggagcagtacgacagcacgtaccgtgtggtcagcgtcctcaccgtcctgcaccaggactg<br>gctgaatggcaaggagtacaagtgcaaggtctccaacaaagccctcccagcccccatcgagaaaaccatctccaaagccaaa<br>ggtgggacccgtggggtgcgagggccacatggacagaggccagctcagcccaccctctgccctgagagtgaccgctgtacca<br>acctctgtccctacagggcagccccgagaaccacaggtgtacGTGctgccccatccagggatgagctgaccaagaaccag<br>gtcagcctgCTGtgcctggtcaaaggcttctatcccagcgacatcgccgtggagtgggagagcaatgggcagccggagaaca<br>actacCTGaccTGGcctcccgtgctggactccgacggctccttcttcctctacagcaagctcaccgtggacaagagcaggtggc<br>agcaggggaacgtcttctcatgctccgtgatgcatgaggctctgcacaacCGTTTTacacagaagagcctctccctgtctcccgg<br>ggcaTGA |
| 26. | PRL (N59D) - GGsGG- Fc A (C220S, K447A, N297D, T350V, L351Y, F405A, Y407V,<br>L234A, L235A, V264E, L309D, Q311H, N434S):<br>TTTAAAGCCGCCACCATGGAGACAGACACACTCCTGCTATGGGTACTGCTGCTCTGGGTT<br>CCAGGTGAGAGCTGCAGCCTGACTGCATaGGGGCTGGGATaGGCATAAGAATAAAGGT<br>CTGTGTGGACAGCCTTCTGaTTCAGCCACGACCTCTGTGTATcCTTCTcACCCCAcagGTTCC<br>ACCGGTTTGCCCATCTGTCCCGGCGGGGCTGCCCGATGCCAGGTGACCCTTCGAGAC<br>CTGTTTGACCGCGCCGTCGTCCTGTCCCACTACATCCATGACCTCTCCTCAGAAATGTTCA<br>GCGAATTCGATAAACGGTATACCCATGGCCGGGGGTTCATTACCAAGGCCATCAACAGC<br>TGCCACACTTCTTCCCTTGCCACCCCCGAAGACAAGGAGCAAGCCCAACAGATGAATCA<br>AAAAGACTTTCTGAGCCTGATAGTCAGCATATTGCGATCCTGGAATGAGCCTCTGTATCATC<br>TGGTCACGGAAGTACGTGGTATGCAAGAAGCCCCGGAGGCTATCCTATCCAAAGCTGTA |

GAGATTGAGGAGCAAACCAAACGGCTTCTAGAGGGCATGGAGCTGATAGTCAGCCAGG
TTCATCCTGAAACCAAAGAAAATGAGATCTACCCTGTCTGGTCGGGACTTCCATCCCTGCA
GATGGCTGATGAAGAGTCTCGCCTTTCTGCTTATTATAACCTGCTCCACTGCCTACGCAGG
GATTCACATAAAATCGACAATTATCTCAAGCTCCTGAAGTGCCGAATCATCCACAACAACAA
CTGCGGCGGTAGCGGTGGCgagcccaaatctagcgacaaaactcacacatgcccaccgtgcccaggtaagcc
agcccaggcctcgccctccagctcaaggcgggacaggtgccctagagtagcctgcatccagggacaggccccagccgggtg
ctgacacgtccacctccatctcttcctcagcacctgaaGCGGCGgggggaccgtcagtcttcctcttccccccaaaacccaagg
acaccctcatgatctcccggacccctgaggtcacatgcgtggtgGAAgacgtgagccacgaagaccctgaggtcaagttcaact
ggtacgtggacggcgtggaggtgcataatgccaagacaaagccgagggaggagcagtacgacagcacgtaccgtgtggtca
gcgtcctcaccgtcGATcacCATgactggctgaatggcaaggagtacaagtgcaaggtctccaacaaagccctcccagcccc
catcgagaaaaccatctccaaagccaaaggtgggacccgtggggtgcgagggccacatggacagaggccagctcagccca
ccctctgccctgagagtgaccgctgtaccaacctctgtccctacagggcagccccgagaaccacaggtgtacGTGTATccccc
atccagggatgagctgaccaagaaccaggtcagcctgCTGtgcctggtcaaaggcttctatcccagcgacatcgccgtggagt
gggagagcaatgggcagccggagaacaactacaagaccacgcctcccgtgctggactccgacggctccttcGCGctcGT
GagcaagctcaccgtggacaagagcaggtggcagcaggggaacgtcttctcatgctccgtgatgcatgaggctctgcacAGC
cactacacacagaagagcctctccctgtctcccggggcaTGA

Fc B (N297D, C220S, K447A, L234A, L235A, V264E, L309D, Q311H, N434S):
TTTAAAGCCGCCACCATGGAGACAGACACACTCCTGCTATGGGTACTGCTGCTCTGGGTT
CCAGGTGAGAGCTGCAGCCTGACTGCATaGGGGCTGGGATaGGCATAAGAATAAAGGT
CTGTGTGGACAGCCTTCTGaTTCAGCCACGACCTCTGTGTATcCTTCTcACCCCAcagGTTCC
ACCGGTgagcccaaatctagcgacaaaactcacacatgcccaccgtgcccaggtaagccagcccaggcctcgccctcca
gctcaaggcgggacaggtgccctagagtagcctgcatccagggacaggccccagccgggtgctgacacgtccacctccatctct
tcctcagcacctgaaGCGGCGgggggaccgtcagtcttcctcttccccccaaaacccaaggacaccctcatgatctcccgga
cccctgaggtcacatgcgtggtgGAAgacgtgagccacgaagaccctgaggtcaagttcaactggtacgtggacggcgtgga
ggtgcataatgccaagacaaagccgagggaggagcagtacgacagcacgtaccgtgtggtcagcgtcctcaccgtcGATca
cCATgactggctgaatggcaaggagtacaagtgcaaggtctccaacaaagccctcccagcccccatcgagaaaaccatctcc
aaagccaaaggtgggacccgtggggtgcgagggccacatggacagaggccagctcagcccaccctctgccctgagagtgac
cgctgtaccaacctctgtccctacagggcagccccgagaaccacaggtgtacGTGctgcccccatccagggatgagctgacca
agaaccaggtcagcctgCTGtgcctggtcaaaggcttctatcccagcgacatcgccgtggagtgggagagcaatgggcagcc
ggagaacaactacCTGaccTGGcctcccgtgctggactccgacggctccttcttcctctacagcaagctcaccgtggacaaga
gcaggtggcagcaggggaacgtcttctcatgctccgtgatgcatgaggctctgcacAGCcactacacacagaagagcctctccc
tgtctcccggggcaTGA

27. PRL (N59D) - GGsGG- Fc A (C220S, K447A, N297D, T350V, L351Y, F405A, Y407V, L234A, L235A, V264E, L309D, Q311H, N434S):
TTTAAAGCCGCCACCATGGAGACAGACACACTCCTGCTATGGGTACTGCTGCTCTGGGTT
CCAGGTGAGAGCTGCAGCCTGACTGCATaGGGGCTGGGATaGGCATAAGAATAAAGGT
CTGTGTGGACAGCCTTCTGaTTCAGCCACGACCTCTGTGTATcCTTCTcACCCCAcagGTTCC
ACCGGTTTGCCCATCTGTCCCGGCGGGGCTGCCCGATGCCAGGTGACCCTTCGAGAC
CTGTTTGACCGCGCCGTCGTCCTGTCCCACTACATCCATGACCTCTCCTCAGAAATGTTCA
GCGAATTCGATAAACGGTATACCCATGGCCGGGGGTTCATTACCAAGGCCATCAACAGC
TGCCACACTTCTTCCCTTGCCACCCCCGAAGACAAGGAGCAAGCCCAACAGATGAATCA
AAAAGACTTTCTGAGCCTGATAGTCAGCATATTGCGATCCTGGAATGAGCCTCTGTATCATC
TGGTCACGGAAGTACGTGGTATGCAAGAAGCCCCGGAGGCTATCCTATCCAAAGCTGTA
GAGATTGAGGAGCAAACCAAACGGCTTCTAGAGGGCATGGAGCTGATAGTCAGCCAGG
TTCATCCTGAAACCAAAGAAAATGAGATCTACCCTGTCTGGTCGGGACTTCCATCCCTGCA
GATGGCTGATGAAGAGTCTCGCCTTTCTGCTTATTATAACCTGCTCCACTGCCTACGCAGG
GATTCACATAAAATCGACAATTATCTCAAGCTCCTGAAGTGCCGAATCATCCACAACAACAA
CTGCGGCGGTAGCGGTGGCgagcccaaatctagcgacaaaactcacacatgcccaccgtgcccaggtaagcc

| | |
|---|---|
| | agcccaggcctcgccctccagctcaaggcgggacaggtgccctagagtagcctgcatccagggacaggccccagccgggtg<br>ctgacacgtccacctccatctcttcctcagcacctgaaGCGGCGgggggaccgtcagtcttcctcttccccccaaaacccaagg<br>acaccctcatgatctcccggacccctgaggtcacatgcgtggtgGAAgacgtgagccacgaagaccctgaggtcaagttcaact<br>ggtacgtggacggcgtggaggtgcataatgccaagacaaagccgagggaggagcagtacgacagcacgtaccgtgtggtca<br>gcgtcctcaccgtcGATcacCATgactggctgaatggcaaggagtacaagtgcaaggtctccaacaaagccctcccagcccc<br>catcgagaaaaccatctccaaagccaaaggtgggacccgtggggtgcgagggccacatggacagaggccagctcagccca<br>ccctctgccctgagagtgaccgctgtaccaacctctgtccctacagggcagccccgagaaccacaggtgtacGTGTATccccc<br>atccagggatgagctgaccaagaaccaggtcagcctgCTGtgcctggtcaaaggcttctatcccagcgacatcgccgtggagt<br>gggagagcaatgggcagccggagaacaactacaagaccacgcctcccgtgctggactccgacggctccttcGCGctcGT<br>GagcaagctcaccgtggacaagagcaggtggcagcaggggaacgtcttctcatgctccgtgatgcatgaggctctgcacAGC<br>cactacacacagaagagcctctccctgtctcccggggcaTGA<br><br>Fc B (N297D, C220S, K447A, L234A, L235A, V264E, L309D, Q311H, N434S, H435R, Y436F):<br>TTTAAAGCCGCCACCATGGAGACAGACACACTCCTGCTATGGGTACTGCTGCTCTGGGTT<br>CCAGGTGAGAGCTGCAGCCTGACTGCATaGGGGCTGGGATaGGCATAAGAATAAAGGT<br>CTGTGTGGACAGCCTTCTGaTTCAGCCACGACCTCTGTGTATcCTTCTcACCCCAcagGTTCC<br>ACCGGTgagcccaaatctagcgacaaaactcacacatgcccaccgtgcccaggtaagccagcccaggcctcgccctcca<br>gctcaaggcgggacaggtgccctagagtagcctgcatccagggacaggccccagccgggtgctgacacgtccacctccatctct<br>tcctcagcacctgaaGCGGCGgggggaccgtcagtcttcctcttccccccaaaacccaaggacaccctcatgatctcccgga<br>cccctgaggtcacatgcgtggtgGAAgacgtgagccacgaagaccctgaggtcaagttcaactggtacgtggacggcgtgga<br>ggtgcataatgccaagacaaagccgagggaggagcagtacgacagcacgtaccgtgtggtcagcgtcctcaccgtcGATca<br>cCATgactggctgaatggcaaggagtacaagtgcaaggtctccaacaaagccctcccagcccccatcgagaaaaccatctcc<br>aaagccaaaggtgggacccgtggggtgcgagggccacatggacagaggccagctcagcccaccctctgccctgagagtgac<br>cgctgtaccaacctctgtccctacagggcagccccgagaaccacaggtgtacGTGctgcccccatccagggatgagctgacca<br>agaaccaggtcagcctgCTGtgcctggtcaaaggcttctatcccagcgacatcgccgtggagtgggagagcaatgggcagcc<br>ggagaacaactacCTGaccTGGcctcccgtgctggactccgacggctccttcttcctctacagcaagctcaccgtggacaaga<br>gcaggtggcagcaggggaacgtcttctcatgctccgtgatgcatgaggctctgcacAGCCGTTTTacacagaagagcctctc<br>cctgtctcccggggcaTGA |
| 28. | PRL N59D - GGsGG- Fc A (C220S, K447A, N297D, T350V, L351Y, F405A, Y407V, L234A, L235A, P329G, V264E, L309D, Q311H, N434S):<br>TTTAAAGCCGCCACCATGGAGACAGACACACTCCTGCTATGGGTACTGCTGCTCTGGGTT<br>CCAGGTGAGAGCTGCAGCCTGACTGCATaGGGGCTGGGATaGGCATAAGAATAAAGGT<br>CTGTGTGGACAGCCTTCTGaTTCAGCCACGACCTCTGTGTATcCTTCTcACCCCAcagGTTCC<br>ACCGGTTTGCCCATCTGTCCCGGCGGGGCTGCCCGATGCCAGGTGACCCTTCGAGAC<br>CTGTTTGACCGCGCCGTCGTCCTGTCCCACTACATCCATGACCTCTCCTCAGAAATGTTCA<br>GCGAATTCGATAAACGGTATACCCATGGCCGGGGGTTCATTACCAAGGCCATCAACAGC<br>TGCCACACTTCTTCCCTTGCCACCCCCGAAGACAAGGAGCAAGCCCAACAGATGAATCA<br>AAAAGACTTTCTGAGCCTGATAGTCAGCATATTGCGATCCTGGAATGAGCCTCTGTATCATC<br>TGGTCACGGAAGTACGTGGTATGCAAGAAGCCCCGGAGGCTATCCTATCCAAAGCTGTA<br>GAGATTGAGGAGCAAACCAAACGGCTTCTAGAGGGCATGGAGCTGATAGTCAGCCAGG<br>TTCATCCTGAAACCAAAGAAAATGAGATCTACCCTGTCTGGTCGGGACTTCCATCCCTGCA<br>GATGGCTGATGAAGAGTCTCGCCTTTCTGCTTATTATAACCTGCTCCACTGCCTACGCAGG<br>GATTCACATAAAATCGACAATTATCTCAAGCTCCTGAAGTGCCGAATCATCCACAACAACAA<br>CTGCGGCGGTAGCGGTGGCgagcccaaatctagcgacaaaactcacacatgcccaccgtgcccaggtaagcc<br>agcccaggcctcgccctccagctcaaggcgggacaggtgccctagagtagcctgcatccagggacaggccccagccgggtg<br>ctgacacgtccacctccatctcttcctcagcacctgaaGCGGCGgggggaccgtcagtcttcctcttccccccaaaacccaagg<br>acaccctcatgatctcccggacccctgaggtcacatgcgtggtgGAAgacgtgagccacgaagaccctgaggtcaagttcaact<br>ggtacgtggacggcgtggaggtgcataatgccaagacaaagccgagggaggagcagtacgacagcacgtaccgtgtggtca |

| | |
|---|---|
| | gcgtcctcaccgtcGATcacCATgactggctgaatggcaaggagtacaagtgcaaggtctccaacaaagccctcGGAgcc<br>cccatcgagaaaaccatctccaaagccaaaggtgggacccgtggggtgcgagggccacatggacagaggccagctcagcc<br>caccctctgccctgagagtgaccgctgtaccaacctctgtccctacagggcagccccgagaaccacaggtgtacGTGTATccc<br>ccatccagggatgagctgaccaagaaccaggtcagcctgCTGtgcctggtcaaaggcttctatcccagcgacatcgccgtgga<br>gtgggagagcaatgggcagccggagaacaactacaagaccacgcctcccgtgctggactccgacggctccttcGCGctcG<br>TGagcaagctcaccgtggacaagagcaggtggcagcaggggaacgtcttctcatgctccgtgatgcatgaggctctgcacAG<br>CcactacacacagaagagcctctccctgtctcccggggcaTGA<br><br>Fc B (N297D, C220S, K447A, L234A, L235A, P329G, V264E, L309D, Q311H, N434S, H435R, Y436F):<br>TTTAAAGCCGCCACCATGGAGACAGACACACTCCTGCTATGGGTACTGCTGCTCTGGGTT<br>CCAGGTGAGAGCTGCAGCCTGACTGCATaGGGGCTGGGATaGGCATAAGAATAAAGGT<br>CTGTGTGGACAGCCTTCTGaTTCAGCCACGACCTCTGTGTATcCTTCTcACCCCAcagGTTCC<br>ACCGGTgagcccaaatctagcgacaaaactcacacatgcccaccgtgcccaggtaagccagcccaggcctcgccctcca<br>gctcaaggcgggacaggtgccctagagtagcctgcatccagggacaggccccagccgggtgctgacacgtccacctccatctct<br>tcctcagcacctgaaGCGGCGgggggaccgtcagtcttcctcttccccccaaaacccaaggacaccctcatgatctcccgga<br>cccctgaggtcacatgcgtggtgGAAgacgtgagccacgaagaccctgaggtcaagttcaactggtacgtggacggcgtgga<br>ggtgcataatgccaagacaaagccgagggaggagcagtacgacagcacgtaccgtgtggtcagcgtcctcaccgtcGATca<br>cCATgactggctgaatggcaaggagtacaagtgcaaggtctccaacaaagccctcGGAgcccccatcgagaaaaccatct<br>ccaaagccaaaggtgggacccgtggggtgcgagggccacatggacagaggccagctcagcccaccctctgccctgagagtg<br>accgctgtaccaacctctgtccctacagggcagccccgagaaccacaggtgtacGTGctgcccccatccagggatgagctgac<br>caagaaccaggtcagcctgCTGtgcctggtcaaaggcttctatcccagcgacatcgccgtggagtgggagagcaatgggcag<br>ccggagaacaactacCTGaccTGGcctcccgtgctggactccgacggctccttcttcctctacagcaagctcaccgtggacaa<br>gagcaggtggcagcaggggaacgtcttctcatgctccgtgatgcatgaggctctgcacAGCCGTTTTacacagaagagcct<br>ctccctgtctcccggggcaTGA |
| 29. | PRL N59D - GGsGG - Fc A (C220S, K447A, N297D, T350V, L351Y, F405A, Y407V, L234A, L235A, M252Y, S254T, T256E):<br>TTTAAAGCCGCCACCATGGAGACAGACACACTCCTGCTATGGGTACTGCTGCTCTGGGTT<br>CCAGGTGAGAGCTGCAGCCTGACTGCATaGGGGCTGGGATaGGCATAAGAATAAAGGT<br>CTGTGTGGACAGCCTTCTGaTTCAGCCACGACCTCTGTGTATcCTTCTcACCCCAcagGTTCC<br>ACCGGTCTCCCGATATGTCCGGGCGGGGCCGCTCGGTGCCAGGTAACTTTGAGGGAC<br>CTGTTTGACCGAGCCGTAGTCCTTTCACACTATATTCACGACCTCTCATCTGAGATGTTTTCC<br>GAGTTCGACAAGAGATATACCCACGGTCGCGGGTTTATAACTAAGGCAATAAACAGTTGC<br>CATACCTCAAGTCTCGCTACACCCGAGGACAAGGAACAAGCGCAACAGATGAATCAGAA<br>GGACTTTTTGTCACTGATAGTGTCCATCCTGCGCAGTTGGAACGAACCCTTGTACCATTTGG<br>TCACCGAAGTCAGGGGGATGCAAGAAGCACCGGAGGCTATACTGTCAAAGGCCGTAG<br>AAATCGAAGAACAGACGAAGAGACTCCTGGAAGGTATGGAACTCATAGTGTCCCAGGTC<br>CACCCAGAGACAAAAGAGAACGAAATATACCCCGTATGGTCTGGCTTGCCTTCCCTGCAA<br>ATGGCAGATGAAGAGAGTCGGTTGAGTGCCTATTACAACCTTCTCCACTGTCTCAGGAGG<br>GACAGTCACAAGATCGATAACTATCTCAAACTCCTTAAGTGTAGGATAATTCATAACAATAAC<br>TGTGGCGGTAGCGGTGGCgagcccaaatctagcgacaaaactcacacatgcccaccgtgcccaggtaagccag<br>cccaggcctcgccctccagctcaaggcgggacaggtgccctagagtagcctgcatccagggacaggccccagccgggtgctg<br>acacgtccacctccatctcttcctcagcacctgaaGCCGCTgggggaccgtcagtcttcctcttccccccaaaacccaaggaca<br>ccctcTATatcACCcggGAAcctgaggtcacatgcgtggtggtggacgtgagccacgaagaccctgaggtcaagttcaactg<br>gtacgtggacggcgtggaggtgcataatgccaagacaaagccgagggaggagcagtacgacagcacgtaccgtgtggtcag<br>cgtcctcaccgtcctgcaccaggactggctgaatggcaaggagtacaagtgcaaggtctccaacaaagccctcccagcccccat<br>cgagaaaaccatctccaaagccaaaggtgggacccgtggggtgcgagggccacatggacagaggccagctcagcccacc<br>ctctgccctgagagtgaccgctgtaccaacctctgtccctacagggcagccccgagaaccacaggtgtacGTGTATcccccatc<br>cagggatgagctgaccaagaaccaggtcagcctgCTGtgcctggtcaaaggcttctatcccagcgacatcgccgtggagtggg<br>agagcaatgggcagccggagaacaactacaagaccacgcctcccgtgctggactccgacggctccttcGCGctcGTGagc |

| | |
|---|---|
| | aagctcaccgtggacaagagcaggtggcagcaggggaacgtcttctcatgctccgtgatgcatgaggctctgcacaaccactac<br>acacagaagagcctctccctgtctcccggggcaTGA<br><br>Fc B (N297D, K447A, H435R, Y436F, L234A, L235A, M252Y, S254T, T256E):<br>TTTAAAGCCGCCACCATGGAGACAGACACACTCCTGCTATGGGTACTGCTGCTCTGGGTT<br>CCAGGTGAGAGCTGCAGCCTGACTGCATaGGGGCTGGGATaGGCATAAGAATAAAGGT<br>CTGTGTGGACAGCCTTCTGaTTCAGCCACGACCTCTGTGTATcCTTCTcACCCCAcagGTTCC<br>ACCGGTgagcccaaatctagcgacaaaactcacacatgcccaccgtgcccaggtaagccagcccaggcctcgccctcca<br>gctcaaggcgggacaggtgccctagagtagcctgcatccagggacaggccccagccgggtgctgacacgtccacctccatctct<br>tcctcagcacctgaaGCCGCTgggggaccgtcagtcttcctcttccccccaaaacccaaggacaccctcTATatcACCcgg<br>GAAcctgaggtcacatgcgtggtggtggacgtgagccacgaagaccctgaggtcaagttcaactggtacgtggacggcgtgg<br>aggtgcataatgccaagacaaagccgagggaggagcagtacgacagcacgtaccgtgtggtcagcgtcctcaccgtcctgca<br>ccaggactggctgaatggcaaggagtacaagtgcaaggtctccaacaaagccctcccagcccccatcgagaaaaccatctcc<br>aaagccaaaggtgggacccgtggggtgcgagggccacatggacagaggccagctcagcccaccctctgccctgagagtgac<br>cgctgtaccaacctctgtccctacagggcagccccgagaaccacaggtgtacGTGctgcccccatccagggatgagctgacca<br>agaaccaggtcagcctgCTGtgcctggtcaaaggcttctatcccagcgacatcgccgtggagtgggagagcaatgggcagcc<br>ggagaacaactacCTGaccTGGcctcccgtgctggactccgacggctccttcttcctctacagcaagctcaccgtggacaaga<br>gcaggtggcagcaggggaacgtcttctcatgctccgtgatgcatgaggctctgcacaacCGTTTTacacagaagagcctctccc<br>tgtctcccggggcaTGA |

Constructs for pichia expression of Fc-PRL-13:
Fc A:
Signal Sequence
Fc-PRL-13 Fc A sequence

ATGAGGTTCCCAAGTATATTTACAGCCGTACTTTTTGCAGCATCATCAGCCCTGGCTGCTCCAGTGAATA
CAACTACAGAAGACGAGACCGCTCAGATACCAGCCGAGGCAGTCATCGGCTACTCAGACCTGGAA
GGAGACTTTGACGTCGCTGTCCTGCCCTTCAGTAACTCCACGAACAATGGATTATTGTTCATCAATACGA
CCATTGCATCCATCGCCGCTAAAGAGGAGGGCGTGAGTTTGGAGAAGAGAGAGGAACCCAAGTCAT
CCGATAAGACTCATACATGTCCCCCATGTCCCGCACCTGAGCTTCTAGGTGGACCTTCTGTTTTTCTTTTTCC
TCCTAAACCAAAGGACACCCTTATGATATCACGTACGCCTGAAGTGACCTGTGTAGTCGTGGATGTCAGTC
ACGAAGACCCCGAAGTTAAATTTAATTGGTATGTAGATGGTGTGGAAGTACATAACGCAAAGACTAAGCCC
AGAGAGGAGCAGTACGATTCAACTTATCGTGTGGTATCTGTGTTGACCGTCCTGCATCAGGACTGGCTTAA
CGGAAAAGAATACAAGTGCAAAGTCTCTAATAAAGCCCTGCCTGCACCCATCGAAAAAACAATCTCAAAA
GCTAAGGGCCAACCTAGGGAACCCCAGGTCTACGTCTACCCCCCATCTAGAGACGAACTGACTAAGAA
TCAGGTAAGTCTGACTTGTCTTGTAAAAGGCTTCTACCCATCCGATATCGCCGTAGAATGGGAGTCCAACG
GTCAGCCTGAAAACAATTACAAAACGACTCCTCCCGTGCTTGACTCCGACGGATCATTTGCACTGGTCTCT
AAATTGACCGTAGATAAGTCCAGGTGGCAGCAGGGCAATGTTTTCAGTTGCAGTGTTATGCACGAGGCC
CTACACAACCATTACACCCAGAAGTCACTTTCCCTTTCCCCTGGAGCAGGAGGATCCGGTGGCCTACCA
ATATGCCCTGGAGGCGCAGCCAGATGTCAAGTCACCTTACGTGACTTATTCGACCGTGCTGTTGTCCTGT
CACATTACATACATGACTTATCTTCAGAAATGTTCAGTGAGTTCGACAAGAGGTACACTCATGGCAGAGGTT
TTATAACCAAAGCAATCAATAGTTGTCACACGTCCTCATTGGCTACCCCCGAAGACAAGGAACAGGCACA
ACAGATGAATCAAAAAGATTTTCTGTCTTTGATCGTCTCCATCCTTAGATCTTGGAACGAGCCCCTTTACCATC
TGGTAACTGAAGTTAGGGGTATGCAAGAGGCACCCGAGGCAATTTTATCCAAGGCTGTGGAGATAGAAG
AACAGACGAAGCGTCTGCTAGAGGGAATGGAGCTTATAGTTTCCCAGGTTCATCCAGAAACCAAAGAGA
ACGAGATTTATCCTGTGTGGTCAGGCCTACCCTCACTGCAAATGGCCGACGAAGAATCTCGTCTATCCGC
CTACTACAATTTGTTACACTGTCTGCGTAGGGATTCACACAAAATCGACAACTATTTGAAATTGCTTAAATGTA
GAATAATCCATAACAATAACTGCTGA

Fc B:

Signal Sequence

Fc-PRL-13 Fc B sequence

ATGAGATTCCCCAGTATATTCACGGCCGTACTTTTCGCAGCATCTTCAGCCCTAGCCGCCCCAGTTAAC
ACTACGACCGAAGATGAAACAGCTCAAATCCCCGCAGAGGCAGTCATCGGCTATTCTGATTTGGAAG
GCGACTTCGATGTCGCCGTCCTGCCATTTAGTAATTCCACGAACAACGGTCTTCTGTTCATTAACACGA
CGATTGCTTCCATCGCAGCAAAAGAAGAAGGTGTTTCCCTGGAAAAGAGAGAGGAGCCAAAATCTTCT
GATAAGACTCATACCTGCCCACCATGCCCTGCTCCCGAGTTATTAGGCGGCCCTTCTGTGTTCTTATTCCC
CCCCAAACCTAAAGATACGTTGATGATAAGTAGGACGCCTGAAGTCACTTGCGTTGTCGTGGATGTATCC
CACGAGGACCCTGAGGTGAAGTTTAACTGGTACGTGGACGGTGTGGAGGTTCATAATGCAAAAACAAAG
CCTCGTGAGGAACAATATGACTCTACCTATCGTGTCGTATCAGTTTTAACAGTCCTTCACCAAGATTGGCTAA
ATGGTAAGGAATACAAATGCAAGGTCTCAAACAAGGCACTTCCTGCACCAATCGAGAAAACTATAAGTAA
GGCAAAGGGACAACCTAGGGAGCCACAGGTTTACGTGCTTCCTCCCAGTAGAGATGAATTAACAAAGAA
TCAAGTATCACTTCTGTGCTTGGTAAAGGGATTCTACCCATCTGATATAGCCGTGGAGTGGGAGTCCAACG
GACAGCCCGAGAATAATTATCTTACATGGCCTCCAGTGCTGGATTCCGATGGCTCCTTTTTCCTTTACTCAAA
ACTGACTGTGGACAAAAGTAGATGGCAGCAAGGTAATGTTTTCAGTTGTTCTGTCATGCACGAGGCATTGC
ATAACCGTTTTACCCAAAAGAGTTTATCCCTAAGTCCCGGTGCATGA

Table S4: Bonferroni-adjusted p-values for multiple comparison via Two-way ANOVA

| | Pup weight compared to Vehicle + PBS control | |
|---|---|---|
| Day | BR + PBS | BR + 5mg/kg Fc-PRL-13 (single dose) |
| 7 | 0.0305 (*) | 0.0726 (ns) |
| 8 | 0.0507 (ns) | 0.2240 (ns) |
| 9 | <0.0001 (****) | 0.0009 (***) |
| 10 | <0.0001 (****) | 0.0779 (ns) |
| 11 | <0.0001 (****) | >.9999 (ns) |
| 12 | <0.0001 (****) | 0.0285 (*) |
| 13 | <0.0001 (****) | <0.0001 (****) |
| 14 | <0.0001 (****) | <0.0001 (****) |
| 15 | <0.0001 (****) | <0.0001 (****) |
| 16 | <0.0001 (****) | <0.0001 (****) |
| 17 | <0.0001 (****) | <0.0001 (****) |
| 18 | – | 0.0006 (***) |
| 19 | – | 0.0356 (*) |
| 20 | – | 0.9615 (ns) |
| 21 | – | 0.2499 (ns) |

Table S5: Bonferroni-adjusted p-values for multiple comparison via Two-way ANOVA

| | Pup weight compared to PBS control | |
|---|---|---|
| Day | I.V. 5mg/kg Fc-PRL-13 (single dose) | S.C. 5mg/kg Fc-PRL-13 (single dose) |
| 7 | 0.2060 (ns) | 0.1398 (ns) |
| 8 | 0.0507 (ns) | 0.1146 (ns) |
| 9 | 0.0096 (**) | 0.0575 (ns) |
| 10 | 0.1632 (ns) | 0.1932 (ns) |
| 11 | 0.5620 (ns) | 0.2990 (ns) |
| 12 | 0.4491 (ns) | 0.5932 (ns) |
| 13 | 0.7302 (ns) | >0.9999 (ns) |
| 14 | >0.9999 (ns) | >0.9999 (ns) |
| 15 | >0.9999 (ns) | >0.9999 (ns) |
| 16 | >0.9999 (ns) | >0.9999 (ns) |
| 17 | >0.9999 (ns) | >0.9999 (ns) |
| 18 | >0.9999 (ns) | >0.9999 (ns) |
| 19 | 0.5366 (ns) | >0.9999 (ns) |
| 20 | 0.7386 (ns) | 0.7695 (ns) |
| 21 | >0.9999 (ns) | 0.8015 (ns) |

Table S6: Bonferroni-adjusted p-values for multiple comparison via Two-way ANOVA

| Day | Pup weight compared to Vehicle + PBS control | | | | |
|---|---|---|---|---|---|
| | BR + PBS | BR + 0.05mg/kg Fc-PRL-13 (repeat dose) | BR + 0.5mg/kg Fc-PRL-13 (repeat dose) | BR + 5mg/kg Fc-PRL-13 (repeat dose) | BR + 5mg/kg PRL N59D (repeat dose) |
| 7 | 0.0508 (ns) | 0.1751 (ns) | >0.9999 (ns) | 0.3461 (ns) | >0.9999 (ns) |
| 8 | 0.0845 (ns) | 0.9613 (ns) | 0.1071 (ns) | >0.9999 (ns) | 0.1998 (ns) |
| 9 | <0.0001 (****) | 0.0016 (**) | 0.8271 (ns) | >0.9999 (ns) | 0.0010 (**) |
| 10 | <0.0001 (****) | <0.0001 (****) | 0.0632 (ns) | >0.9999 (ns) | <0.0001 (****) |
| 11 | <0.0001 (****) | <0.0001 (****) | 0.0052 (**) | 0.1509 (ns) | <0.0001 (****) |
| 12 | <0.0001 (****) | <0.0001 (****) | <0.0001 (****) | 0.0013 (**) | <0.0001 (****) |
| 13 | <0.0001 (****) | <0.0001 (****) | <0.0001 (****) | 0.0005 (***) | <0.0001 (****) |
| 14 | <0.0001 (****) | <0.0001 (****) | <0.0001 (****) | <0.0001 (****) | <0.0001 (****) |
| 15 | <0.0001 (****) | 0.0048 (**) | <0.0001 (****) | <0.0001 (****) | <0.0001 (****) |
| 16 | <0.0001 (****) | <0.0001 (****) | <0.0001 (****) | <0.0001 (****) | <0.0001 (****) |
| 17 | <0.0001 (****) | 0.0048 (**) | <0.0001 (****) | <0.0001 (****) | <0.0001 (****) |
| 18 | - | <0.0001 (****) | <0.0001 (****) | <0.0001 (****) | 0.0145 (*) |
| 19 | - | - | <0.0001 (****) | 0.0638 (ns) | 0.0423 (*) |
| 20 | - | - | 0.0002 (***) | 0.1586 (ns) | 0.1831 ns) |
| 21 | - | - | <0.0001 (****) | 0.1067 (ns) | 0.0823 (ns) |

Table S7: Bonferroni-adjusted p-values for multiple unpaired t-tests

| Day | 5mg/kg Fc-PRL-13 (repeat dose) v. PBS control |
|---|---|
| 7 | >0.999999 (ns) |
| 8 | >0.999999 (ns) |
| 9 | >0.999999 (ns) |
| 10 | >0.999999 (ns) |
| 11 | >0.999999 (ns) |
| 12 | >0.999999 (ns) |
| 13 | >0.999999 (ns) |
| 14 | >0.999999 (ns) |
| 15 | >0.999999 (ns) |
| 16 | >0.999999 (ns) |
| 17 | >0.999999 (ns) |
| 18 | >0.999999 (ns) |
| 19 | 0.584205 (ns) |
| 20 | >0.999999 (ns) |
| 21 | >0.999999 (ns) |
