## Extended Data for "A long-acting prolactin to combat lactation insufficiency"

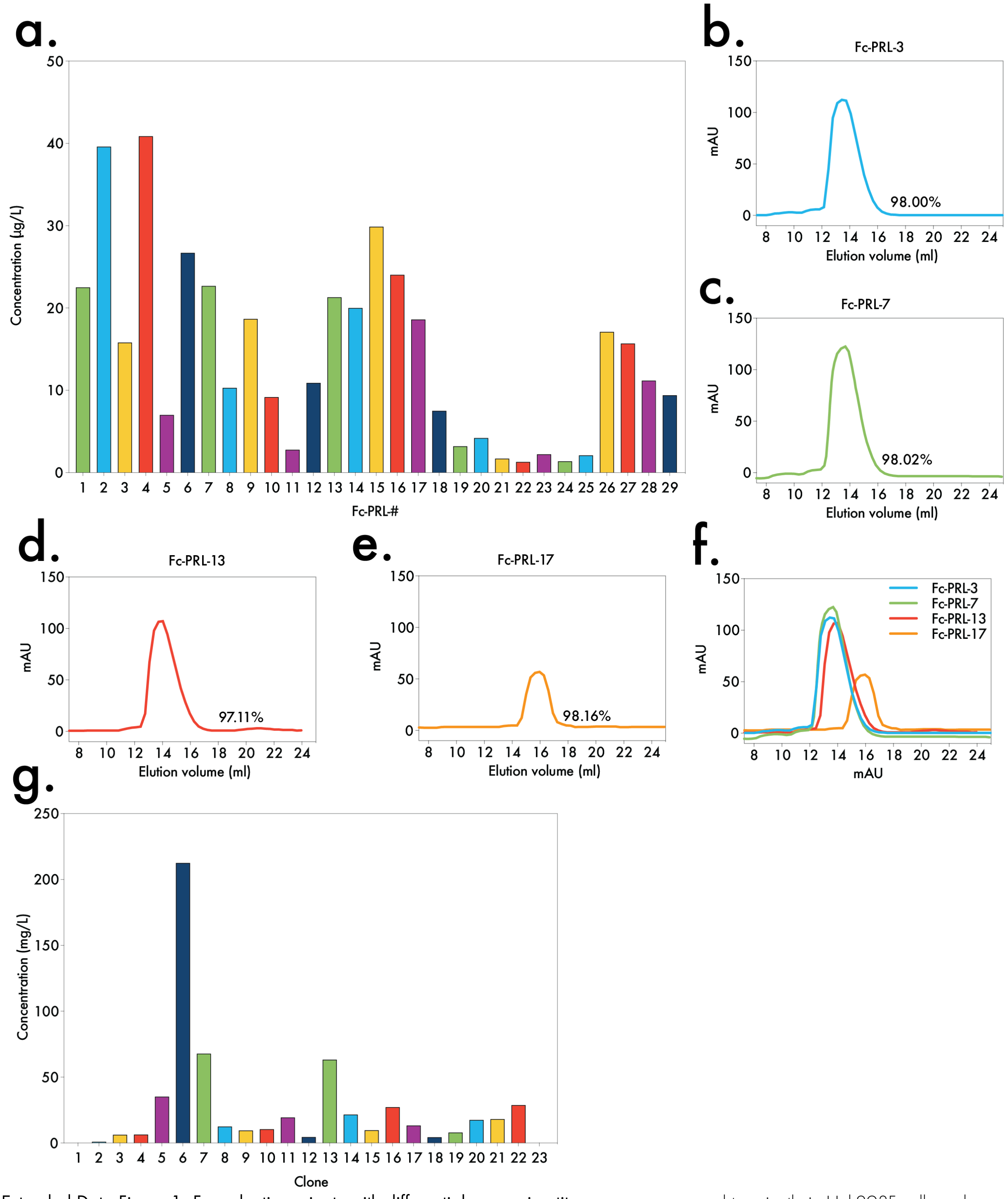


**Extended Data Figure 1: Fc-prolactin variants with differential expression titers** were **a** expressed transiently in Hek293F cells and purified either by Protein A (Fc-PRL- 1, 3, 5-7, 9-17, 19, 22-29) or His-Tag (Fc-PRL- 2, 4, 8, 18, 20-21). Molecular weights were verified by reducing SDS-PAGE gels and/or western blots (data not shown). Protein concentration was determined by BCA. **b-e** FPLC analysis of top 4 Fc-prolactin fusions using size-exclusion chromatography columns was performed to determine purity and aggregation. The SEC prolife (**b** Fc-PRL-3, **c** Fc-PRL-7, **d** Fc-PRL-13, **e** Fc-PRL-17, and **f** overlay) and the abundance (percentage) is presented for the different fusions. **c** P*ichia pastoris* clones of Fc-PRL-13 were grown in 3mL scout cultures of and expression titers were determined via ELISA. Clone 6 was identified as the highest expressing clone, and was used to produce protein for *in vivo* studies.

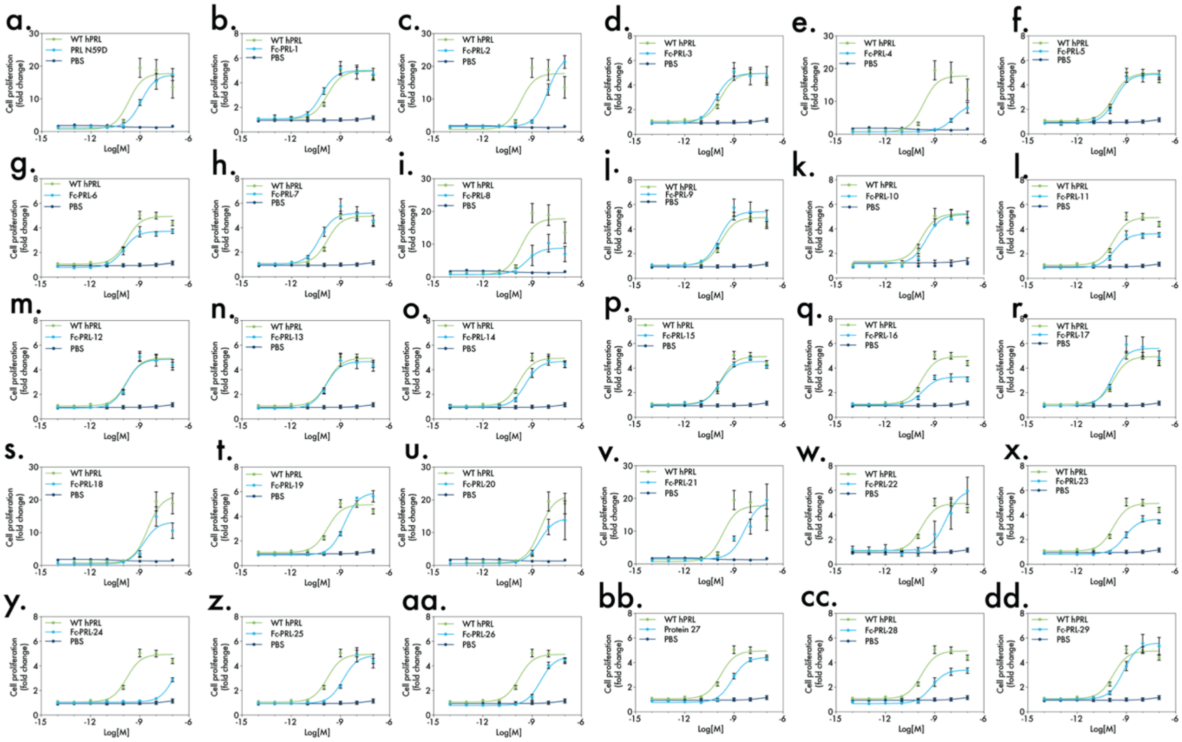


**Extended Data Figure 2: Dose response curves demonstrating bioactivity of Fc-prolactin variants via human PRLR signaling.** Fc-prolactin variants were assayed in an *in vitro* cell-based signaling assay previously described[79]. PRISM was used to fit a non-linear curve and calculate Log(EC50) and Emax (Extended data table 3). Data is depicted as mean ± SEM.

Extended Data Table 3:
Bioactivity of Fc-prolactin variants via human PRLR signaling

| Protein | N | Log (EC50) | % WT hPRL Emax |
|---|---|---|---|
| **PRL WT +** | 5 | -9.8 | 100% |
| **PRL WT *** | 16 | -9.7 | 100% |
| **PRL N59D +** | 3 | -8.9 | 95% |
| **Fc-PRL-1 *** | 3 | -10.2 | 101% |
| **Fc-PRL-2 +** | 3 | -8.0 | 127% |
| **Fc-PRL-3 *** | 3 | -10.1 | 104% |
| **Fc-PRL-4 +** | 3 | -7.9 | 49% |
| **Fc-PRL-5 *** | 3 | -9.7 | 103% |
| **Fc-PRL-6 *** | 3 | -10.0 | 75% |
| **Fc-PRL-7 *** | 3 | -10.3 | 107% |
| **Fc-PRL-8 +** | 3 | -9.4 | 47% |
| **Fc-PRL-9 *** | 3 | -9.9 | 113% |
| **Fc-PRL-10*** | 3 | -9.5 | 103% |
| **Fc-PRL-11 *** | 3 | -9.7 | 71% |
| **Fc-PRL-12 *** | 3 | -9.8 | 95% |
| **Fc-PRL-13 *** | 3 | -9.9 | 98% |
| **Fc-PRL-14 *** | 3 | -9.4 | 96% |
| **Fc-PRL-15 *** | 3 | -9.9 | 91% |
| **Fc-PRL-16*** | 2 | -9.7 | 58% |
| **Fc-PRL-17 *** | 3 | -9.8 | 120% |
| **Fc-PRL-18 +** | 3 | -8.8 | 75% |
| **Fc-PRL-19 *** | 1 | -8.8 | 131% |
| **Fc-PRL-20 +** | 3 | -8.5 | 81% |
| **Fc-PRL-21 +** | 3 | -8.4 | 103% |
| **Fc-PRL-22 *** | 2 | -8.3 | 126% |
| **Fc-PRL-23 *** | 3 | -9.1 | 73% |
| **Fc-PRL-24 *** | 3 | -6.9 | 109% |
| **Fc-PRL-25 *** | 3 | -8.8 | 100% |
| **Fc-PRL-26 *** | 3 | -8.5 | 102% |
| **Fc-PRL-27 *** | 3 | -9.1 | 94% |
| **Fc-PRL-28 *** | 3 | -9.2 | 70% |
| **Fc-PRL-29 *** | 3 | -9.2 | 118% |

* Readout by MTS reagent

+ Readout by WST-1 reagent

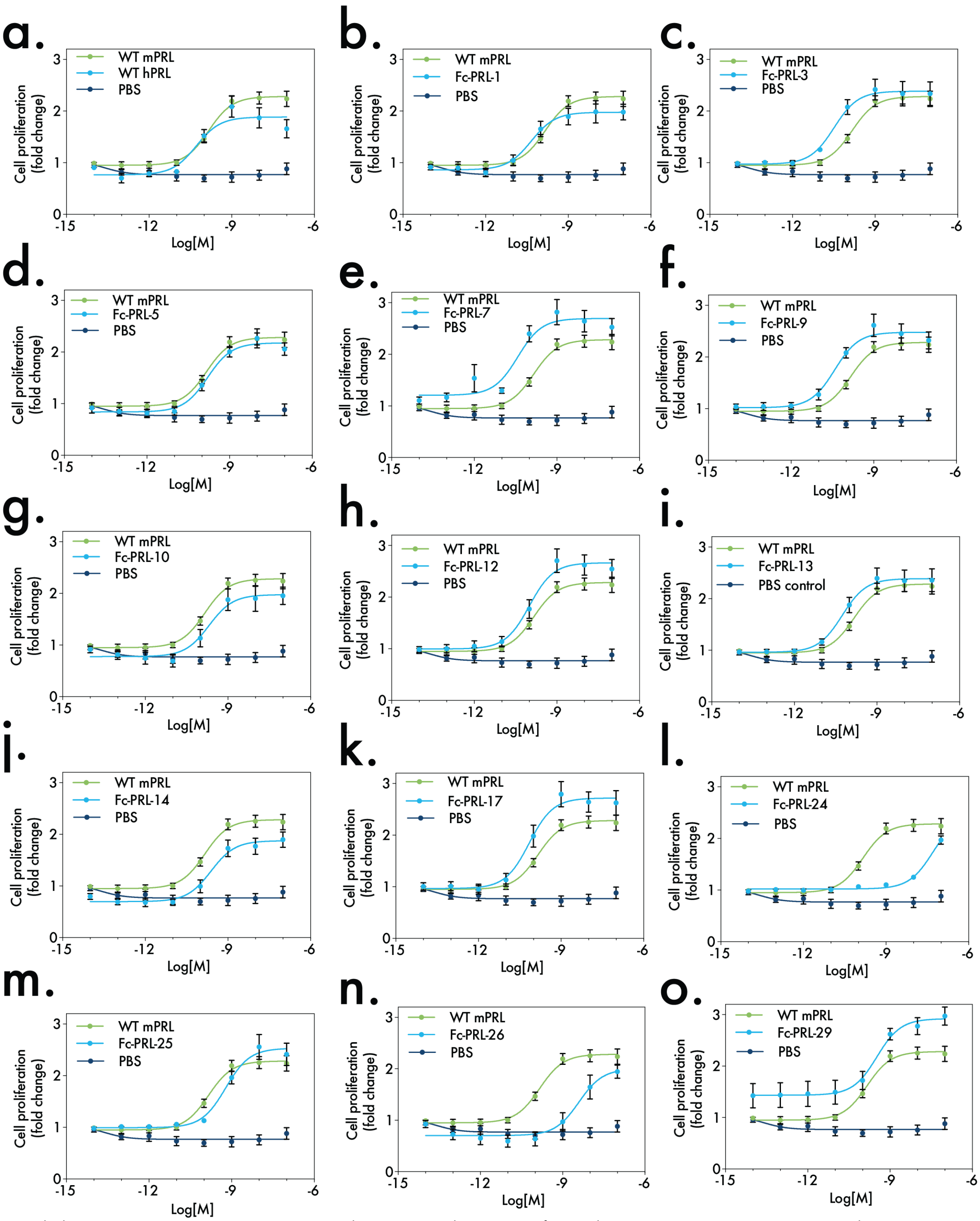


Extended Data Figure 4: Dose response curves demonstrating bioactivity of Fc-prolactin cariants via mouse PRLR signaling. Fc-prolactin variants were assayed in an *in vitro* cell-based signaling assay previously described[79]. PRISM was used to fit a non-linear curve and calculate Log(EC50) and Emax (Extended data table 5). Data is depicted as mean ± SEM.

Extended Data Table 5:
Bioactivity of Fc-prolactin variants via mouse PRLR signaling

| Protein | N | Log (EC50) | % WT mPRL Emax |
|---|---|---|---|
| mPRL WT | 6 | -9.8 | 100% |
| hPRL WT | 3 | -10.3 | 84% |
| Fc-PRL-1 | 3 | -10.3 | 84% |
| Fc-PRL-3 | 3 | -10.5 | 106% |
| Fc-PRL-5 | 3 | -9.8 | 100% |
| Fc-PRL-7 | 3 | -10.4 | 112% |
| Fc-PRL-9 | 3 | -10.4 | 109% |
| Fc-PRL-10 | 3 | -9.7 | 90% |
| Fc-PRL-12 | 3 | -10.0 | 125% |
| Fc-PRL-13 | 3 | -10.3 | 107% |
| Fc-PRL-14 | 3 | -9.6 | 88% |
| Fc-PRL-17 | 3 | -10.2 | 132% |
| Fc-PRL-24 | 3 | -7.3 | 107% |
| Fc-PRL-25 | 3 | -9.2 | 115% |
| Fc-PRL-26 | 3 | -8.4 | 97% |
| Fc-PRL-29 | 3 | -9.5 | 112% |

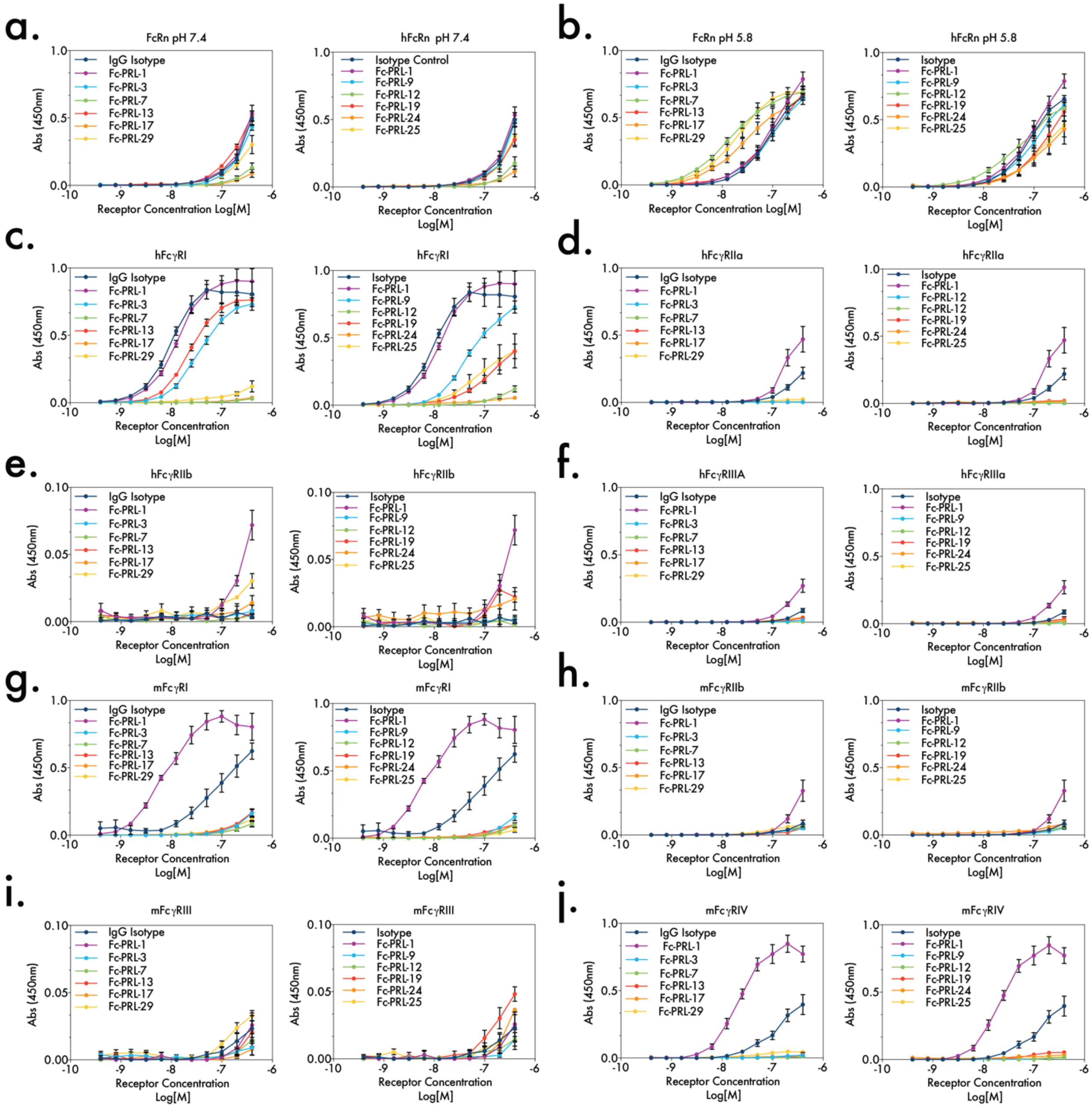


**Extended Data Figure 6: Binding affinity of human Fc-prolactin variants to human FcRn, human Fc receptors, or mouse Fc receptors by ELISA.** The affinity of the fusions to hFcRn (**a-b**) hFcɣRI (**c**), hFcɣRIIa (**d**), hFcɣRIIb (**e**), hFcɣRIIa (**f**), mFcɣRI (**g**), mFcɣRIIb (**h**), mFcɣRIII (**i**), and mFcɣRIV (**j**) was measured via ELISA. For positive controls, we used IgG isotype control and Protein 1, which is wildtype, glycosylated Fc fused to PRL (N59D). Data is depicted at mean ± SEM triplicates except Fc-PRL-24 (n=2 for human receptors and n=1 for mouse receptors) and Fc-PRL-29 (n=2 for human and mouse receptors).

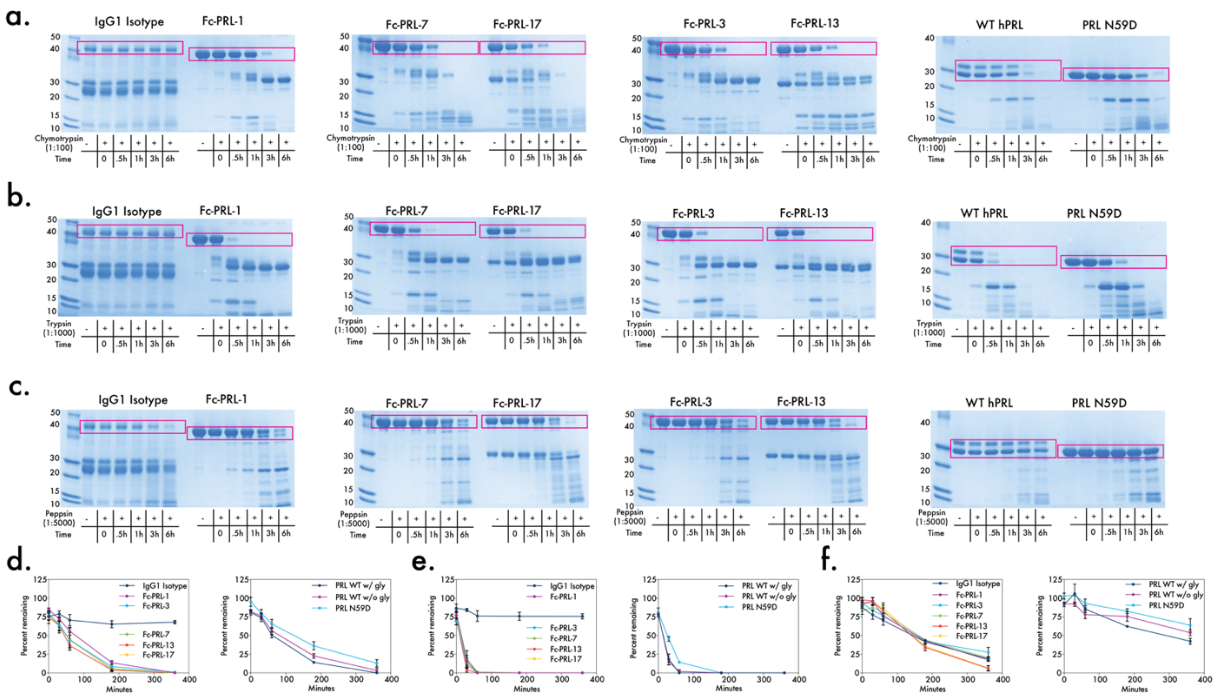


Figure S7: Gastric protease degradation of Fc-prolactin variants. Fc-prolactin variants were incubated with gastric proteases trypsin, chymotrypsin, and pepsin at 1:1000, 1:100, and 1:5000 and 37C, 25C, and 37C respectively. Aliquots were taken from the reaction at different time points, and the percent of the fusion remaining in each aliquot was measured by SDS-PAGE (a-c) and densitometry (d-f). For positive controls, we used IgG isotype control and Protein 1, which is wildtype, glycosylated Fc fused to PRL (N59D). Experiments were conducted in triplicate. The arrows indicate bands that were used to calculate the percent remaining for each fusion. Data is depicted as mean ± SEM of triplicates.

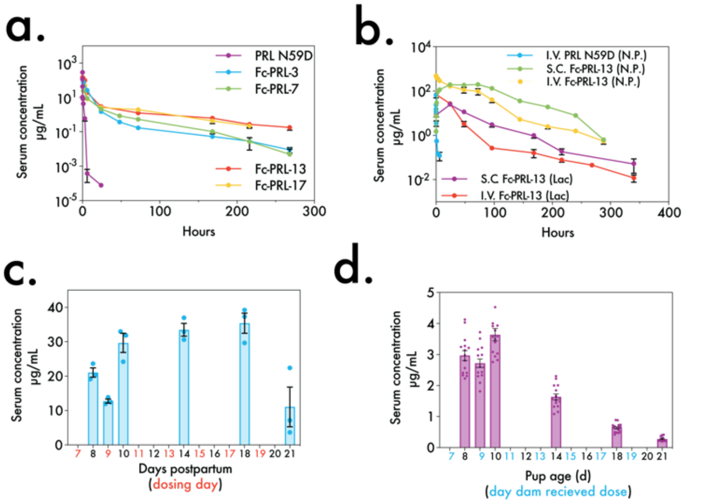


**e.** PK parameters of top 4 perfomring Fc-prolactin variants in Tg276 mice

| Protein | $AUC_{inf}$ (ug days/ mL) | Clearance (mL/ day) | β-phase $T_{1/2}$ (h) | $V_{SS}$ (mL/ Kg) |
|---|---|---|---|---|
| Tg276 mice | | | | |
| I.V. PRL N59D(N.P.) | 24 | 4.2 | 0.027* | 2.3 |
| I.V. Fc-PRL-3 (N.P.) | 649.1 | 0.15 | 49.3 | 1.2 |
| I.V. Fc-PRL-7 (N.P.) | 311.2 | 0.32 | 35.1 | 7.0 |
| I.V. Fc-PRL-13 (N.P.) | 2305 | 0.04 | 70.9 | 0.39 |
| I.V. Fc-PRL-17 (N.P.) | 1416 | 0.07 | 40.2 | 0.88 |

N.P. = nulliparous mice
* Data fit using one-phase decay, all other data was fit using two-phase decay

**f.** PK parameters of Fc-PRL-13 in C57bl/ 6j mice

| Protein | $AUC_{inf}$ (ug days/ mL) | Clearance (mL/ day) | β-phase $T_{1/2}$ (h) | $V_{SS}$ (mL/ Kg) |
|---|---|---|---|---|
| I.V. PRL N59D (NP) | 9.0 | 11.1 | 0.17* | 13.3 |
| I.V. Fc-PRL-13 (NP) | 15227 | 0.0066 | 26.8 | 0.27 |
| S.C. Fc-PRL-13 (NP) | 22708 | 0.0044 | 112+ | 0.32 |
| I.V. Fc-PRL-13 (Lac) | 1586 | 0.063 | 10.9* | 0.88 |
| S.C. Fc-PRL-13 (Lac) | 1369 | 0.0730 | 31.9+ | 1.13 |

NP = nulliparous mice
Lac = lactating mice
* Data fit using one-phase decay, all other data was fit using two-phase decay
+ A one-phase decay was used to fit the data after peak serum concentration

**Extended Data Figure 8: Pharmacokinetic profiles of Fc-PRL-13 in C57Bl/6j mice. a** Nulliparous Tg276 mice were injected with 5mg/kg I.V. of Fc-prolactin fusions (n=4) and PRL N59D (n=5). Blood was collected by tail nick post injection, and the concentration of the fusions in serum was measured by ELISA. The data are depicted as mean ± SEM. PRISM was used to fit either a one-phase decay (PRL N59D) or a two-phase decay (Fc-prolactin fusions). The Pharmacokinetic parameters are listed in the Table in **e. b** C57bl/6j mice were injected with I.V. or S.C. administered 5mg/kg of Fc-PRL-13 on the 7$^{th}$ day post-partum, and litters were normalized to n=5. Dose groups consist of nulliparous (N.P.) or lactating (Lac) mice (n=5 for I.V. PRL N59D (N.P.), n=8 for S.C. Fc-PRL-13 (N.P.), n=5 for I.V. Fc-PRL-13 (N.P.), n=5 for I.V. Fc-PRL-13 (Lac), and n=7 for S.C. Fc-PRL-13 (Lac)). Blood was collected by tail nick post injection, and the concentration of Fc-PRL-13 in serum was measured by ELISA. The data are depicted as mean ± SEM. PRISM was used to fit either a one-phase decay (PRL N59D or S.C. administered S.C.) or a two-phase decay (I.V. administered Fc-prolactin fusions). The relevant pharmacokinetic parameters are listed in **f. c** Lactating C57bl/6j mice were administered S.C. 5mg/kg of Fc-PRL-13 every other day. Mice were sacrificed at 6 different time points (n=3), and blood was collected by cardiac puncture. The concentration of Fc-PRL-13 in serum was measured by ELISA. The data are depicted as mean ± SEM. **d** The pups of the dams repeatedly dosed were also sacrificed (n=15), and their blood was collected by decapitation. The concentration of the fusions in serum was measured by ELISA. The data are depicted as mean ± SEM.

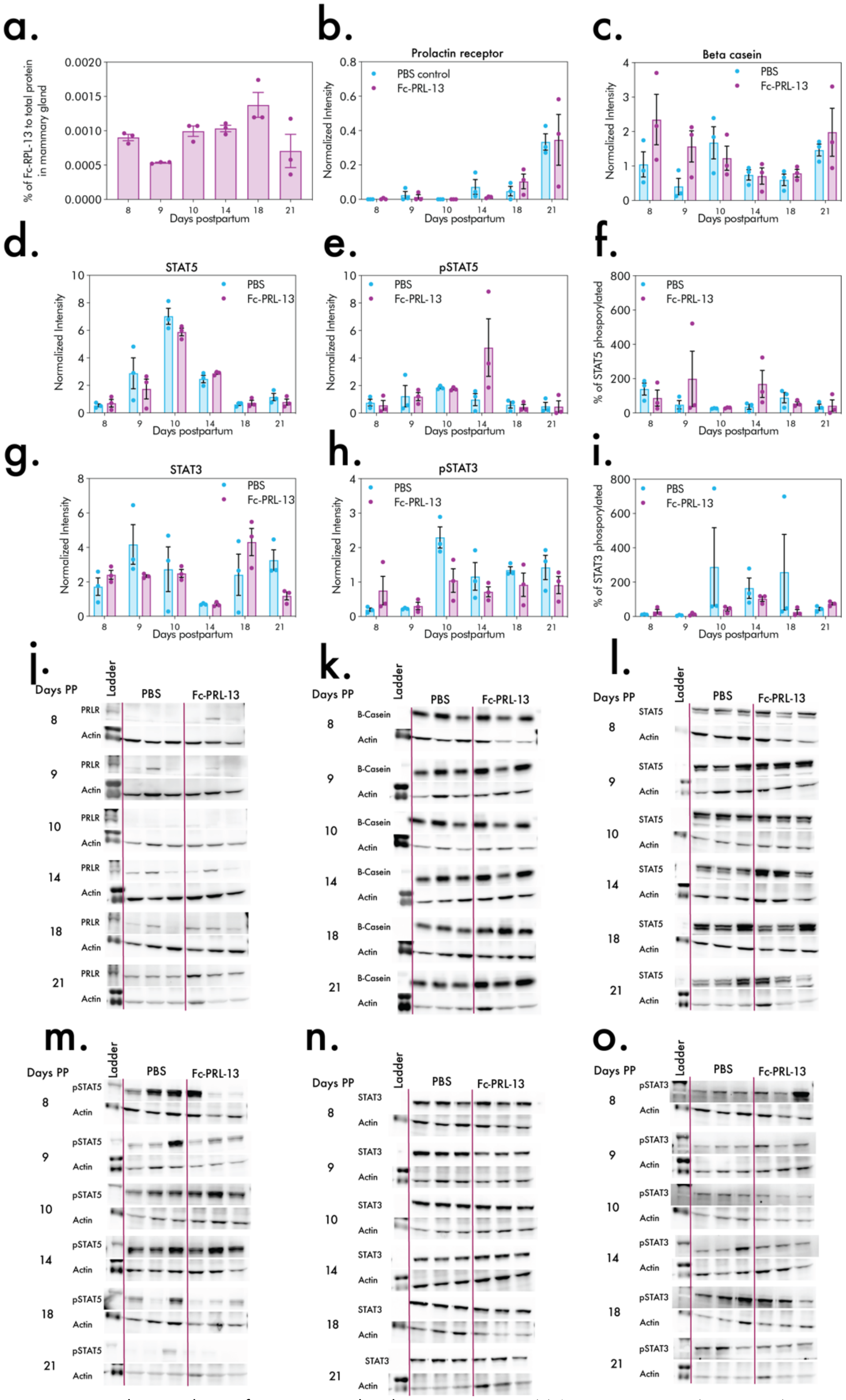

Extended Data Figure 9: Biomarker analysis of mammary glands. Lactating C57bl/6j mice were administered S.C. 5mg/kg of Fc-PRL-13 every other day. Mice were sacrificed at 6 different time points (n=3). Their abdominal mammary glands were collected, homogenized in RIPA buffer with protease and phosphatase inhibitors, and stored at -20C. **a** The concentration of Fc-PRL-13 in serum was measured by ELISA. The data are depicted as mean ± SEM. Expression of prolactin receptor (**b** and **j**), β-casein (**c** and **k**), STAT5 (**d** and **l**), pSTAT5 (**e** and **m**), STAT3 (**g** and **n**), and pSTAT3 (**h** and **o**) were analyzed by western blot and measured by densitometry. Intensity was normalized to an β-actin loading control. The data are depicted as mean ± SEM. The percent of phosphorylated STAT5 to total STAT5 and the percent of phosphorylated STAT3 to total STAT3 is depicted in **f** and **i**, respectively. Multiple unpaired t-tests with

Bonferroni correction for multiple comparison were used to calculate the statistical. All comparisons were not statistically significant (alpha = 0.05).

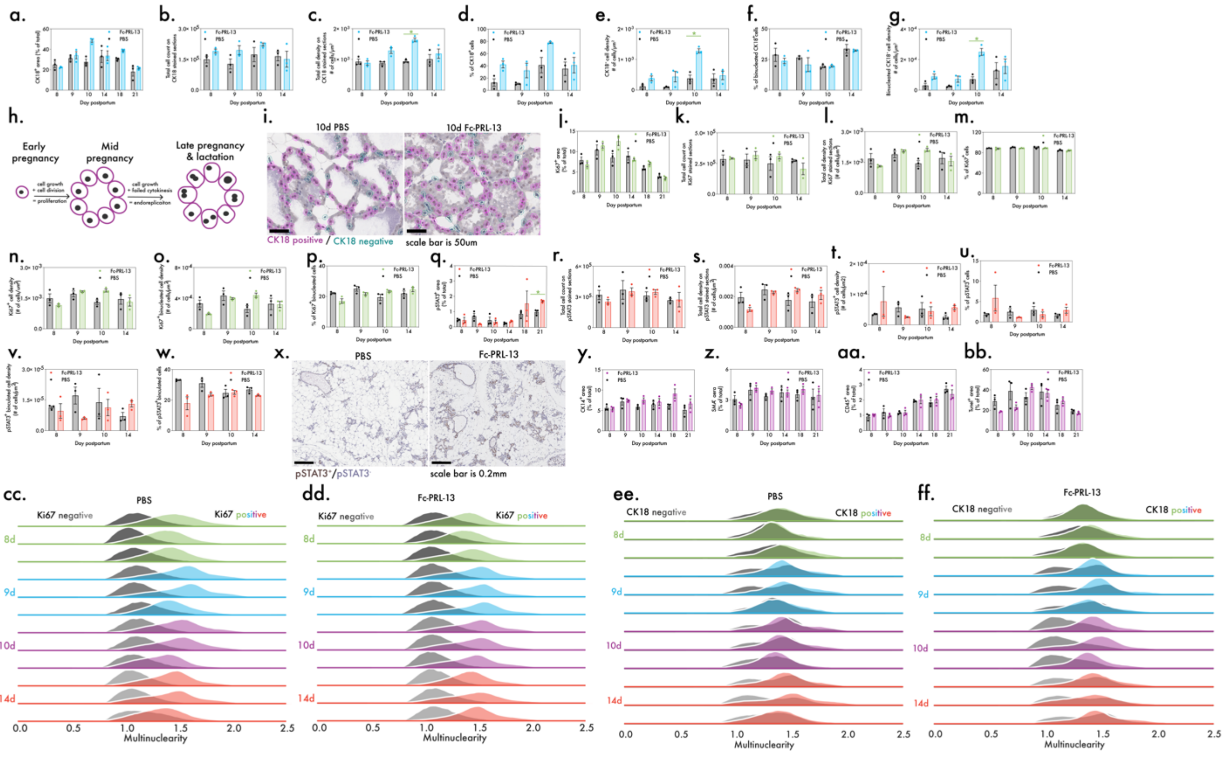


**Extended Data Figure 10: Immunohistochemistry analysis of mammary glands.** The following histological and biomarkers analyses are from a single experiment where B6 dams were dosed with subcutaneously administered Fc-PRL-13 (5mg/kg) or PBS every other day starting on the 7$^{th}$ day postpartum. Litters were normalized (n=5) on the 7$^{th}$ day postpartum. Mammary glands were collected from each mouse (time points= 8d, 9d, 10d, 14d, 18d, and 21d postpartum; n=3 mice per time point) for immunohistochemical staining of CK18(**a-g**), Ki67 (**i-o**), pSTAT3 (**q-w**), CK14 (**y**), smooth muscle actin (SMA) (**z**), CD45 (**aa**) and TUNEL (**bb**). Qupath was used to measure overall DAB staining, annotate the area of the mammary gland, and to perform cell segmentation. For each cell, Qupath measures a variety of morphology parameters including nuclear area, perimeter, and max calipers. Downstream analysis of the segmented cells was done in Python. For **a-g**, **i-w**, and **y-bb**, the data are depicted as mean ± SEM. Multiple unpaired t-tests with Bonferroni correction for multiple comparison were used to calculate the statistical. Significant comparisons are labeled (green *, alpha = 0.05), otherwise the comparisons were not significant. **h** A cartoon illustrating mammary gland growth during early to mid-pregnancy via cell proliferation of lactocytes. During late pregnancy and lactation, lactocytes become cuboidal and cease to proliferate. A large proportion of lactocytes grow in size without dividing via endoreplication. They become binucleated, polypoidal, hypersecretory milk producing cells. **i** Sections were stained for CK18 via IHC, and cells were segmented using Qupath (n=3 per time point). Representative images are depicted (scale bar = 50um). **x** Sections were stained for STAT3 via IHC. Representative images are depicted (scale bar = 0.2mm). The ratio of area to perimeter, hereby called multinuclearity, can tell us if the distribution of cells with more than 1-2 nuclei is increasing. **cc** and **ff** show the distributions of multinuclearity for Ki67$^+$ (green, blue, purple red) and Ki67$^-$ (grayscale) cells across the time series (n=3 per time point). **ee** and **ff** show the distributions of multinuclearity for CK18$^+$ (green, blue, purple red) and CK18$^-$ (grayscale) across the time series (n=3 per time point). Mononuclear cells have a multinuclearity closer to 1, whereas binucleated cells have a multinuclearity closer to 1.5.
